# Long-range promoter-enhancer contacts are conserved during evolution and contribute to gene expression robustness

**DOI:** 10.1101/2021.02.26.432473

**Authors:** Alexandre Laverré, Eric Tannier, Anamaria Necsulea

**Affiliations:** Univ Lyon, Université Claude Bernard Lyon 1, CNRS, Laboratoire de Biométrie et Biologie Evolutive, F-69100, Villeurbanne, France; Centre de recherche Inria de Lyon, 69603 Villeurbanne, France

**Author notes:** Corresponding authors: **Alexandre Laverré**, Laboratoire de Biométrie et Biologie Evolutive, CNRS, Université Claude Bernard - Lyon 1, Université de Lyon, 43 Bd. du 11 Novembre 1918, 69622 Villeurbanne CEDEX, France,; **Anamaria Necsulea**, Laboratoire de Biométrie et Biologie Evolutive, CNRS, Université Claude Bernard - Lyon 1, Université de Lyon, 43 Bd. du 11 Novembre 1918, 69622 Villeurbanne CEDEX, France.

**Keywords:** regulatory evolution, gene expression evolution, chromatin conformation capture, PCHi-C, genome rearrangements, *cis*-regulatory landscapes, large-scale genome evolution, Promoter Capture Hi-C

## Abstract

Gene expression is regulated through complex molecular interactions, involving *cis*-acting elements that can be situated far away from their target genes. Data on long-range contacts between promoters and regulatory elements is rapidly accumulating. However, it remains unclear how these regulatory relationships evolve and how they contribute to the establishment of robust gene expression profiles. Here, we address these questions by comparing genome-wide maps of promoter-centered chromatin contacts in mouse and human. We show that there is significant evolutionary conservation of *cis*-regulatory landscapes, indicating that selective pressures act to preserve not only regulatory element sequences but also their chromatin contacts with target genes. The extent of evolutionary conservation is remarkable for long-range promoter-enhancer contacts, illustrating how the structure of regulatory landscapes constrains large-scale genome evolution. We show that the evolution of *cis*-regulatory landscapes, measured in terms of distal element sequences, synteny or contacts with target genes, is significantly associated with gene expression evolution.

## Introduction

The evolution of gene expression and the evolution of regulatory mechanisms have attracted considerable attention ever since the proposal that phenotypic differences between species may be driven by changes in gene activity rather than by changes in gene products (King and Wilson, 1975). In the past decade, these two topics have been extensively scrutinized through comparative “omics” approaches (Khaitovich et al., 2005; Gilad et al., 2006; Brawand et al., 2011; Villar et al., 2015; Wong et al., 2015; Berthelot et al., 2018; Cardoso-Moreira et al., 2019). These studies showed that gene expression patterns are well conserved across species (Brawand et al., 2011; Cardoso-Moreira et al., 2019), while the sequences and the activities of regulatory elements (in particular those of expression enhancers) evolve rapidly (Villar et al., 2015; Berthelot et al., 2018). These paradoxical observations warrant further exploration, to better understand the determinants of gene expression robustness in the presence of rapidly evolving regulatory landscapes. However, so far few attempts have been made to directly connect the evolution of gene expression to the evolution of regulatory mechanisms, at a genome-wide scale (Berthelot et al., 2018; Wong et al., 2017). These studies proposed that the presence of complex regulatory landscapes, involving numerous expression enhancers with potentially redundant roles, is the key driver of the robustness of gene expression levels (Berthelot et al. 2018).

To date, identifying the regulatory elements that control each gene remains a challenging task. Transcription is regulated through complex interactions between *trans*-acting factors and *cis*-acting elements. Genes are typically associated with multiple *cis*-acting elements, which can refine their expression levels, control different expression domains, or confer robustness through partial redundancy (Spitz and Duboule, 2008; Kvon et al., 2021). Conversely, each regulatory element can influence multiple genes, either concomitantly or in a context-specific manner (Schoenfelder and Fraser, 2019). The complexity of *cis*-acting regulatory landscapes is now better perceived thanks to chromatin conformation capture techniques, which identify pairs of genomic segments found in physical proximity in the nucleus (Dekker et al., 2002; Schoenfelder et al., 2015; Zhao et al., 2006). These techniques revealed numerous long-range chromatin contacts between promoters and distal regulatory elements (de Laat and Duboule, 2013). These long-range interactions challenge a common assumption, namely that the targets of *cis*-regulatory elements are the neighboring genes in the genome, within a certain genomic distance (McLean et al., 2010; Villar et al., 2015; Wong et al., 2017; Berthelot et al., 2018; Danko et al., 2018; Dukler et al., 2020).

Here, we study the evolution of *cis*-regulatory interactions and the evolution of gene expression, using genome-wide, high-resolution chromatin contact data. We perform a comparative analysis of *cis*-regulatory landscapes using promoter-centered chromatin interaction maps for human and mouse (Choy et al., 2018; Comoglio et al., 2018; Freire-Pritchett et al., 2017; Javierre et al., 2016; Koohy et al., 2018; Mifsud et al., 2015; Novo et al., 2018; Pan et al., 2018; Rubin et al., 2017; Schoenfelder et al., 2015, 2018; Siersbæk et al., 2017). Through this study, we aim to better understand the constraints imposed by the three-dimensional structure of *cis*-regulatory landscapes on genome evolution and the consequences of regulatory landscape changes on gene expression evolution.

## Results

### Promoter Capture Hi-C data collection and construction of a simulated interaction dataset

To examine the evolution of *cis*-regulatory landscapes, we processed and analyzed Promoter Capture Hi-C (PCHi-C) data derived from 16 human cell types and 7 mouse cell types (Methods, Supplemental Table S1). The PCHi-C technique was designed to detect interactions between gene promoters and other genomic regions, with high sensitivity and spatial resolution (Schoenfelder et al., 2015; Mifsud et al., 2015). Briefly, promoter-containing restriction fragments are targeted using RNA baits and interactions are scored between pairs of restriction fragments, involving at least one baited fragment (Schoenfelder et al., 2015; Mifsud et al., 2015). All data were generated with the same experimental protocol, ensuring that restriction maps are identical across all samples within a species (Methods). The dataset included interactions for 19,389 baited fragments for human and 21,858 for mouse. We focused on intra-chromosome (*cis*) interactions occurring at a linear genomic distance comprised between 25 kb and 2 Mb, and involving a baited and an un-baited restriction fragment (Methods).

To evaluate the significance of the observations obtained with this data, we simulated interactions that reproduce the distribution of distances between baited fragments and contacted fragments, as well as the numbers of contacts *per* baited region, for each sample (Fig. 1A-C, Supplemental Fig. S1, Methods). The simulated interactions involve the same set of baited fragments and are constructed on the same restriction map as the PCHi-C data (Fig. 1A, Methods). We designed this simulated dataset to account for the effect of the genomic distance between promoters and contacted regions, which is traditionally the main criterion for inferring regulatory interactions, in addition to being the main factor driving the likelihood of observing a chromatin contact (Cairns et al., 2016). However, our simulations cannot reproduce other characteristics of the PCHi-C data, such as the number of contacts *per* un-baited genomic fragment (Supplemental Fig. S1) or the number of contacts *per* baited region across all samples (Supplemental Fig. S2). The simulated interaction data cover a larger fraction of the genome than the PCHi-C data, for which contacted regions are more often shared among baits and among samples (Supplemental Fig. S2, Supplemental Text). Moreover, restriction fragments contacted in PCHi-C data may differ from those included in the simulations in terms of sequence uniqueness or mappability, because interactions can only be detected if sequencing reads can be correctly aligned to restriction fragments. To minimize this possible source of discrepancy between the PCHi-C and the simulated data, we filtered interactions to keep only those involving restriction fragments with a sufficient theoretical mappable fraction and PCHi-C read coverage (Methods, Supplemental Text).

**Figure 1.**
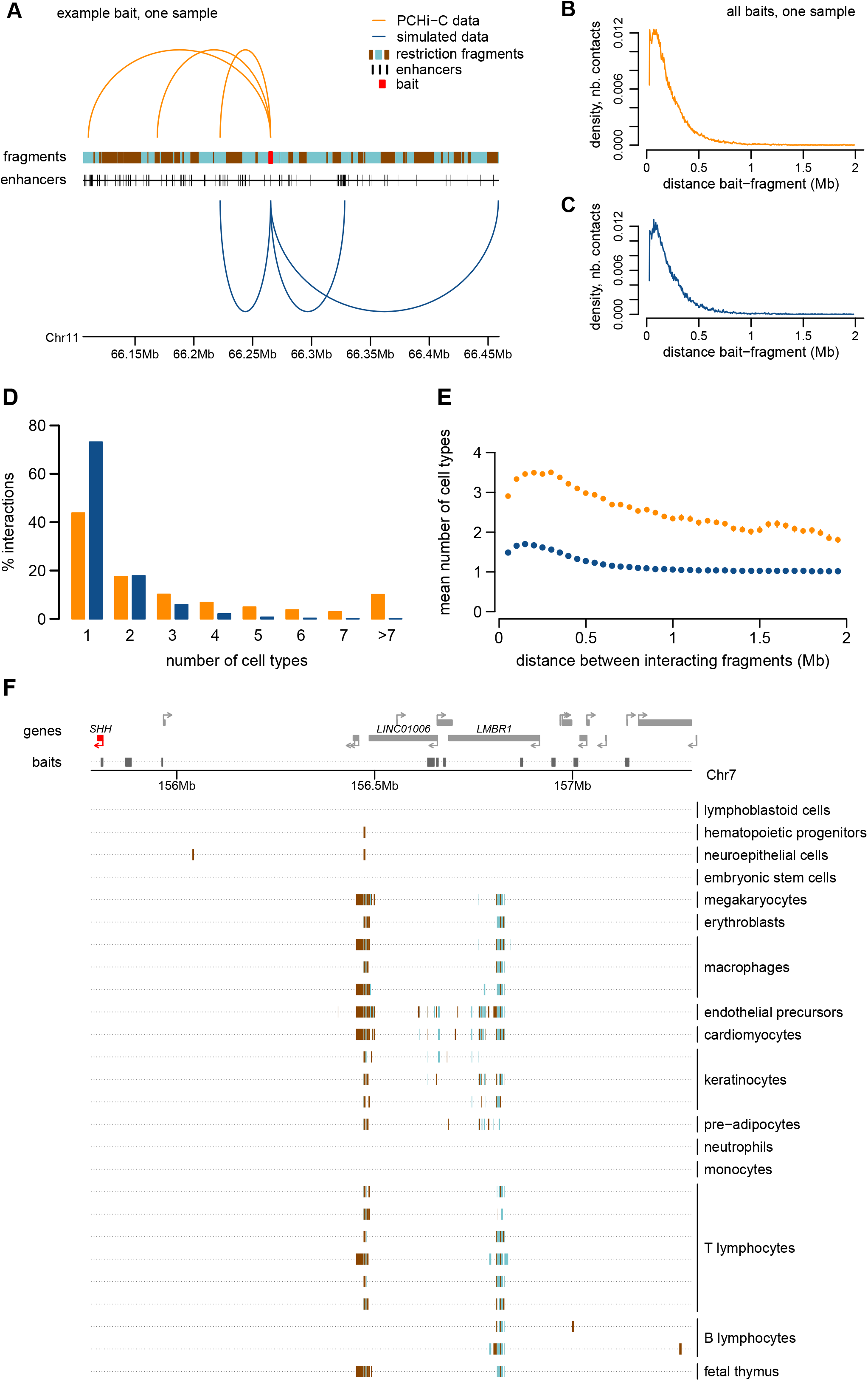
Chromatin interactions measured by PCHi-C data are shared among cell types. **A**. Example of interactions between a baited restriction fragment (red) and other restriction fragments, for PCHi-C data (orange) and simulated data (blue). The positions of ENCODE enhancers are displayed below the restriction fragments track. **B**. Distribution of genomic distances between baited fragments and contacted restriction fragments, in PCHi-C data (CD34 sample, human). **C**. Same as **B**, for simulated data. **D**. Histogram of the number of cell types in which interactions are observed, for human PCHi-C data and simulated data. **E**. Average number of cell types in which interactions are observed, as a function of the distance between baits and contacted fragments. Dots represent mean values, vertical segments represent 95% confidence intervals of the mean, obtained with a non-parametric bootstrap approach (Methods). **F**. Chromatin contacts between the SHH gene promoter and other genomic regions. From top to bottom: gene coordinates; localization of PCHi-C baited fragments; localization of restriction fragments contacted by the SHH bait in different samples. Rectangles with alternating colors indicate individual restriction fragments that are contacted by the *SHH* bait.

### Promoter-centered chromatin contacts can occur in the absence of gene expression

We first verified to what extent chromatin interactions are shared among cell types. Within each species, chromatin contacts cluster by cell type (Supplemental Fig. S3). This result is reassuring, given the inevitable batch effects that arise from combining PCHi-C data from several publications. Overall, 56% of the analyzed PCHi-C interactions (N=661,097) are detected in at least 2 cell types in human, compared to only 27% in the corresponding simulated data (N=1,828,577) (Fig. 1D, chi-squared test p-value < 10^−10^). The increase compared to simulated data is stronger for interactions that occur at large genomic distances. For example, at approximately 500 kb, human PCHi-C interactions are observed on average in 3 cell types, compared to only 1 cell type in the simulated data (Fig. 1E). We obtained similar results for mouse (Supplemental Text). We wanted to verify that this observation is not simply due to the presence of ubiquitously expressed genes, which may contact similar sets of regulatory elements in all cell types. We used published gene expression data from multiple organs and developmental stages (Cardoso-Moreira et al., 2019). The number of cell types in which chromatin contacts are observed is positively correlated with expression breadth, defined as the number of samples in which gene expression is detected (RPKM>=1, Methods, Supplemental Fig. S4). However, gene expression breadth spans a wide range of values (from 10% to 100% of samples) even for those genes that have contacts in all cell types (Supplemental Fig. S4). This is illustrated by the interaction landscape of the developmental gene *SHH*, which contacts regions situated in the introns of the neighboring *LMBR1* gene in almost all cell types (where its main enhancer, ZRS, is known to reside (Lettice et al., 2003)), even where *SHH* expression is not detected (Fig. 1F, Supplemental Fig. S5). This result confirms previous reports that promoter-centered interactions can be observed in PCHi-C data even in the absence of gene expression (Schoenfelder et al., 2015).

Thus, PCHi-C data provide a broad overview of promoter-centered chromatin interactions, likely including pre-formed contacts, which precede gene activation (de Laat and Duboule, 2013). These data thus extend beyond regulatory interactions that function exclusively in the sampled cell types. For interactions shared across cell types, differences between the human and mouse PCHi-C datasets are genuine between-species differences, rather than consequences of unequal cell type sampling. Thus, although similar cell types were not always available for human and mouse, we are confident that this dataset is suited for between-species comparisons.

### Inference of *cis*-regulatory landscapes from PCHi-C chromatin contact maps

The promoter-centered chromatin contacts defined with PCHi-C data are known to be enriched in regulatory interactions (Schoenfelder et al., 2015). We validated this observation by jointly analyzing PCHi-C contact maps and genome-wide enhancer prediction datasets (Methods). For all datasets, the average restriction fragment length covered by predicted enhancers is significantly higher in the PCHi-C data than in the simulated data (Fig. 2A, Wilcoxon rank sum test, p-value <10^−10^). For example, for ENCODE data (N=408,738 enhancers), the average length fraction that is covered by enhancers is 3.5% in the PCHi-C data, compared to 2.4% in the simulated data. Moreover, the proportion of restriction fragments that overlap with at least one enhancer is significantly higher in PCHi-C data than in simulations. For ENCODE, these proportions are 36% and 27% in PCHi-C data and in simulations, respectively (chi-squared test, p-value <10^−10^). The overlap with enhancers decreases when the distance between contacting regions increases, for both PCHi-C and simulated data (Fig. 2B). The number of cell types in which interactions are observed is also positively correlated with the presence of enhancers (Fig. 2C). Similar results were obtained for both species and for all four enhancer datasets (Supplemental Text). Hereafter, we consider that promoters and enhancers are in contact if the corresponding baited fragments of the promoters are in contact with restriction fragments that overlap with the corresponding enhancers, for both PCHi-C and simulated data (Methods).

**Figure 2.**
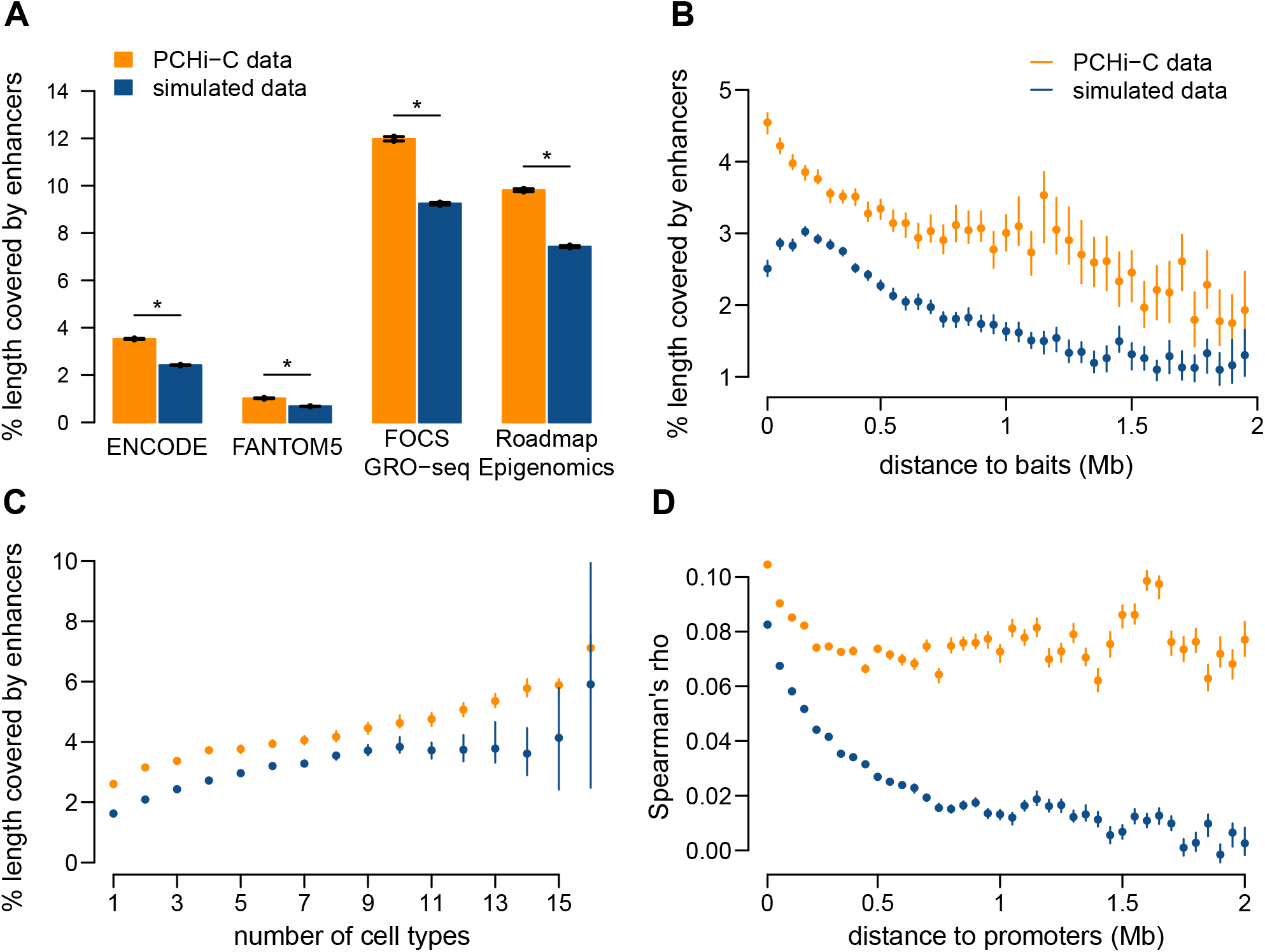
Gene-enhancer pairs connected by PCHi-C data are enriched in genuine regulatory interactions. **A**. Average length fraction covered by predicted enhancers, for restriction fragments contacted in human PCHi-C data (orange) and simulated data (blue). **B**. Average length fraction covered by ENCODE enhancers, as a function of the distance between baits and contacted restriction fragments. **C**. Average length fraction covered by ENCODE enhancers, as a function of the number of cell types in which interactions are observed, for human restriction fragments. **D**. Distribution of Spearman’s correlation coefficient between promoter and enhancer activity levels, for promoter-enhancer pairs in contact in PCHi-C data or in simulated data, according to the distance between them (Methods). **A-D**. Bars and dots represent mean values, vertical segments represent 95% confidence intervals of the mean, obtained with a non-parametric bootstrap approach (Methods). “*” indicates a significant difference between PCHi-C and simulated data (FDR <10^−10^) based on a chi-squared test.

We note that the numbers of enhancers assigned to genes based on PCHi-C data are considerably higher than those obtained with a classical genomic proximity approach where enhancers are assigned to the neighboring genes within a 25 kb - 2 Mb distance interval (Methods). For example, for the human ENCODE dataset, we find that 323,995 gene-enhancer pairs are in contact in PCHi-C data, while the genomic proximity approach predicts 224,446 pairs, for the 9,395 genes with enhancers assigned to them by both methods. The median gene contacts 25 enhancers, but has only 13 neighbor enhancers (Supplemental Fig. S6). Moreover, the PCHi-C data predicts regulatory interactions at larger genomic distances (median 277 kb for human ENCODE) than those predicted with the genomic proximity approach (median 88 kb). Only 42,095 gene-enhancer regulatory pairs were predicted by both approaches; the proportion of gene-enhancer pairs in common increases with the genomic distance between the two (Supplemental Fig. S6). As a general rule, we can thus confirm that contacts do not generally form between immediately neighboring promoters and enhancers (Smemo et al., 2014).

To further test the presence of genuine regulatory interactions in the PCHi-C data, we evaluated the correlations between gene expression and enhancer activity across samples (Methods). Due to the complexity of gene regulatory mechanisms, as well as due to inherent noise in activity measurements, these correlations are expected to be weak (Hait et al., 2018). Nevertheless, the activity levels of genes and enhancers connected in PCHi-C data are significantly better correlated than in simulated data (mean Spearman’s correlation coefficient 0.08 in the PCHi-C dataset and 0.04 in the simulations, Wilcoxon rank sum test p-value <10^−10^). The correlations between promoter and enhancer activity levels decrease when the distance between the two elements increases (Fig. 2D). This occurs for both PCHi-C and simulated data, although correlations always remain higher for the PCHi-C dataset. The decrease with the distance could be explained by a higher proportion of genuine regulatory relationships at a short distance from the promoter. However, it could also be explained by the influence of the chromatin environment on regulatory elements found in the vicinity of the promoter: transcriptional activation of the gene could lead to the establishment of open chromatin in the broader region around the promoter, leading to enhancer activation.

### Genomic sequences contacted by promoters are conserved during evolution

We analyzed the sequence conservation of restriction fragments and enhancers contacted by baited promoters by calculating the aligned sequence length in pairwise comparisons with nine other vertebrate genomes (Methods, Fig. 3A). We also used the phyloP score (Pollard et al., 2010) to investigate evolutionary conservation at broader evolutionary scales. We masked exonic regions for both approaches (Methods).

**Figure 3.**
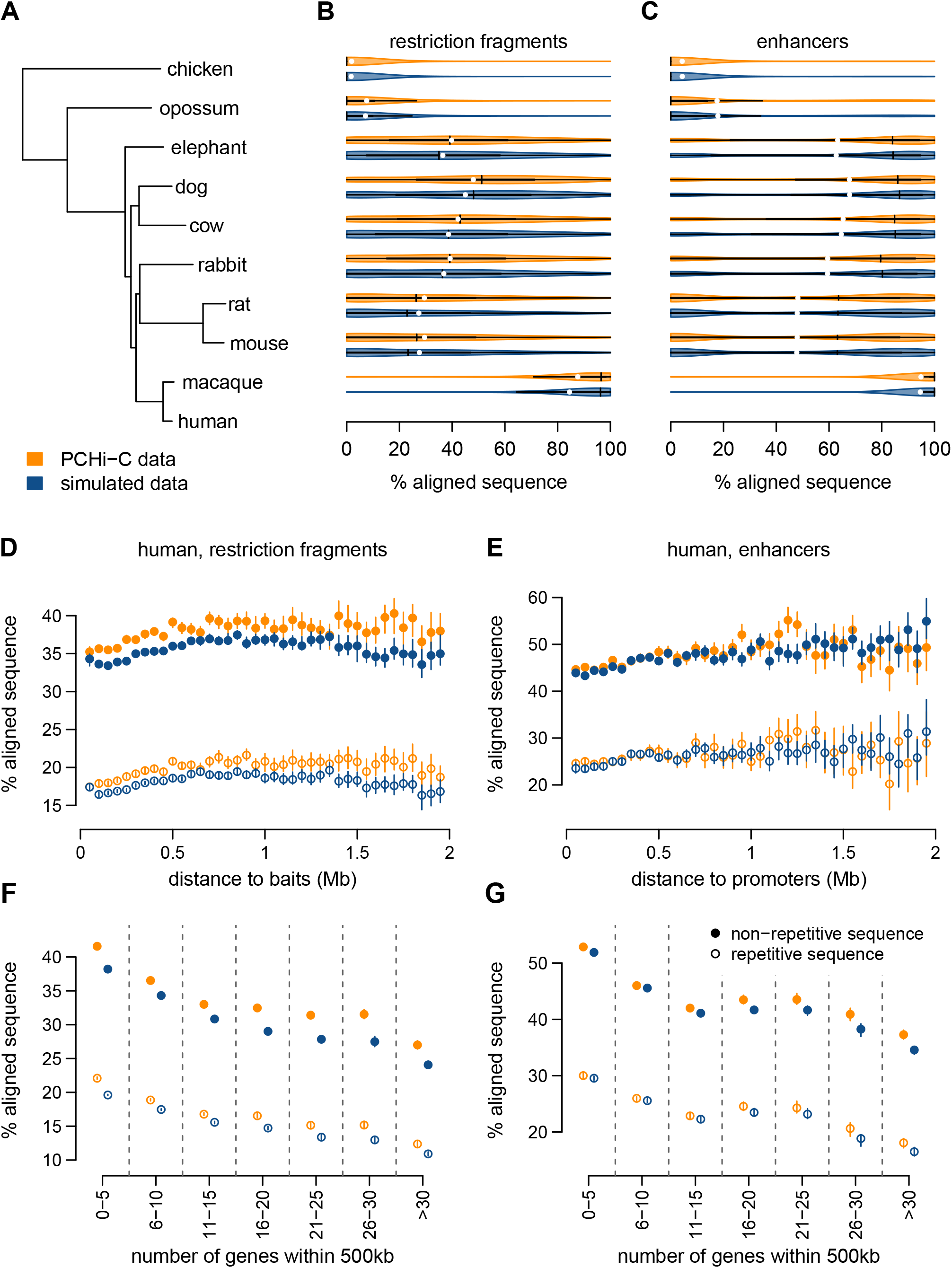
Sequences contacted by promoters are conserved during evolution. **A**. Phylogenetic tree for the analyzed species. **B**. Sequence conservation levels, derived from pairwise alignments, for contacted restriction fragments. Violin plots represent the distribution of the percentage of aligned nucleotides between human and other species. Vertical segments represent median values; white dots represent mean values. **C**. Sequence conservation levels for contacted ENCODE enhancers. **D**. Conservation of contacted restriction fragments between human and mouse, as a function of the median genomic distance between restriction fragments and contacting baits. **E**. Same as **D**, for ENCODE enhancers. **F**. Conservation of contacted restriction fragments between human and mouse, as a function of the number of genes found within at most 500kb from the restriction fragment. **G**. Same as **F**, for ENCODE enhancers. **D-G**. Dots represent mean values, vertical segments represent 95% confidence intervals of the mean, obtained with a non-parametric bootstrap approach (Methods). Filled dots represent non-repetitive sequences; empty dots represent repetitive sequences.

Our analyses reveal that restriction fragments contacted by baits in PCHi-C data are significantly more conserved than those included in simulated data (Fig. 3B, Supplemental Fig. S7, Supplemental Text). For the comparison between human and mouse, the median aligned length fraction of contacted fragments is 26.57% in PCHi-C data (median over N=263,176 fragments), which is significantly higher than the 23.25% observed in the simulated dataset (median over N=563,651 fragments) (Wilcoxon rank sum test, false discovery rate (FDR) <10^−10^). This observation holds for all pairwise comparisons between vertebrates, although conservation scores are expectedly weak for comparisons between divergent species (Fig. 3B, Supplemental Text).

In contrast, enhancers contacted by promoters in the PCHi-C dataset (N=170,306) are not significantly more conserved than enhancers included in the simulated dataset (N=292,599) (e.g., median aligned fraction 63.21% for PCHi-C data, 63.16% for simulated data, for human and mouse, Wilcoxon rank sum test, FDR 0.051, Fig. 3C, Supplemental Text). This suggests that the higher evolutionary conservation of PCHi-C contacted fragments compared to simulated data may be explained by overlap with more enhancers, but not by overlap with better conserved enhancers. We note that enhancers that overlap with restriction fragments in the simulated dataset but not in the PCHi-C data may be missing from the latter because they regulate other genes or function in other cell types.

The extent of sequence conservation tends to increase with the distance from gene promoters, for contacted restriction fragments as well as for enhancers (Supplemental Fig. S7, Supplemental Text). The distance between promoters and contacted regions or enhancers also co-varies with other factors that correlate with sequence conservation, such as the GC content, the overlap with repeats and the gene density in the neighboring regions (Supplemental Fig. S8). Regions included in the PCHi-C data and in the simulated data also often differ with respect to these genomic characteristics. In particular, restriction fragments contacted in PCHi-C data have lower proportions of repetitive sequences than those present in the simulated data (Supplemental Fig. S8). This observation holds true when separating regions in classes of similar theoretical mappability (Supplemental Fig. S9). We thus wanted to verify whether these genomic correlates could explain the sequence conservation patterns observed in PCHi-C and in simulated data. We first evaluated sequence conservation scores separately for repetitive and non-repetitive positions (Methods). For both classes of positions, conservation levels are higher for restriction fragments involved in PCHi-C contacts than for restriction fragments included in the simulations (Fig. 3D, Supplemental Fig. S10). The increase in conservation with the genomic distance in PCHi-C is more subtle when analyzing repetitive and non-repetitive sequences separately, indicating that it largely stems from the decrease in repeat proportion with the genomic distance (Fig. 3D,E, Supplemental Fig. S10).

Sequence conservation levels for restriction fragments and enhancers are negatively associated with the gene density in the neighboring regions, for both evolutionary conservation measures (Fig. 3F,G, Supplemental Fig. S10). This observation is true for both repetitive and non-repetitive sequences (Fig. 3F,G, Supplemental Fig. S10), indicating that it is not simply a consequence of the association between repeat frequency and gene density (Supplemental Fig. S11). For both repetitive and non-repetitive sequences, and irrespective of the gene density class, restriction fragments that are contacted in PCHi-C data are significantly more conserved than those included in the simulated data (Fig. 3F,G, Supplemental Fig. S10). We also observed higher sequence conservation in observed versus simulated PCHi-C data when we divided restriction fragments into classes of similar GC content (Supplemental Fig. S12), to account for the strong correlation between sequence composition and the rate of evolutionary divergence (Duret and Arndt, 2008).

These results illustrate the complexity of the factors affecting the evolution of *cis*-regulatory elements. The proportion of repeated elements, gene density, GC content and the distance to the baits are all highly correlated with measures of sequence conservation (Supplemental Table S2). The higher fraction of overlap with enhancers observed in PCHi-C data compared to simulated data may explain part of the difference in sequence conservation. The low frequency of repetitive elements observed in PCHi-C data may in itself be an indication of purifying selection acting on regulatory elements. Indeed, the most repeat-poor regions in the human and mouse genomes are the HOX gene clusters, which are crucial for embryonic development (International Human Genome Sequencing Consortium, 2001). Likewise, the negative association between sequence conservation levels and gene density may be partly explained by the presence of functionally constrained “gene deserts”, rich in regulatory elements (Ovcharenko et al., 2005). Consistent with these observations, genes that are in contact with highly conserved enhancers are enriched in functional categories related to development (Supplemental Table S3). Moreover, highly constrained human genes, as measured by the probability of intolerance to loss-of-function mutations (Lek et al., 2016, Methods) tend to contact more conserved enhancers (Supplemental Fig. S13), as previously proposed (Dukler et al., 2020).

### Pairs of promoters and enhancers involved in chromatin contacts are maintained in synteny

Genomic rearrangements that separate promoter-enhancer pairs to different chromosomes or to contact-prohibiting distances are expected to be counter-selected. To test this hypothesis, we assessed the proportion of promoter-enhancer pairs that are maintained in synteny (on the same chromosome and within a maximum distance of 2Mb) through pairwise comparisons between human/mouse and other vertebrate species. We restricted this analysis to promoter-enhancer pairs that are separated by distances between 100 kb and 1.5 Mb in the reference species (Methods). With this convention, synteny “breaks” are evolutionary events that add at least 500 kb to the distance between promoters and enhancers. We found that pairs of contacting promoters-enhancers are maintained in synteny significantly more often than in the simulated dataset (Fig. 4A, Supplemental Text). For example, 95.9% of pairs between promoters and ENCODE enhancers (N=207,144) are maintained in synteny between human and mouse, compared to 94.7% in the simulated dataset (N=496,622) (Fig. 4A, chi-squared test FDR < 10^−10^). At larger evolutionary distances, there is less difference between the PCHi-C data and the simulated interactions (Fig. 4A, chi-squared test FDR = 0.034 for the comparison between human and chicken). Our synteny conservation measure is influenced by the genome assembly quality, which likely explains the lower values observed for the comparison between human and rabbit (Fig. 4A). The excess of synteny conservation compared to simulated data is mainly visible at large genomic distances, which are likely to accumulate genomic rearrangements with time if these are not counter-selected (Fig. 4B,C, Supplemental Text).

**Figure 4.**
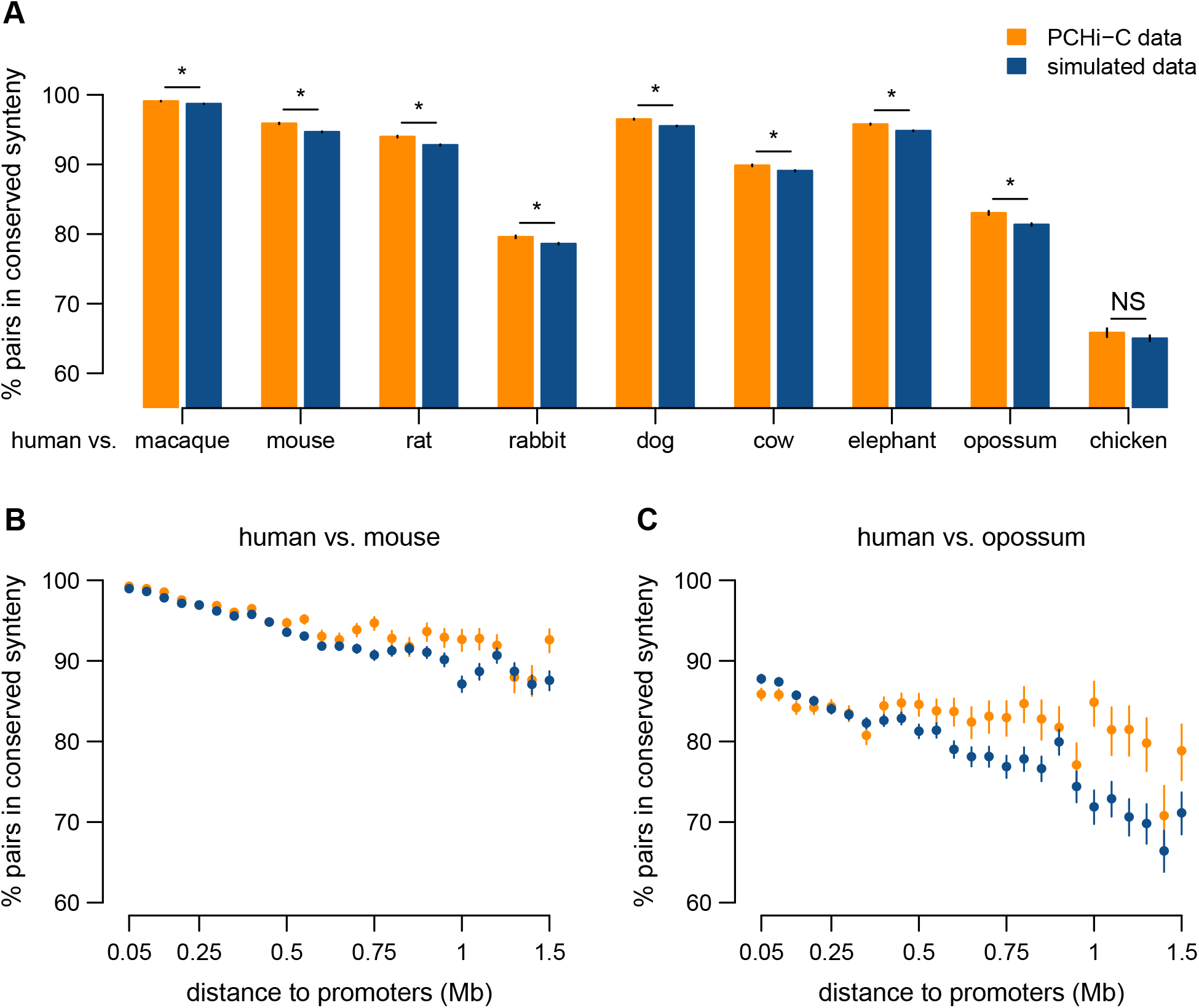
Long-range promoter-enhancer pairs are conserved in synteny. **A**. Proportion of human promoter-enhancer (ENCODE) pairs maintained in synteny (*i*.*e*., found on the same chromosome, within a maximum distance of 2 Mb) in other vertebrate genomes, for PCHi-C data (orange) and simulated data (blue). **B**. Proportion of human promoter-enhancer pairs maintained in synteny in the mouse genome, as a function of the distance between them in the human genome. **C**. Same as **B**, for the comparison between human and opossum. **B-C**. Bars represent 95% two-sided confidence intervals for the proportions (Methods). “*” indicates a significant difference between PCHi-C and simulated data (FDR <10^−10^) based on a chi-squared test.

### Promoter-enhancer contact maps are conserved during evolution

To further test the presence of selective pressures to maintain chromatin contacts between promoters and enhancers, we directly compared PCHi-C interaction landscapes between human and mouse. We tested for contact conservation between pairs of cell types in human and mouse (Methods). For this analysis, we selected promoter-enhancer pairs that are maintained in synteny in the target species, to avoid the confounding effect of genomic rearrangements that break contacts. We also drew the same number of interactions for each sample, to avoid the apparent excess of contact conservation in comparisons involving data with better sequencing depth (Methods, Supplemental Text).

For all cell type pairs, the frequency of conserved contacts is higher in the PCHi-C data (median=12.6%) than in the simulated data (median=0.99%, Wilcoxon rank sum test p-value <10^−10^; Fig. 5B). The extent of contact conservation is high for comparable cell types (embryonic stem cells or epiblast-derived stem cells, pre-adipocytes and B-lymphocytes), but higher values could be observed in comparisons involving different cell types (Fig. 5A). This could be explained by technical artifacts leading to better detection sensitivity in some samples, despite our subsampling procedure. Our chromatin contact conservation measures are necessarily under-estimates, given that human and mouse PCHi-C datasets do not consist of the same cell types and that cell-type specific interactions may thus appear as species-specific. Consistent with this, the extent of contact conservation is higher for interactions observed in multiple cell types (Fig. 5C, Supplemental Text). Furthermore, the extent of contact conservation increases with the score attributed to interactions by the CHiCAGO processing pipeline (Supplemental Fig. S14).

**Figure 5.**
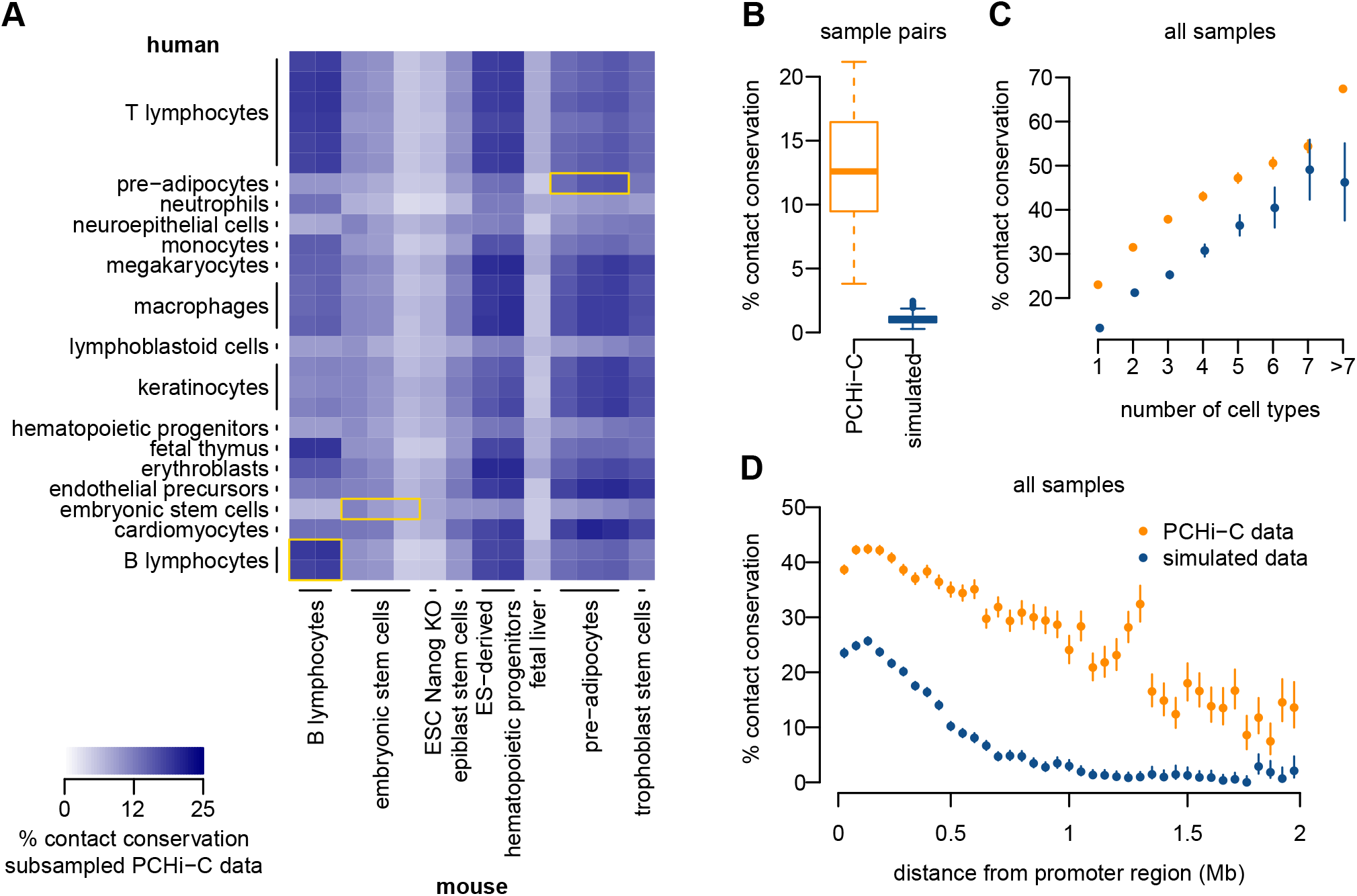
Chromatin contacts between promoters and enhancers are conserved. **A**. Heatmap representing the frequency of contact conservation in comparisons between pairs of PCHi-C samples (one human sample and one mouse sample). We sub-sampled the PCHi-C data to obtain the same numbers of interactions for each sample (Methods). Yellow squares highlight comparable cell types. **B**. Distribution of the frequency of contact conservation between all pairs of samples, for PCHi-C data (orange) and for simulated data (blue). **C**. Proportion of human promoter-enhancer contacts conserved in mouse, as a function of the number of human cell types in which interactions are observed. **D**. Proportion of human promoter-enhancers contacts conserved in mouse, as a function of the distance between the two elements in the human genome. **B-C**. Bars represent 95% two-sided confidence intervals for the proportions (Methods).

For both PCHi-C and simulated data, the proportion of conserved contacts between promoters and enhancers decreases as the genomic distance between the two increases (Fig. 5D, Supplemental Text), as expected given the over-representations of contacts at relatively short distances in both species (Fig. 1, Supplemental Fig. S1). However, here again, the excess of contact conservation compared to simulated data is stronger at large genomic distances: for example, around 1 Mb, the proportion of conserved contacts is 0.99% for simulated data, while for PCHi-C data the value is 25.78% (Fig. 5D). We observed an enrichment for functional categories related to developmental patterning among genes that have high conservation levels for long-distance (minimum 500 kb) promoter-enhancer contacts (Supplemental Table S4). We also show that the most highly constrained human genes (Lek et al., 2016, Methods) have a higher rate of chromatin contact conservation (Supplemental Fig. 13). These observations are consistent with the presence of strong functional constraints on the *cis*-regulatory landscapes of developmental genes, and more generally of dosage-sensitive genes.

### The complexity of *cis*-acting regulatory landscapes is associated with gene expression characteristics and with the rates of gene expression evolution

To better understand the functional relevance and the phenotypic implications of the promoter-enhancer interactions predicted with PCHi-C data, we examined their relationship with gene expression evolution. We evaluated gene expression patterns using a comparative transcriptome collection spanning several organs and developmental stages (Cardoso-Moreira et al., 2019). This dataset allowed us to identify changes in expression profiles, such as changes in organ or developmental stage “preference”, the gain or loss of an expression domain, etc. We measured the extent of expression conservation through Spearman’s correlation coefficient between relative expression values, for each pair of orthologous genes (Methods). Given that gene expression levels and gene expression breadth are correlated with our estimates of the rate of expression profile evolution (Supplemental Fig. S15), we corrected for the effect of these two factors with a multiple regression model (Methods). We repeated all analyses using the Euclidean distance to contrast orthologous gene expression profiles, and obtained similar results (Supplemental Fig. S16, Supplemental Fig. S17, Methods). Genes with the highest expression profile conservation levels are enriched in processes related to RNA metabolism and transcriptional regulation (Supplemental Table S5).

We observe that genes that are in contact with a large number of predicted enhancers in PCHi-C data exhibit higher average expression levels, for both human and mouse (Kruskal-Wallis test, p-value <10^−10^, Fig. 6A). This confirms previous observations showing that the number of enhancers in the gene vicinity is positively correlated with expression levels (Berthelot et al., 2018). We also show that these genes generally have broad expression patterns, that is, they are expressed in large numbers of samples (Kruskal-Wallis test, p-value < 10^−10^, Fig. 6B). This suggests that the contact with a large number of enhancers may enable gene activation in a wide variety of spatio-temporal contexts. Consistent with the enrichment observed for genes with conserved expression profiles, we note that functional categories associated with regulation of expression are over-represented among genes with a large number of contacted enhancers (Supplemental Table S6).

**Figure 6.**
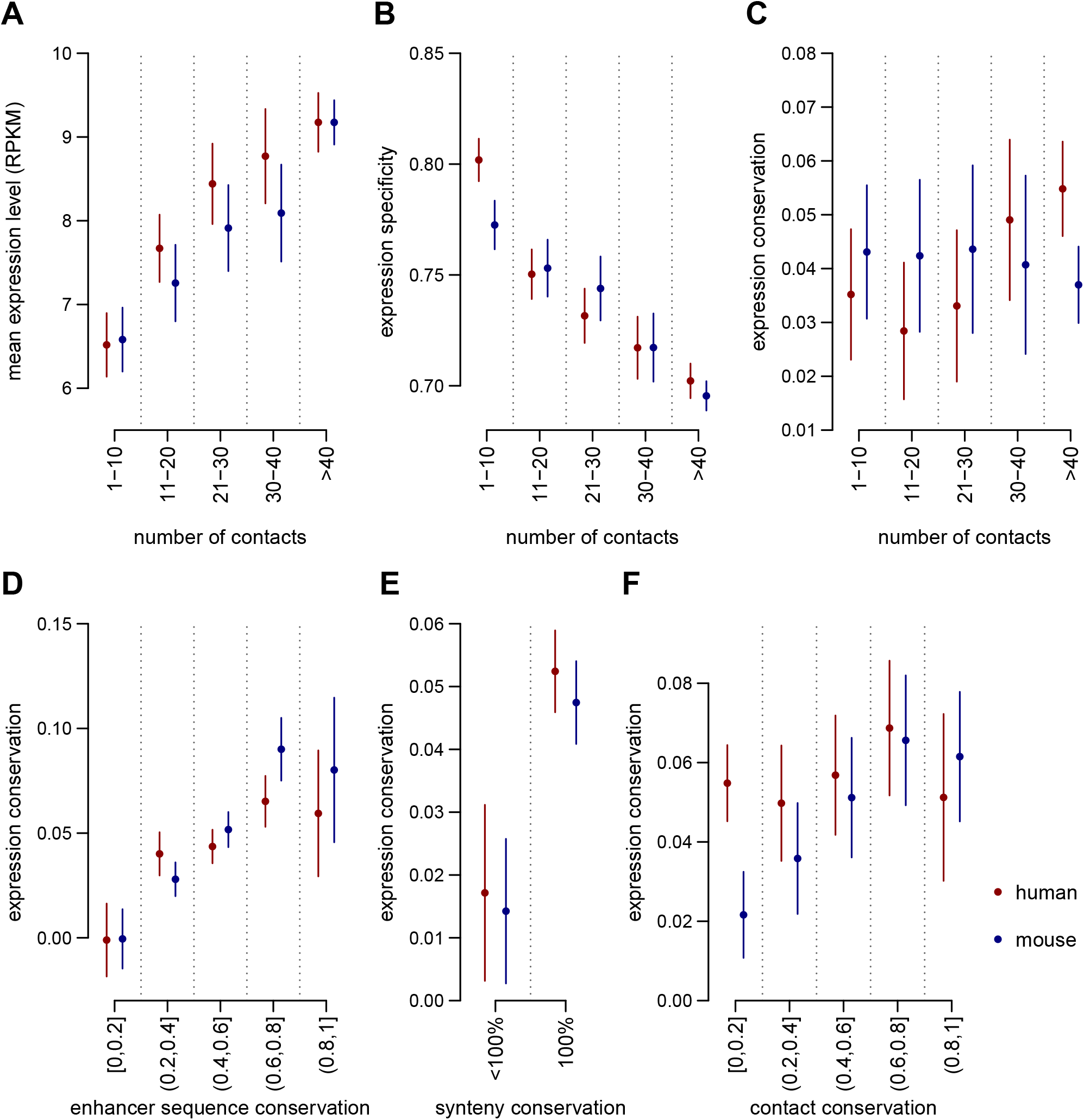
The complexity and the evolution of *cis*-regulatory landscapes are associated with gene expression evolution. **A**. Average expression levels as a function of the number of contacted enhancers in PCHi-C data, for human (red) and mouse (blue). **B**. gene expression specificity index (tau, which ranges from 0 for housekeeping genes to 1 for highly specific genes; Methods) as a function of the number of contacted enhancers. **C**. Expression conservation as a function of the number of contacted enhancers in PCHi-C data. **D**. Expression conservation as a function of the average sequence conservation of contacted ENCODE enhancers. **E**. Expression conservation, depending on whether or not genes underwent at least one break of synteny with the contacted enhancers between human and mouse genomes. **F**. Expression conservation as a function of the proportion of promoter-enhancers contacts conserved in the other species’ PCHi-C data. **A-F**. Dots represent median values across all genes in a class, vertical segments represent 95% confidence intervals for the median. **C-F**. Gene expression conservation is measured with Spearman’s correlation coefficient between human and mouse relative expression profiles, for pairs of 1-to-1 orthologous genes, across organs and developmental stages (expression data from Cardoso-Moreira et al., 2019). Expression conservation is further corrected to account for the effect of expression levels and of expression specificity with a multiple linear regression model (Methods). Enhancer predictions are taken from ENCODE. Expression conservation values are the same for both species, but PCHi-C contact maps differ.

Our results thus confirm that the complexity of the regulatory landscape is linked to the pattern of gene expression (Berthelot et al., 2018). However, after correcting for the effect of gene expression levels and expression specificity, our expression conservation measure is not strongly correlated with the number of contacted enhancers (Kruskal-Wallis test, p-value = 3.4 × 10^−3^ for human, p-value = 0.85 for mouse, Fig. 6C). These observations indicate that the complexity of *cis*-acting regulatory landscapes - as measured with chromatin interaction data - contributes to the robustness of gene expression during evolutionary time, but only through the association between the number of regulatory elements, expression level and expression breadth. The numbers of enhancers assigned to genes with the genomic proximity approach (Methods) do correlate positively with gene expression conservation (Kruskal-Wallis test, p-value < 10^−10^ for both species, Supplemental Fig. S18), as previously reported (Berthelot et al., 2018, Danko et al., 2018). The distance to the next promoter, which is the only determinant of the numbers of enhancers attributed to genes with this approach, also correlates positively with the extent of expression conservation (Kruskal-Wallis test, p-value < 10^−10^ for both species, Supplemental Fig. S18). These observations may be explained by the strong enrichment of developmental functions among the genes with large numbers of neighbor enhancers (Supplemental Table S7). Thus, they may reflect the genomic architecture of developmental genes, rather than their pattern of chromatin interactions.

We also evaluated the conservation of gene expression levels using transcriptome sequencing data for the cell types sampled in both human and mouse PCHi-C datasets (Methods, Supplemental Table S8). We confirm that there is a strong positive correlation between the number of contacted enhancers and gene expression levels in each cell type (Kruskal-Wallis test, p-value < 10^−10^, Supplemental Fig. S19). However, as for the conservation of gene expression profiles, we do not find a significant association between the number of contacted enhancers and the conservation of gene expression levels after correcting for the effect of the gene expression level within species (Kruskal-Wallis test, p-value > 0.05 for all comparisons, Supplemental Fig. S19).

### The evolution of *cis*-regulatory landscapes is correlated with the evolution of gene expression profiles

We next investigated the relationship between the evolution of *cis-*regulatory landscapes and the evolution of gene expression patterns. We evaluated the evolutionary conservation of PCHi-C-predicted regulatory landscapes at three different levels: the conservation of the contacted enhancer sequences, of the synteny between promoters and enhancers, and of their chromatin contacts (Methods). We correlated these measures of regulatory landscape evolution with the evolution of gene expression profiles, corrected for the effect of expression level and expression breadth as described above (Methods).

We found significant positive associations between the rate of regulatory landscape evolution and the rate of gene expression profile evolution (Fig. 6D-F, Supplemental Fig. S16, Supplemental Fig. S17). Specifically, genes that contact enhancers with highly conserved sequences tend to have well-conserved expression profiles (Kruskal-Wallis test, p-value < 10^−10^ for both species, Fig. 6D). We observe a similar correlation when measuring enhancer sequence evolution with phyloP scores (Supplemental Fig. S20). Moreover, genes that underwent synteny breaks in their regulatory landscapes tend to have less conserved expression profiles than genes in conserved synteny (Kruskal-Wallis test, p-value 4.7 × 10^−5^ for human, p-value 1.2 × 10^−6^ for mouse, Fig. 6E). Finally, we tested the correlation between the proportion of conserved contacts and the rate of expression profile conservation. Although we observe a positive tendency, we could not conclude on the presence of a consistent significant signal for both species (Kruskal-Wallis test, p-value 0.26 for human, 5 × 10^−4^ for mouse). Similar results are found when we contrast gene expression profiles between species with the Euclidean distance (Supplemental Fig. S17). These conclusions are also confirmed by multiple regression models that explain the rate of gene expression profile evolution as a function of gene expression characteristics (gene expression level and gene expression specificity) and regulatory landscape characteristics (Supplemental Table S9). We note that we did not observe significant correlations between quantitative expression level differences and *cis*-regulatory landscape evolution in the cell types that were sampled for both mouse and human (Supplemental Fig. S19).

To test whether the genomic distance between interacting promoters and enhancers has an effect on the robustness of gene expression, we analyzed separately medium range interactions (genomic distance below 500 kb) and long range interactions (genomic distance above 500 kb). We found significant associations between the conservation of regulatory landscapes, on one hand, and gene expression patterns and gene expression conservation, on the other hand, for both distance classes (Supplemental Fig. S21). However, the effect is generally weaker for long-range interactions, suggesting that distal regulatory elements contribute less to gene expression evolution than proximal elements (Supplemental Fig. S21). We also found significant associations between the degree of sequence conservation of neighbor enhancers and the extent of gene expression conservation (Kruskal-Wallis test, p-value = 2.1 × 10^−6^ for human, p-value = 1.4 × 10^−10^ for mouse, Supplemental Fig. 22). However, these associations are less strong than the ones observed with the PCHi-C data (above) and the presence of synteny breaks does not correlate significantly with gene expression conservation with the neighbor enhancer dataset (Supplemental Fig. 22).

## Discussion

In this study, we investigated the evolution of *cis-*regulatory landscapes using experimentally determined promoter-enhancer contacts. This data allowed us to evaluate long-range promoter-enhancer interactions, which are thought to be critical part of *cis*-regulatory networks (Montavon and Duboule 2012). The use of chromatin contact data avoids the simplification made by most previous studies, which predicted target genes based on genomic proximity alone (McLean et al., 2010; Villar et al., 2015; Wong et al., 2017; Berthelot et al., 2018; Danko et al., 2018; Dukler et al., 2020). The PCHi-C data revealed *cis*-regulatory landscapes that are more complex than those predicted based with the “genomic proximity approach”: genes are assigned higher numbers of enhancers, and a larger fraction of these enhancers are situated far away from their predicted target genes.

We were able to assess the evolution of *cis*-regulatory landscapes at multiple levels. Starting with primary sequence analyses, we show that regions contacted by promoters in PCHi-C data are better conserved than in simulated data, even after correcting for confounding factors such as repetitive sequence content, GC content etc. Moreover, distant regulatory elements have higher overall levels of sequence conservation than those found in the immediate vicinity of their putative target genes, largely due to a lower overlap with repetitive sequences. Enforcing the idea that long-range contacts between promoters and enhancers are an important mode of gene expression regulation, we showed that synteny breaks that would prohibit these interactions are under-represented, and thus potentially counter-selected. Moreover, we observed substantial contact conservation for long-range promoter-enhancer pairs, though very little is expected by chance. These results confirm early hypotheses stating that the presence of long-range regulatory interactions constrains the large-scale evolution of vertebrate genomes (Mongin et al. 2009; Lemaitre et al. 2009). We thus validate previous computational studies, which predicted long-range regulatory interactions based on long-term conservation (Mongin et al., 2009; Clément et al., 2020). Our findings are also consistent with a recent comparative analysis of human and chimpanzee Hi-C data, which proposed that changes in 3D genome structure may contribute to regulatory evolution (Eres et al., 2019).

We re-evaluated the relationship between gene expression evolution and regulatory evolution. Although it seems intuitive that changes in *cis*-regulatory landscapes should affect gene expression, it is well established that, overall, protein-coding gene expression patterns evolve slowly (Necsulea and Kaessmann, 2014), while distal *cis*-regulatory elements such as enhancers evolve rapidly (Cheng et al., 2014; Villar et al., 2015). Consistently, so far only mild associations between expression evolution and regulatory evolution were reported in vertebrates (Pai et al., 2011; Zhou et al., 2014; Wong et al., 2015; Berthelot et al., 2018). The robustness of regulatory networks, achieved through the presence of redundant *cis*-regulatory elements, is a plausible explanation for this paradoxical finding (Frankel et al., 2010; Cannavò et al., 2016; Osterwalder et al., 2018). Consistently, it was previously shown that the number of enhancers attributed to genes is positively associated with gene expression conservation (Berthelot et al., 2018; Danko et al., 2018). However, this result was obtained with the traditional approach of inferring regulatory relationships based on genomic proximity. With this approach, genomic architecture plays an important role, because the size of the neighboring intergenic and intronic regions directly influences the number of enhancers assigned to a given gene. Thus, genes involved in developmental processes or transcriptional regulation, which can be surrounded by large gene deserts (Montavon and Duboule, 2012), tend to have large numbers of regulatory elements attributed to them (Supplemental Table S7). Because these genes need to be tightly regulated to avoid deleterious phenotypic consequences, this functional enrichment could explain part of the positive association between regulatory landscape complexity and gene expression robustness. These observations raise the question of whether expression robustness is achieved not only through the number of regulatory elements that are available to genes, but also through the evolution of a specific genomic architecture. For example, the presence of large intergenic regions around developmental genes may contribute to the “resilience” of their expression patterns during evolution, by preventing unwanted transcriptional or regulatory interference (Montavon et al., 2011).

With PCHi-C data, we do not observe a strong association between the numbers of contacted enhancers and gene expression conservation, beyond what is explained by expression levels and expression breadth. This puzzling observation might be explained by the fact that regulatory relationships inferred with PCHi-C data are to a great extent orthogonal to genomic architecture: enhancers contacted by promoters are not necessarily their immediate neighbors. Another explanation may reside in the discrepancy between the PCHi-C, gene expression and enhancer datasets that we used here, in terms of biological sampling. Unlike previous studies, which analyzed genes and enhancers that are active in the same tissue (Berthelot et al., 2018), here we rely on heterogeneous sample collections for the different types of data. Despite this drawback, we could uncover significant associations between the rate of regulatory landscape evolution and the pattern of gene expression evolution, by analyzing relative expression profiles across comparable organs and developmental stages (Cardoso-Moreira et al., 2019). Although it does not include biological samples directly related to organ development, there is evidence that our PCHi-C dataset provides a good starting point to study its underlying *cis*-regulatory landscapes. For example, we found that genes that have evolutionarily conserved chromatin contacts at large genomic distances are enriched in functional categories associated with embryonic development. Analyzing expression profiles with this comparable transcriptome collection has the advantage of reducing technical biases linked to gene expression comparisons between distant species, as well as of providing a broader overview of the pattern of expression evolution. Despite the functional redundancy of *cis*-regulatory networks, by jointly analyzing numerous biological conditions we increase the likelihood of observing the molecular consequences of enhancer evolution. Nevertheless, performing similar analyses with comparable PCHi-C and transcriptome sequencing data would likely reveal even stronger relationships between regulatory landscape evolution and gene expression evolution.

We show that genomic rearrangements that affect *cis*-regulatory landscapes are associated with increased divergence of expression profiles. By partially restructuring *cis-*regulatory landscapes, genomic rearrangements likely contribute to gene expression evolution, not just by disrupting existing regulatory relationships, but also by redistributing regulatory elements and thus allowing their adoption by other genes (Lettice et al., 2011). These effects on gene expression explain why rearrangements are generally counter-selected, as indicated by our synteny conservation analyses. Our findings offer an intermediate point of view between reports that large-scale rearrangements that perturb regulatory landscapes can have strong phenotypic consequences, in mouse models of human diseases (Lupiáñez et al., 2015), and reports that multiple chromosomal rearrangements in *Drosophila* laboratory strains have little to no effects on gene expression (Ghavi-Helm et al., 2019). We note that our work, like previous studies, does not provide a complete overview of the phenotypic consequences of regulatory landscape rearrangements. On one hand, studies that were motivated by the need to understand the genomic underpinnings of human diseases are necessarily biased towards events with deleterious consequences (Lupiáñez et al., 2015). On the other hand, studies of *Drosophila* strains are likely biased towards genomic alterations with little impact on organism fitness (Ghavi-Helm et al., 2019). Here, we can only observe those genomic rearrangements that were maintained during evolution, and thus also exclude events with highly deleterious effects. This inherent limitation may explain why we do not observe stronger correlations between gene expression evolution and regulatory evolution. Moreover, we only analyze genes that are kept as orthologs between human and mouse. We speculate that cases where promoter-enhancer interactions are affected by major evolutionary events, such as large-scale genome rearrangements, could often lead to loss of function or pseudogenization, rather than gene expression profile changes.

We note that recent studies have proposed that the presence of chromatin contacts or loops between promoters and enhancers may be dispensable for gene activation in mammals (Alexander et al., 2019; Benabdallah et al., 2019) and in *Drosophila* (Heist et al., 2019; Ing-Simmons et al., 2021). This idea challenges the model of gene regulation centered on contacts between promoters and regulatory elements, built over the last decade by chromatin conformation studies (de Laat and Duboule, 2013; Schoenfelder and Fraser, 2019). The PCHi-C technique, like other “C” techniques, cannot inform on the precise molecular mechanisms that underlie the physical proximity between genomic regions. Thus, the promoter-enhancer interactions that we analyze here may be the result of more complex cellular dynamics, such as the one described in the “phase separation” model (Hnisz et al., 2017). Nevertheless, irrespective of the underlying molecular process, our results support the idea that interactions between promoters and enhancers separated by large distances in the linear genome are a critical part of the complex regulatory networks that control gene expression in mammals.

## Methods

### Promoter Capture Hi-C data processing

We collected and processed publicly available Promoter Capture Hi-C (PCHi-C) data for human (Choy et al., 2018; Freire-Pritchett et al., 2017; Javierre et al., 2016; Mifsud et al., 2015; Pan et al., 2018; Rubin et al., 2017) and mouse samples (Comoglio et al., 2018; Koohy et al., 2018; Novo et al., 2018; Schoenfelder et al., 2015, 2018; Siersbæk et al., 2017). We selected PCHi-C datasets that were generated with experimental procedures similar to those described by Schoenfelder and co-authors (Schoenfelder et al., 2015). Genome fragmentation was generated with the HindIII restriction enzyme in all cases, ensuring identical restriction maps across all samples within a species. We processed PCHi-C data for 26 samples and 16 cell types for human and 14 samples and 8 cell types for mouse (Supplemental Table S1). The data include several cell types (embryonic stem cells, epiblast-derived stem cells, adipocytes and B cells) that are comparable across species, although cell culture procedures and differentiation stages may differ (Supplemental Table S1).

To homogenize computational analyses across datasets and species, we re-processed all PCHi-C raw data. We used the HiCUP pipeline (Wingett et al., 2015), which aligns reads, filters artifactual fragments (such as circularized reads and re-ligations), and removes duplicates. We mapped PCHi-C reads to the human hg38 (GRCh38.p12) and mouse mm10 (GRCm38.p6) genome assemblies, downloaded from Ensembl release 94 (Cunningham et al., 2019), using Bowtie version 2.3.4.3 (Langmead and Salzberg, 2012). We called interactions with the CHiCAGO pipeline (Cairns et al., 2016). We selected chromatin contacts with a CHiCAGO score greater or equal to 5 in at least one sample (Supplemental Table S1). An interaction between a bait and a restriction fragment is said to be detected in a given cell type if it was detected in at least one of the corresponding samples. We combined detected interactions across samples and found 910,180 unique interactions between 19,389 baited restriction fragments and 308,359 other fragments for human, and 824,406 interactions between 21,858 baited fragments and 247,668 other fragments for mouse. These data are provided in Supplemental Dataset S1.

We further focused on intra-chromosomal contacts *(cis*-interactions) separated by a genomic distance of 25 kb to 2 Mb, computed between the midpoint of baited and contacted regions. We thus exclude short-range interactions that have high levels of background noise (Cairns et al., 2016), as well as interactions that are beyond the typical size observed for topologically-associating domains (Dixon et al., 2012). We discarded interactions that involved restriction fragments smaller than 150 bp or larger than 50 kb. We also removed interactions probably involved in structural variation or potential genome assembly issues, *i*.*e*. pairs of contacting regions separated by a large distance in a reference species (>1Mb), and by a small distance (<100kb) in the target species. The list of excluded restriction fragments is provided in Supplemental Dataset S1. Finally, we restricted our analyses to contacts between baited and non-baited restriction fragments.

### Simulated interactions

We generated simulated interaction landscapes that reproduce the observed distribution of distances between baited restriction fragments and non-baited contacted fragments, as well as the number of contacted fragments *per* bait, for each sample (Supplemental Text). To do this, we computed the absolute linear genomic distance between the center position of each baited fragment and each interacting fragment. We then divided the observed interactions into 5 kb distance classes, from 25 kb to 2 Mb upstream and downstream of the baited region. We computed the fraction of contacts observed in each distance class, across all baited fragments, for each sample. This distance distribution was used to simulate contacts, as follows: for each baited fragment, we computed the contact probability for all fragments found on the same chromosome, within the 25 kb - 2 Mb distance window, as the average probability of the overlapping distance classes (a single fragment can overlap with multiple distance classes). We then randomly drew contacts among the list of all possible interactions based on this empirical probability distribution. We respected the number of interactions observed in the real PCHi-C data, for each bait. However, we discarded *a posteriori* those simulated interactions that fell within a baited restriction fragment; after this filtering step, the number of contacts *per* bait are lower in the simulated data than in the PCHi-C data (Supplemental Fig. 1). The simulated interaction data are available in Supplemental Dataset S2.

### Theoretical mappability and PCHi-C read mapping statistics

We evaluated the theoretical mappability of all restriction fragments by drawing artificial sequencing reads from the genome and re-mapping them with Bowtie 2 (Langmead and Salzberg, 2012), with the same parameters as the HiCUP pipeline. The starting points of the reads were spaced by 5 nucleotides. We did not simulate sequencing errors. Regions for which artificial reads were aligned unambiguously to their original location were said to be mappable. We computed the percentage of mappable bases and the maximum mappable stretch (the largest perfectly mappable interval) for each restriction fragment. As several read lengths were available for each species, we repeated this procedure for each read length and computed the minimum values. We also estimated the actual number of mapped PCHi-C reads attributed to each fragment in each sample, using BEDTools utilities (Quinlan and Hall, 2010). We discarded restriction fragments that had a maximum theoretical mappable stretch lower than 150bp and fewer than 50 mapped PCHi-C reads, combined across all samples.

### Subsampled chromatin interaction dataset

To minimize differences in detection sensitivity among PCHi-C samples (Supplemental Table S1, Supplemental Text), we generated subsampled datasets. We first computed the minimum number of observed interactions (N) across all samples for each species (79,843 for human and 70,475 for mouse). We then ranked interactions based on their CHiCAGO score and kept the strongest N interactions from each PCHi-C sample, reasoning that the relative ranking would remain unchanged if detection power were reduced. For simulated datasets we randomly re-sampled N interactions. We applied the same filtering steps described previously (e.g., discarding bait-bait interactions, *trans* interactions and interactions occurring at distances < 25 kb or > 2 Mb) on the subsampled data before analyzing them further. We used these subsampled interaction datasets to evaluate sample clustering within species (Supplemental Fig. S3, Supplemental Text), and for pairwise comparisons of contact conservation between human and mouse samples (Fig. 5A,B, Supplemental Text).

As PCHi-C and simulated datasets differ in terms of total numbers of interactions after pooling all available samples (Supplemental Text), we randomly subsampled the pooled simulated dataset to obtain the same number of interactions as in pooled PCHi-C data. These datasets were used in the contact conservation analyses that rely on pooled samples across species (Fig. 5C,D, Supplemental Text) and are available in Supplemental Dataset S1 and Supplemental Dataset S2.

### Sample clustering

We evaluated the similarity between pairs of samples from the same species starting from the percentage of shared interactions, *i*.*e*. 100 times the ratio between the number of shared interactions and the number of interactions observed in at least one of the samples. For each pair of samples, we computed the difference between the percentage of shared interactions in PCHi-C data and the percentage of shared interactions in simulated data. We computed this measure of similarity on the subsampled PCHi-C and simulated data. We used this pairwise measure of similarity to cluster samples within a species, using the hierarchical clustering approach implemented in the “hclust” function in R (R Core Team, 2014). We also used functions within the “ade4” R package (Dray et al., 2007) to perform a correspondence analysis for each species, starting from a contingency table describing for each unique chromatin interaction whether it was observed or not in each sample. We performed this analysis on the subsampled PCHi-C dataset.

### Baited region annotation

We assigned transcription start sites to PCHi-C baited restriction fragments using annotations from the Ensembl database, release 94. We downloaded transcript coordinates from Ensembl using the BioMart interface (Kinsella et al., 2011). For each baited fragment, we extracted all transcription start sites that were found within at most 1 kb of the fragment. The baited fragment annotation is available in Supplemental Dataset S1; gene and transcript annotations are provided in Supplemental Dataset S3.

### Genomic characteristics of PCHi-C contacted sequences

We extracted the repeat-masked DNA sequence of all baits and contacted regions, and used BLAT (Kent et al., 2012) to search for sequence similarity in the same genome. For each sequence, we counted the number of BLAT hits corresponding to at least 80% of their repeat-masked length with at least 95% sequence identity. We discarded sequences with more than 1 BLAT hit, which could be potentially duplicated in the reference genome. We evaluated the repetitive sequence content for restriction fragments and enhancers using RepeatMasker annotation tables provided by the UCSC Genome Browser (Hinrichs, 2006). We evaluated the number of protein-coding genes found within a maximum distance of 500 kb, upstream and downstream of the contacted sequences. We also analyzed the GC content of restriction fragments and enhancers. These genomic characteristics are available in Supplemental Dataset S1 for restriction fragments and in Supplemental Dataset S4 for enhancers.

### Predicted enhancer elements

We evaluated the presence of predicted enhancer elements in the PCHi-C contacted regions using different data. For human, we used pre-filtered data from a recent study (Hait et al., 2018) (http://acgt.cs.tau.ac.il/focs/download.html), obtaining 408,802 enhancer positions predicted with DNase I hypersensitivity (DHS) assays by the ENCODE consortium (Thurman et al., 2012). For mouse, we extracted enhancer positions predicted based on the presence of H3K4me1, H3K4me3 and H3K27ac histone modifications from the ENCODE consortium (The Mouse ENCODE Consortium et al., 2014). We downloaded the midpoint coordinates of predicted enhancer elements for 23 tissues or cell lines from http://promoter.bx.psu.edu/ENCODE/download.html and we extended them by 75 bp on each side. We then combined the enhancer coordinates across all samples, obtaining 740,058 enhancer regions.

To confirm our results, we also used human enhancers prediction from the Roadmap Epigenomics consortium (Roadmap Epigenomics Consortium et al., 2015), and from global run-on sequencing analyzed in the same study (Hait et al., 2018) (http://acgt.cs.tau.ac.il/focs/download.html). Moreover, we downloaded enhancers predicted with the Cap Analysis of Gene Expression method by the FANTOM5 consortium (Andersson et al., 2014) for human (https://zenodo.org/record/556775#.X3Gvf5rgprl) and mouse (https://zenodo.org/record/1411211#.X3Gvq5rgprm).

We converted enhancer coordinates to the latest genome assembly of each species if needed (hg38 for human and and mm10 for mouse) using liftOver and associated genome alignment files downloaded from the UCSC Genome Browser (Kent et al., 2012). Finally, we applied the same procedure described above for restriction fragments to discard duplicated enhancers and to evaluate their genomic characteristics (GC content, repetitive sequences, proximal gene density). The predicted enhancers data and their genomic characteristics are available in Supplemental Dataset S4.

### Prediction of contacts between gene and enhancers

We constructed gene-enhancer pairs by associating to each “baited” protein-coding gene all the predicted enhancers that overlapped with the restriction fragments contacted by its baits. We applied this procedure to each enhancer dataset described above, obtaining four different contact datasets for human and two datasets for mouse (Supplemental Dataset S4).

### Prediction of gene-enhancer association using the genomic proximity approach

We inferred regulatory relationships between promoters and enhancers using the genomic proximity approach, as implemented by Berthelot and co-authors (Berthelot et al 2018). We first determined for each gene the canonical transcript, based on Ensembl APPRIS annotations when available (Cunningham et al., 2019). For genes that did not have APPRIS annotations, we kept the transcripts with the longest coding sequence (for protein-coding genes) or with the longest exonic sequence (for non-coding RNAs). For this analysis, we kept only protein-coding genes, long non-coding RNAs and antisense transcripts. Each gene was assigned a unique transcription start site (TSS), belonging to the canonical transcript. Then, we defined for each gene a putative regulatory region delimited by the closest TSS upstream and downstream of the gene’s TSS. Enhancers found within this region were then assigned to the focal gene. We restricted promoter-enhancer relationships defined with the genomic proximity approach to the same 25kb to 2Mb distance interval used for the PCHi-C data. The corresponding data are available in Supplemental Dataset S4.

### Correlated activity of gene-enhancer pairs

We evaluated the correlation of gene expression and enhancer activity levels for each promoter-enhancer pair, across samples. Depending on the dataset, activity levels were evaluated with ChIP-seq or DNase I hypersensitivity experiments (ENCODE, RoadmapEpigenomics consortia), with the CAGE technique (FANTOM5 consortium) or with the GRO-seq technique. In all cases, we used processed promoter and enhancer activity data (Hait et al., 2018). We downloaded normalized activity profiles for promoters and enhancers from http://acgt.cs.tau.ac.il/focs/download.html. We computed pairwise Spearman correlations, based on normalized activity profiles across samples, for all pairs of promoters - enhancers in contact in the real PCHi-C data or in the simulated data. We then tested whether the correlation coefficient distributions were significantly different, using the Wilcoxon rank-sum test for median comparisons, as implemented in R (R Core Team, 2014). The resulting data are available in Supplemental Dataset S12.

### Sequence conservation

To evaluate the conservation of sequences contacted in PCHi-C data, we first identified putative homologous regions using liftOver on whole-genome alignments between a reference species (human or mouse) and a target species (human, macaque, mouse, rat, rabbit, cow, elephant, dog, opossum or chicken), downloaded from the UCSC Genome Browser (Hinrichs, 2006). We set a low threshold (10%) for the minimum ratio of bases that must remap in the liftOver conversion. We discarded regions that were duplicated or split in the target genome. The regions that could not be projected with liftOver were considered as non-conserved and were given a conservation score of 0. The predicted homologous regions were then aligned with Pecan (Paten et al., 2008). We computed the conservation score as the ratio between the total number of aligned (without gaps) base pairs and the total number of positions in the alignment. To better evaluate the determinants of sequence conservation patterns in PCHi-C data, we also measured sequence conservation separately for repetitive and non-repetitive sequences, using the information provided in the repeat-masked genome sequence available in Ensembl (Zerbino et al., 2018). We extracted exonic coordinates from the Ensembl database and we masked exons before evaluating sequence conservation. We discarded sequences whose alignment length was smaller than 10 bp. We applied the same alignment procedure to predicted enhancers.

We also analyzed the phyloP basewise conservation score (Pollard et al., 2010). We retrieved from the UCSC Genome Browser (Hinrichs, 2006) phyloP scores calculated from multiple alignments of 30 vertebrate species for human and for 60 vertebrate species for mouse. We computed average phyloP scores for each restriction fragment and enhancer, across all non-exonic bases that had phyloP coverage. We also computed average phyloP scores separately for repetitive and non-repetitive sequences, using RepeatMasker annotations downloaded from the UCSC Genome Browser (Hinrichs, 2006).

The alignment statistics and phyloP scores are available in Supplemental Dataset S6 and in Supplemental Dataset S7.

### Synteny conservation

We defined synteny conservation between a reference and a target species as the presence of the gene and of the predicted enhancer on the same chromosome and at a distance of less than 2 Mb in both species. We restricted this analysis to protein-coding genes with 1-to-1 orthologues in the target species, as predicted in the Ensembl database (release 99 for macaque, release 94 for all other species). We also selected only enhancers that had predicted homologous regions in the target genome, as defined above. For each pair of species, we discarded the 10% least conserved enhancers based on the alignment score defined above. We further restricted this analysis to promoter-enhancer pairs that are separated by distances between 100 kb and 1.5 Mb in the reference species. We then asked whether the predicted homolog of the contacted enhancer was on the same chromosome and within 2 Mb of the TSS of the orthologous gene. If the orthologous gene in the target species had more than one TSS, we considered the minimum distance between these and the homologous contacted region. The resulting data are available in Supplemental Dataset S8 and in Supplemental Dataset S9.

### Chromatin contact conservation

We restricted this analysis to protein-coding genes with 1-to-1 orthologues in the target species, in contact with enhancers that had predicted homologs in the target species, as described above. We further restricted the analysis to cases where the 1-to-1 orthologous gene was also baited in the PCHi-C data of the target species (human or mouse). To dissociate contact conservation from synteny conservation, we require that the bait and restriction fragment be situated on the same chromosome in the target species. We excluded interactions for which the restriction fragment contacted was baited or the distance bait-contacted fragment was below 25 kb or above 2 Mb, for either the reference or the target species. We then asked whether the predicted homolog of the contacted enhancer overlapped any of the regions found in contact with the bait(s) associated with the orthologous gene in the PCHi-C data of the target species. The proportion of conserved contacts was computed with respect to the number of pairs satisfying all previous criteria (*i*.*e*., homology prediction for the baited gene and the contacted enhancer, presence of baits for the orthologous gene in the target species, distance and unbaited contacted fragment filters).

We evaluated the contact conservation for each pair of human and mouse samples. We performed this analysis on the downsampled dataset, which comprises the same number of contacts for each sample. We also evaluated the extent of chromatin contact conservation for the pooled PCHi-C dataset. In this case, we used as a comparison the subsampled pooled simulated dataset, which matches the number of interactions observed in the pooled PCHi-C dataset. For the analyses that contrast the extent of regulatory landscape conservation and gene expression conservation, we use the entire PCHi-C data, without subsampling.

The resulting data are available in Supplemental Dataset S10 and Supplemental Dataset S11.

### Gene expression data

To evaluate the gene expression patterns, we used expression data for human and mouse across multiple organs and developmental stages (Cardoso-Moreira et al., 2019). We downloaded gene-level RPKM (reads *per* kilobase of exon *per* million mapped reads) values. For evolutionary comparisons, we analyzed protein coding genes predicted as 1-to-1 orthologs for human and mouse in the Ensembl database release 94, and to organ/developmental stage combinations that were directly comparable between the two species (Cardoso-Moreira et al., 2019). We re-normalized the data across samples and species using a median-scaling procedure based on the genes that vary the least in terms of ranks across samples (Brawand et al., 2011). For the expression conservation analyses, we required genes to be expressed above a minimum threshold (RPKM=1) in at least three samples. The resulting expression data and sample details are provided in Supplemental Dataset S5.

We also aimed to compare gene expression levels between human and mouse, in cell types that are comparable between the two species and for which PCHi-C data is also available. We analyzed embryonic stem cells or epiblast-derived stem cells (ESC or epiSC), adipocytes and B cells. For these cell types, we downloaded RNA-seq data from the ENCODE consortium and from the Sequence Read Archive (SRA) database. We computed gene expression levels using kallisto (Bray et al., 2016), on cDNA sequences extracted from Ensembl 94. For each gene, we computed mean and median TPM (transcript *per* million) expression levels, across all replicates for each cell type. We applied the same normalization procedure as above (Brawand et al., 2011). These values were used to compute cell type-specific gene expression conservation values between human and mouse. The resulting expression data and sample details are provided in Supplemental Dataset S5 and Supplemental Table S8.

### Gene expression characteristics

We defined gene expression breadth as the number of organ/developmental stage combinations where the average RPKM level across biological replicates was above 1, using the expression data described above. We analyzed the distribution of this estimate of expression breadth as a function of the maximum number of cell types in which interactions were observed for baited genes. We also computed a tissue/developmental stage specificity index with the formula tau = sum (1 – r_i_)/(n-1), where r_i_ represents the ratio between the expression level in sample i and the maximum expression level across all samples, and n represents the total number of samples (Liao et al., 2006). Genes with perfectly homogeneous expression levels across all samples thus have a tau value of 0, while genes expressed in a single condition have a tau value of 1. We computed this index on RPKM values, averaged across all replicates for a given species / organ / developmental stage combination.

### Evolutionary conservation of gene expression profiles

To evaluate the conservation of gene expression patterns between human and mouse, we first computed relative expression profiles, by dividing the RPKM values (averaged across biological replicates) by the maximum value observed among samples for each gene. We used the transcriptome data described above (Cardoso-Moreira et al., 2019). We measured expression conservation as Spearman’s correlation coefficient between the relative expression profiles of orthologous genes. We also measured expression conservation as 1-d, where d is the Euclidean distance between orthologous gene expression profiles. As these measures of expression conservation are significantly correlated with the average gene expression level and with the expression specificity index (Supplemental Fig. S15), we built linear regressions that model the relationship between expression conservation, expression specificity and expression levels (averaged across all samples and across species, for each gene), and extracted the residual values. We referred to these residual values as “corrected expression conservation” in the figures. The gene expression conservation data is provided in Supplemental Dataset S5.

We analyzed the factors associated with gene expression evolution with multiple regression models (Supplemental Table S9). The response variable in these models is the conservation of gene expression profiles, measured by Spearman’s correlation coefficient before and after correction. The explanatory variables are various regulatory landscape characteristics (number of contacted enhancers, sequence conservation for contacted enhancers, synteny conservation, contact conservation) and gene expression characteristics (gene expression level, gene expression specificity).

### Evolutionary conservation of gene expression levels

We measured quantitative gene expression differences between species for the three cell types that were present in the PCHi-C datasets of both human and mouse: B lymphocytes, pre-adipocytes and embryonic stem cells. We also performed a comparison between human embryonic stem cells and mouse epiblast-derived stem cells. To do this, we computed average TPM levels across all replicates available for a given cell type, for each species. We then estimated expression divergence as the absolute value of the difference between human and mouse, divided by the maximum of the two values, for each gene. We note that this expression divergence measure is strongly correlated with gene expression levels. To correct for this effect, we constructed a linear regression that models the relationship between the expression divergence and the average expression levels, across both species, and extracted the residual values. Gene expression conservation measures are defined as expression divergence values subtracted from 1. We referred to these values as “corrected expression conservation”. The gene expression conservation data are provided in Supplemental Dataset S5.

### Evolutionary divergence of regulatory landscapes

We performed this analysis on 1-to-1 orthologous protein-coding genes between human and mouse. For each gene pair, we evaluated: the number of enhancers found in contact with the reference gene promoter(s); the sequence alignment score of the contacted enhancers; the number of enhancers with conserved sequences in the target species (*i*.*e*., successfully projected with liftOver in the target genome and with an alignment score greater than or equal to 40%) and found in conserved synteny with the orthologous gene in the target genome (as defined above); the number of enhancers with conserved sequences, whose predicted homolog was also in contact with the orthologous gene in the target genome. We could thus evaluate for each pair of orthologous genes: the median alignment score of contacted enhancers, the percentage of conserved enhancers maintained in synteny and the percentage of conserved contacts. These two last measures were calculated only for genes that had between 5 and 100 conserved enhancers. We then correlated these measures with the gene expression conservation values calculated above.

### Constraints on gene sequence

We analyzed the probability of intolerance to loss-of-function mutations for a gene (pLI score), inferred from variation in human exome-sequencing data leading to truncating proteins (Lek et al. 2016). We downloaded pre-computed pLI scores for transcripts from the website indicated in the original article (ftp://ftp.broadinstitute.org/pub/ExAC_release/release0.3/functional_gene_constraint). We then associated pLI scores to genes using Ensembl94 annotations. We retained the 17,558 genes associated with a unique transcript. The pLI scores are provided in Supplemental Dataset S6.

### Gene Ontology enrichment

We used GOrilla (Eden et al., 2009) to perform Gene Ontology enrichment analyses on single-ranked lists of genes. We considered only protein coding genes present in the human PCHi-C data and with an 1-to-1 orthologous gene in the mouse. Genes were ranked by the following factors: the number of contacted ENCODE enhancers, the number of neighbor enhancers, the median distance between enhancers and the gene TSS, the mean alignment score of contacted enhancers, the proportion of enhancers that are conserved in synteny and in contact, the gene expression profile conservation. The ontology enrichment results are provided in Supplemental Dataset S13 and Supplemental Tables S3-S7.

### Statistics and graphical representations

We used R 3.5.2 for all statistical analyses and graphical representations (R Core Team, 2014). Given that the variables we analyze are often not normally-distributed, we used non-parametric statistical tests as a general rule. We computed 95% confidence intervals for mean values using the BCa non-parametric bias-corrected and accelerated bootstrap method (DiCiccio and Efron, 1996), as implemented in the coxed_0.3.3 package in R (Kropko and Harden, 2019). We performed 100 bootstrap replicates. We computed 95% confidence intervals for median values using the formula +/-1.58 IQR/sqrt(n), where IQR is the inter-quartile range of the distribution and n the total number of values. This formula is implemented in the boxplot.stats function in the grDevices package in R (Chambers et al., 1983). We performed pairwise comparisons of distributions with the Wilcoxon rank sum test and multiple comparisons with the Kruskal-Wallis test, both implemented in the stats package in R. We compared proportions with the chi-squared test and computed two-sided 95% confidence intervals with the prop.test function in R. For all tests, we display p-values as “<10^−10^” if lower values are found. For the analyses where we performed multiple tests, we computed false discovery rates with the Benjamini-Hochberg approach.

## Data access

All processed data generated in this study are available as Supplemental Datasets on the publisher’s website as well as at the following address: http://pbil.univ-lyon1.fr/members/necsulea/RegulatoryLandscapes

All scripts used in this analysis are available at: https://github.com/AlexandreLaverre/Regulatory_Landscape, as well as in the Supplemental Code.

## Competing interest statement

The authors declare no competing interests.

## Acknowledgements

We thank C. Berthelot, Y. Ghavi-Helm, F. Picard and D. Mouchiroud for discussions and advice on the project. We also thank T. Latrille and T. Tricou for their valuable advice and for assistance with bioinformatics and P. Veber for advice on statistics. Computational analyses were performed using the computing facilities of the CC LBBE/PRABI and the Core Cluster of the Institut Français de Bioinformatique (IFB) (ANR-11-INBS-0013). This work was funded by the French National Research Agency (ANR-17-CE12-0019-01 «LncEvoSys»).

## Author contributions

The project was originally designed by A.L., E.T and A.N.; A.L. conducted data analyses with help from A.N; A.L., E.T. and A.N. wrote the manuscript.

## Notes

http://pbil.univ-lyon1.fr/members/necsulea/RegulatoryLandscapes

https://github.com/AlexandreLaverre/Regulatory_Landscape

### Competing Interest Statement

The authors have declared no competing interest.

