## Supplemental Text for "Long-range promoter-enhancer contacts are conserved during evolution and contribute to gene expression robustness"

**2 Supplemental text figures** **6**

**3 Additional figures related to the main and supplemental figures** **12**

### 1 Supplemental text

#### 1.1 Construction of a simulated interaction dataset

To test the biological significance of the observations made with PCHi-C data, we constructed a simulated interaction dataset, which served as a basis for most ensuing analyses. Regulatory evolution studies have traditionally inferred target genes for *cis*-regulatory elements based on genomic proximity [1, 2]. Furthermore, the genomic distance between restriction fragments is the main factor determining the likelihood of observing chromatin contacts [3]. Because of this, we designed our simulated interaction dataset to mimic the distribution of distances between interacting restriction fragments observed in the PCHi-C data.

The PCHi-C technique involves chromatin digestion with a restriction enzyme, in this case HindIII [4]. A certain number of the resulting restriction fragments are targeted using RNA baits; interactions involve at least one (sometimes two) baited restriction fragments. To construct the simulated data set, we used as a starting point the real restriction map obtained by digestion with HindIII and the real baited restriction fragments. For each sample, we compute the distances between baited fragments and un-baited contacted restriction fragments, using the center coordinates of restriction fragments. We then divided distances into 5 kb-wide classes, starting at 25 kb and ending at 2 Mb (e.g., 25-30 kb, 30-35 kb, . . . ). We count the number of contacts occurring in each distance class, combining all interactions across all baits, for each sample. These frequencies are transformed into an empirical probability distribution. We then draw random interactions for each bait: the probability for a restriction fragment to appear in interaction with a baited fragment is computed based on the distance between the two, given the probability distribution computed above. We draw the same number of interactions *per* bait as observed in the real data, before filtering (see below).

#### 1.2 Similarities and differences between real and simulated PCHi-C data

Our simulation procedure ensures that the distribution of distances between baited fragments and contacted fragments is almost identical between real and simulated PCHi-C data, in each sample (Fig. 1, Supplemental Fig. S1). The degree of each bait (*i.e.*, the number of contacted fragments) is also generally respected, although some slight differences arise from filtering procedures that we applied after the initial simulations (selection of un-baited fragments, minimum and maximum contacted fragment size, removal of restriction fragments that are potentially duplicated in the reference species, *etc.*). However, the degree of the contacted fragments (*i.e.* the number of contacting baits) is not respected. In the simulated data there is an excess of restriction fragments that are contacted by a single bait compared to the real PCHi-C data (Supplemental Fig. S1). In the real PCHi-C data, restriction fragments are often contacted by multiple baits. This is expected for genuine regulatory interactions - indeed, a single enhancer can regulate multiple genes. This behavior is not enforced in the simulated data. As a consequence, the total number of distinct restriction fragments that are contacted by at least one bait is always higher in the simulated data than in PCHi-C data (Supplemental Fig. S2). Also as a consequence, the total genomic length covered by contacted restriction fragments is also generally higher in simulated data than in PCHi-C data (Supplemental Fig. S2).

We also note that our simulated PCHi-C data does not reproduce the variability among baits observed in the real PCHi-C data. This is inevitable given that we estimate the distribution of distances between pairs of contacting fragments by pooling all baits; this average distribution is then used to simulate contact data for each bait. To quantify this intuition, we measured the variability among baits, in terms of the distribution of distances between baits and contacted fragments, for each sample. Specifically, for each bait, we counted the number of interactions (observed and simulated) in each 5 kb distance bin, within the 25 kb - 2 Mb distance interval. We then computed the relative abundance of interactions among distance bins, separately for each bait and for each dataset (PCHi-C or simulated). We then computed the difference, measured by an Euclidean distance, between each *per*-bait empirical distance distribution and the average empirical distance distribution across all baits, which was used for the simulations. We show examples for human and mouse in Supplemental Text Fig. 1. This analysis shows that, as expected, the variability among baits is lower for the simulated data than for the PCHi-C data (Supplemental Text Fig. 1). However, globally, the variability among baits in simulated data remains comparable with the variability observed in PCHi-C data.

Interactions are significantly more often shared among samples and among cell types in the real PCHi-C data than in the simulated data (Fig. 1). As a consequence, when combining all samples available for a species in a single dataset, we observe different distributions for the numbers of contacted fragments *per* bait and for the numbers of contacting baits *per* fragment (Supplemental Fig. S2). Namely, in the simulated dataset, combined across all samples, each bait contacts more distinct restriction fragments and each restriction fragment is contacted by more distinct baits than in the PCHi-C data (Supplemental Fig. S2). In contrast, in the pooled PCHi-C data, the same interacting pair of restriction fragments can be encountered multiple times.

Combined across all samples, the simulated data contains a much larger number of contacted restriction fragments than the real PCHi-C data (Table 1). We note that most (92% in human, 78% in mouse) of the fragments contacted in the real PCHi-C data are also found in the simulated data, though not necessarily in the same samples or in contact with the same baits. In contrast, smaller proportions (43% in human, 27% in mouse) of pairs of contacting fragments are found in common in PCHi-C and simulated data (Table 2).

| | PCHi-C data | simulated data | in common |
|---|---|---|---|
| human | 263540 | 569301 | 243300 |
| mouse | 207949 | 376635 | 161362 |

Table 1: Number of restriction fragments contacted by baited fragments in PCHi-C data and in simulated data, for human and mouse. Note that many restriction fragments are present in both real and simulated data, though they are not necessarily contacted by the same baits.

| | PCHi-C data | simulated data | in common |
|---|---|---|---|
| human | 662562 | 1846495 | 283880 |
| mouse | 564678 | 898976 | 153600 |

Table 2: Number of unique contacts (pairs of baits - contacted fragments) in PCHi-C data and in simulated data, for human and mouse.

| | PCHi-C data | simulated data |
|---|---|---|
| human | 930 | 1998 |
| mouse | 644 | 1204 |

Table 3: Total length (Mb) covered by distinct contacted restriction fragments, in PCHi-C data and in simulated data, for human and mouse.

In total, restriction fragments contacted in PCHi-C data cover 930Mb in the human genome and 644Mb in the mouse genome. The genomic length fractions are approximately twice as large in the simulated data.

#### 1.3 Theoretical mappability differences between PCHi-C and simulated data

We analyzed the genomic characteristics of the restriction fragments that are contacted in PCHi-C data or in simulated data and we observed that the latter overlap more frequently with repeats than the former (main text, Supplemental Fig. S8). This observation could be affected by an inherent difference between the PCHi-C dataset and the simulated dataset: restriction fragments that are genuinely contacted in the PCHi-C data have to be mappable, that is, their sequences must be unique enough to enable unambiguous mapping of sequencing reads that stem from them. Because simulated PCHi-c data do not have this requirement, this could induce a difference between the two datasets in terms of repetitive sequence content.

To quantify differences between the PCHi-C and the simulated data in terms of mappability, we first estimated the theoretical mappability of each restriction fragment by drawing artificial reads and remapping them to the genome (Methods). We say that a region is mappable if the sequencing reads that are derived from it can be aligned unambiguously to their original position. For each fragment, we then computed the total fraction of mappable bases, as well as the size of the longest mappable stretch. This latter aspect is relevant because the longer mappable intervals enable correct mapping for multiple overlapping reads. We further normalized the size of the longest mappable stretch to the total size of the restriction fragment. We compared these two parameters between restriction fragments contacted in the PCHi-C data and in the simulated data. For both PCHi-C and simulated datasets, the proportion of mappable bases is very high: 25% quantile is at 98.9% for simulated data and at 99.4% for PCHi-C data, for human; at 98.1% for mouse simulated data and at 99.5% for mouse observed data (Supplemental Text Fig. S2). We draw similar conclusions from the analysis of the longest mappable stretch: both PCHi-C and simulated fragments have very long mappable stretch. The PCHi-C data has better mappability statistics than the simulated data, as expected, but the two are nevertheless comparable.

The difference between PCHi-C and simulated data is more striking in terms of mapped PCHi-C reads, as expected (Supplemental Text Fig. S2). Indeed, restriction fragments that have many mapped PCHi-C reads are likely to be included in the PCHi-C contact data.

#### 1.4 Sequencing depth and number of detected interactions

The PCHi-C samples included in this dataset are derived from many different publications, and are thus not perfectly matched in terms of detection power. Indeed, we observed highly variable numbers of interactions in the different samples, for both human and mouse (Supplemental Text Fig S3). The sequencing depth is an obvious determinant of the number of detected interactions, but we observe some outlier samples, which have good sequencing depth but relatively few interactions (Supplemental Text Fig. S3).

The unequal numbers of interactions detected in the different samples can bias some of our analyses, namely, the analysis of the sample clustering within species (Supplemental Fig. S3), and the contact conservation analysis (Fig. 5). To overcome this bias, we subsampled the same numbers of interactions *per* sample, within each species (Methods). The subsampled data were used for the two analyses cited above.

#### 1.5 Gene expression evolution and regulatory landscape evolution

We quantified the rate of gene expression evolution by contrasting relative expression profiles across comparable samples (organ/developmental stage combinations) for human and mouse, for each pair of 1-to-1 orthologous genes. We used two different measures of expression conservation: Spearman's correlation coefficient and a similarity measure based on the Euclidean distance (Methods). Both expression conservation measures correlate with the average gene expression level and with gene expression breadth (Methods, Supplemental Fig. 15). We corrected for these confounding factors by constructing a multiple regression model of each expression conservation measure against these two variables, and by using as a "corrected" expression conservation measure the residuals of the regression model. With this correction, we found that our Euclidean distance-based measure of expression similarity correlates well with our estimates of regulatory landscape evolution (Supplemental Fig. S16, S17).

For these expression analyses, we used a recently published transcriptome collection, which spans multiple organs and developmental stages. However, this transcriptome dataset is not matched with the cell type collection available in the PCHi-C data. Unfortunately, only three cell types were available for both human and mouse in the PCHi-C data (B lymphocytes, embryonic stem cells and pre-adipocytes). We also analyzed epiblast-derived stem cells in the mouse, for the comparison with human stem cells. Gene expression levels are well conserved between human and mouse: samples tend to group by cell type rather than by species in hierarchical clustering and principal component analyses (Supplemental Text Fig. S4). Human embryonic stem cells tend to correlate better with epiblast-derived stem cells (median Spearman's rho 0.81) than with mouse embryonic stem cells (median Spearman's rho 0.78; Supplemental Text Fig. S4).

Despite the limited data, we note that there are parallel trends in the rates of expression evolution, protein-coding sequence evolution and regulatory landscape evolution among these cell types (Supplemental Fig. 15). Specifically, the immune cell type (B lymphocytes) has rapidly evolving transcriptomes, proteomes and regulomes; in contrast, pre-adipocytes and embryonic stem cells show slower rates of evolution, for all three cellular components (Supplemental Text Fig. 5).

#### 1.6 Analysis of sequence conservation variance for each reference species with multiple regression models

We analysed the variance of sequence conservation for each reference species with multiple regression models (Supplemental Table S2). Their response variable is the sequence conservation, measured by pairwise alignment score or phyloP score, of contacted fragments in PCHi-C and simulated dataset. Their explanatory variables are sequence characteristics: number of repeated bases, number of genes found within 500kb, GC content and median distance from baited fragments.

The proportion of repeated elements, gene density, GC content and the distance to the baits are all highly correlated with measures of sequence conservation (Supplemental Table S2)

#### 2 Supplemental text figures

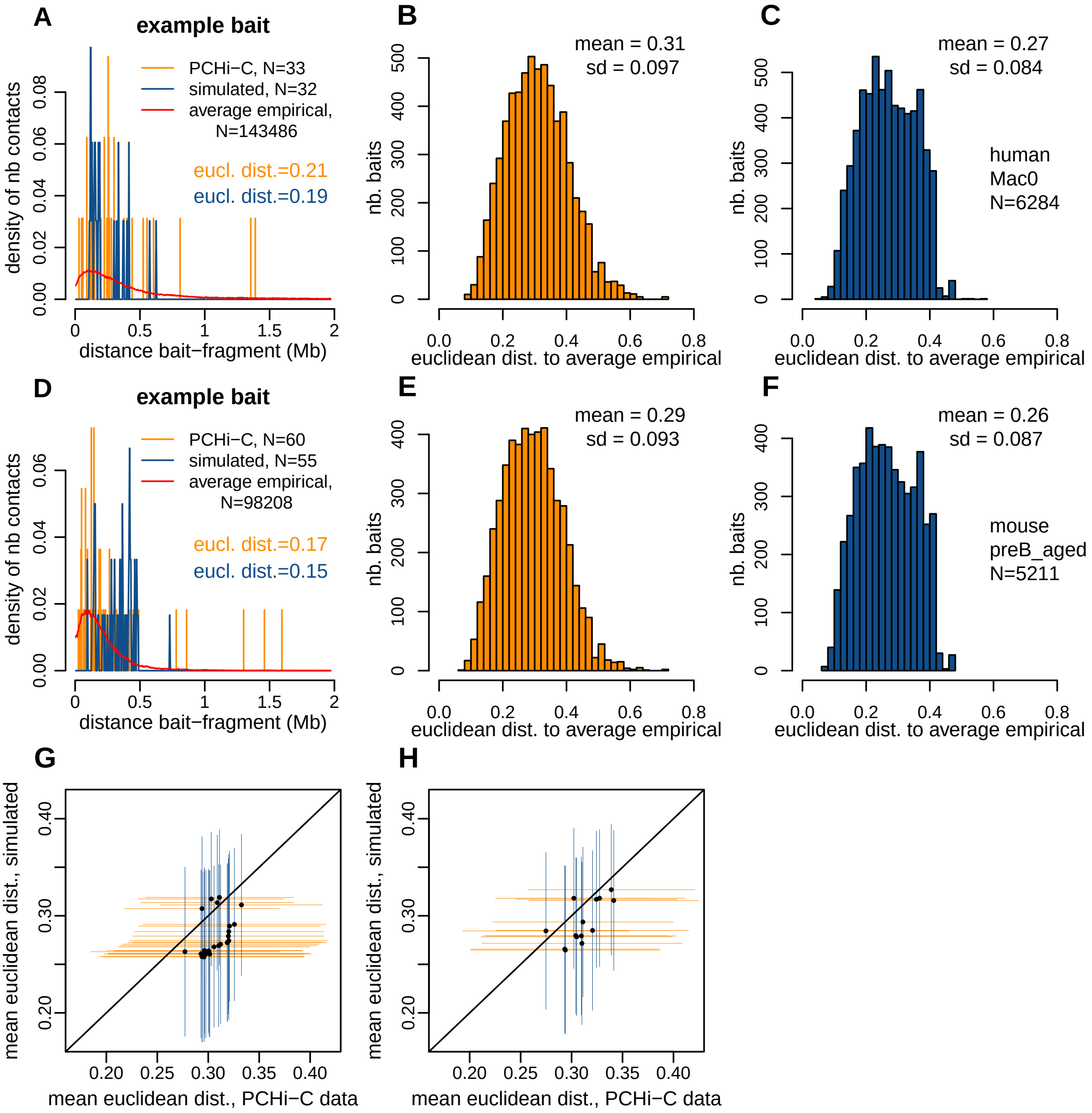

**Supplemental Text Fig. S1**: Variability among baits in the PCHi-C data and in the simulated data, in terms of distances to contacted fragments. **A)** Density plot representing the distribution of distances between an example bait and their non-baited contacted fragments for PCHi-C data (orange) and simulated data (blue), for one human sample (Mac0). The red line represents the distance distribution for all PCHi-C interactions in this sample (N=143,486), hereafter termed the average empirical distribution. The other N values displayed are the number of interactions for this baited fragment in PCHi-C and simulated data. We display the values of Euclidean distance between the average empirical distribution and the distance distribution of this baited fragment, in the PCHi-C and in the simulated data. **B)** Histogram representing the distribution of the Euclidean distances between the average empirical distribution and the distance distribution observed for each baited fragment in the PCHi-C data. Data shown is for the human Mac0 sample. **C)** Same as **A)**, for the simulated data. N represents the number of baits contacted in this sample. **D-F)** Same as **A-C)**, for the mouse preB_aged sample. **G)** Comparison between the mean Euclidean distance for baited restriction fragments in PCHi-C data (orange) and in simulated data (blue). Each dot represents one human sample. Horizontal lines represent the standard-deviation to the mean value for each PCHi-C sample. Vertical lines represent the standard-deviation to the mean value for each simulated sample. **H)** Same as **G)**, for mouse.

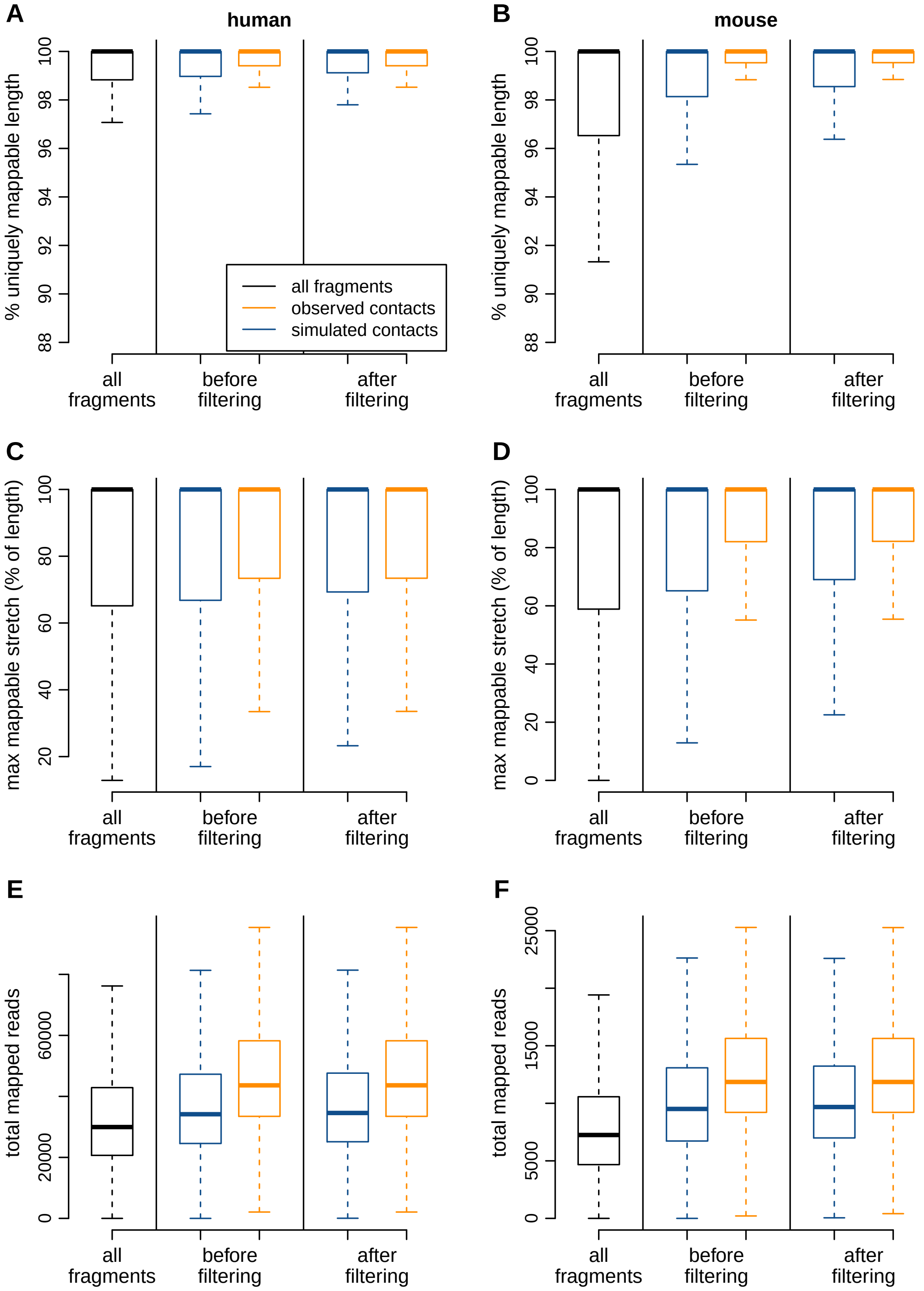

**Supplemental Text Fig. S2**: Theoretical mappability and number of mapped PCHi-C reads for restriction fragments. **A)** Percentage of restriction fragments length covered by uniquely mappable bases (Methods), for all human restriction fragments (black), restriction fragments contacted in human PCHi-C data (orange) and restriction fragments contacted in human simulated data (blue). **B)** Same as **A)**, for mouse restriction fragments. **C)** Distribution of the maximum mappable stretch (i.e the largest perfectly theoretical mappable interval), as a proportion of the length of human restriction fragments. **D)** Same as **C)**, for mouse restriction fragments. **E)** Distribution of the total number of PCHi-C reads mapped on retriction fragments, combined across all human samples. **F)** Same as **E)**, for mouse restriction fragments. **A-F)** Distributions are shown before and after filtering steps of reducing the discrepancies between the PCHi-C and the simulated dataset in term of mappability (Methods). Outlier points are not shown in the boxplots (outline=F in R).

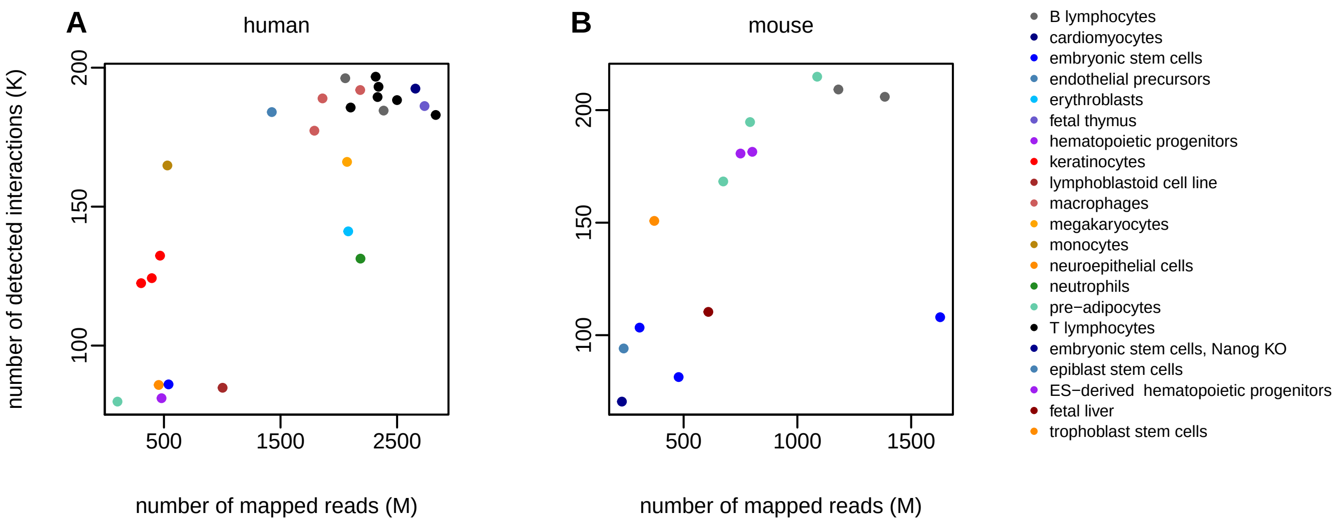

**Supplemental Text Fig. S3**: Sequencing depth is correlated with the number of detected PCHi-C interactions. **A)** Number of detected chromatin interactions as a function of the number of mapped reads in each human PCHi-C sample. Points color is assigned according to the cell type of the sample. **B)** Same as **A)**, for mouse PCHi-C samples.

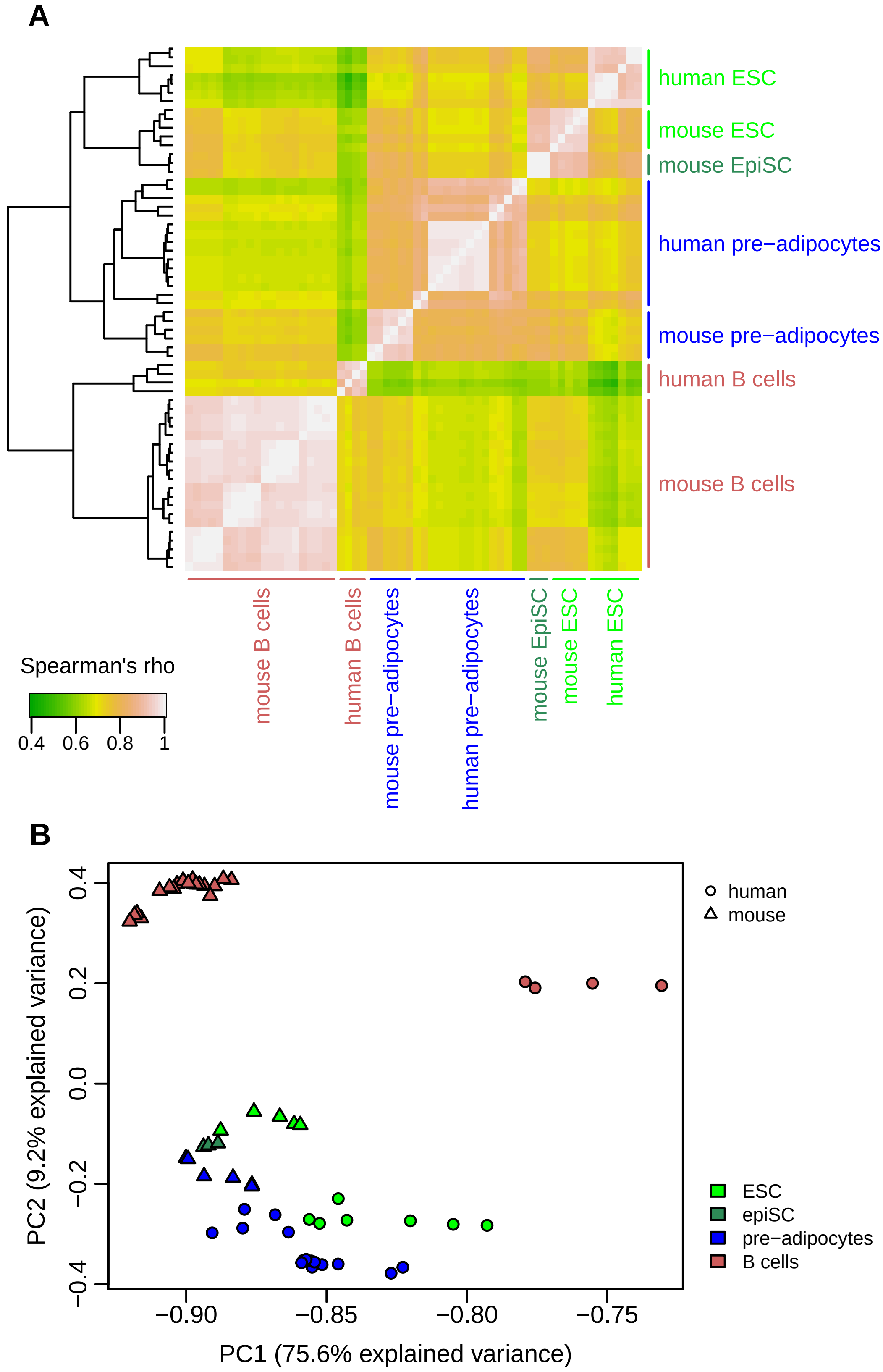

**Supplemental Text Fig. S4**: Gene expression levels are well conserved between human and mouse, for embryonic stem cells and epiblast-derived stem cells, pre-adipocytes and B lymphocytes. **A)** Hierarchical clustering performed on pairwise distances between samples. Distances are constructed based on Spearman's correlation coefficient (distance = 1-rho). **B)** Principal component analysis for human and mouse expression levels, for the same 4 cell types.

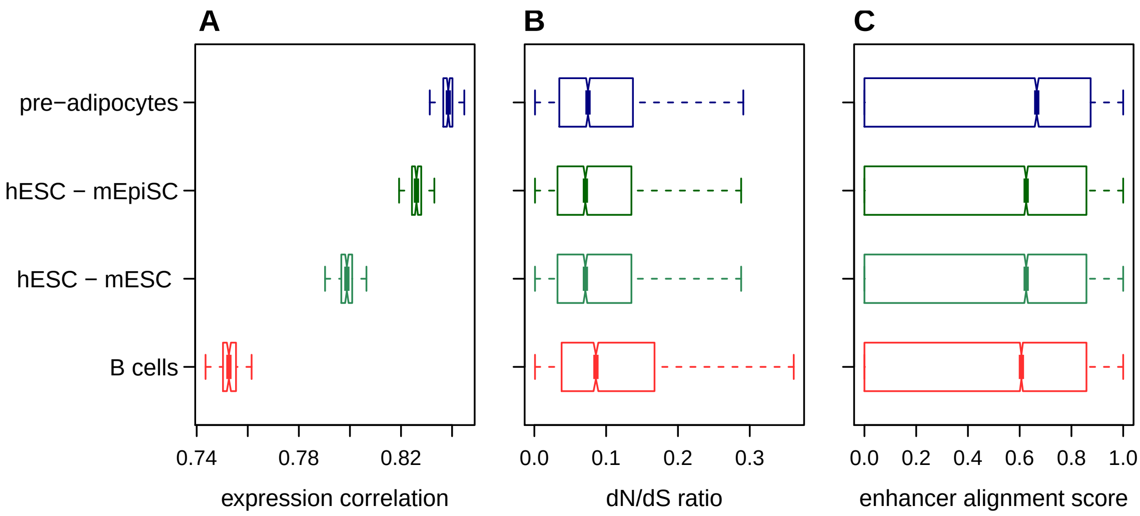

**Supplemental Text Fig. S5**: Parallel trends in gene expression evolution and *cis*-regulatory landscape evolution among cell types. **A)** Gene expression level correlation between human and mouse 1-to-1 orthologous genes in pre-adipocytes (blue), emybronic stem cells (green), B lymphocytes (red). The distribution is obtained from 100 bootstrap replicates. **B)** Protein-coding sequence evolution (dN/dS ratio) for the genes with the highest expression levels (top 25%) in each cell type. Only 1-to-1 orthologous genes between human and mouse were considered. **C)** Distribution of the average percentage of aligned nucleotides of human ENCODE enhancers contacted in PCHi-C samples of each cell type.

#### 3 Additional figures related to the main and supplemental figures

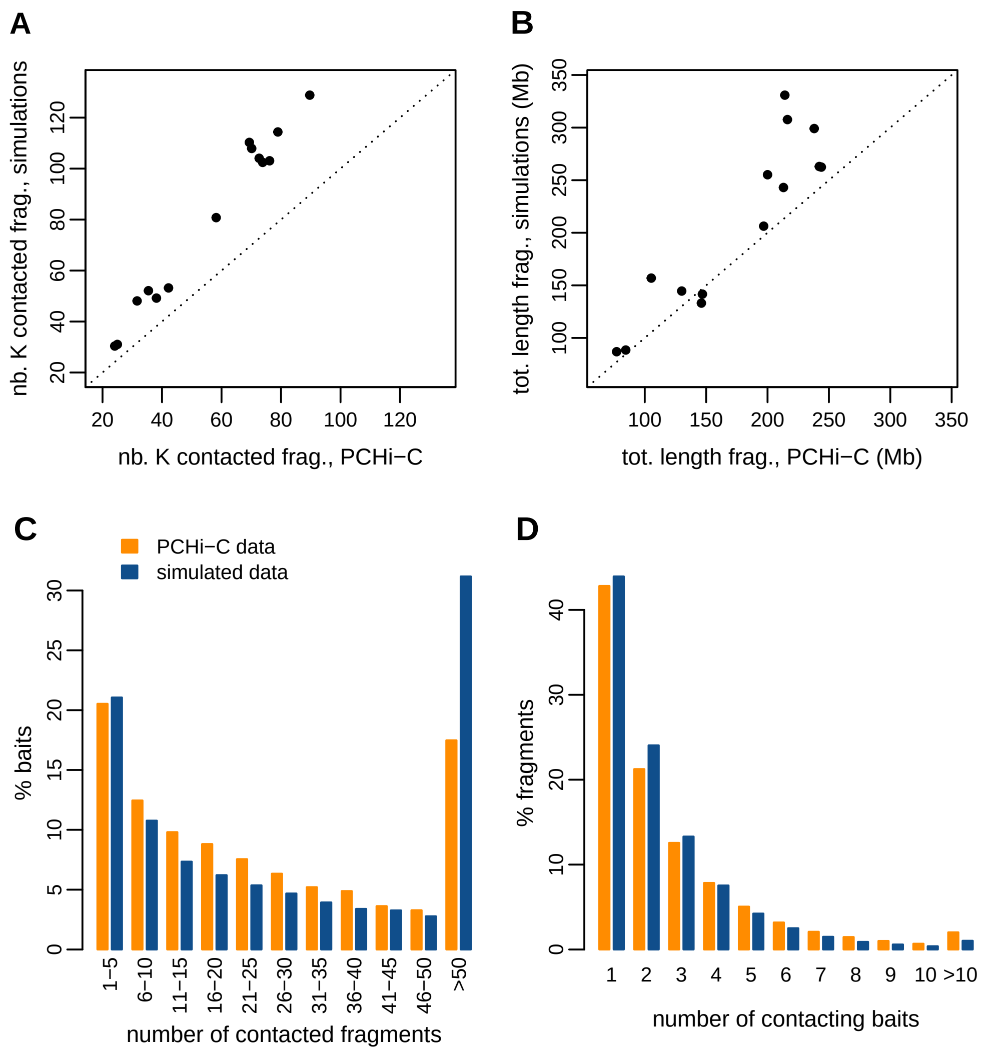

**Replication Fig. S1**: Number and total length of contacted restriction fragments, **mouse** PCHi-C and simulated data. Related to **Supplemental Fig. S2**. **A)** Comparison between the total number of contacted fragments in PCHi-C data and in simulated data. Each dot represents one mouse sample. **B)** Comparison between the total length covered by contacted fragments in PCHi-C data and in simulated data. Each dot represents one mouse sample. **C)** Histogram of the number of contacted restriction fragments *per* bait, for mouse PCHi-C data (orange) and simulated data (blue). **D)** Histogram of the number of contacting baits *per* restriction fragment, for mouse PCHi-C data (orange) and simulated data (blue). **C-D)** All mouse samples are combined in a single dataset for this analysis.

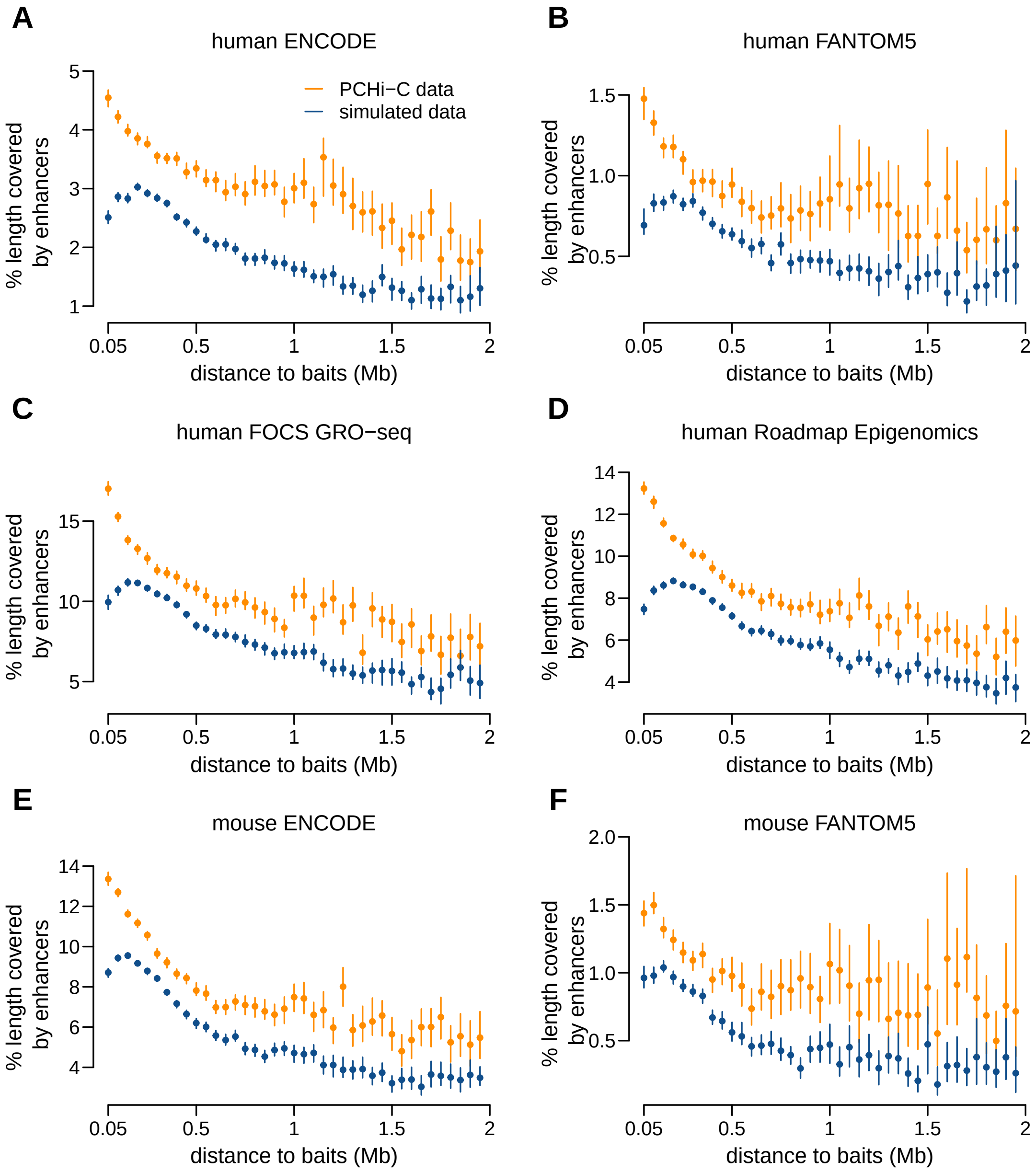

**Replication Fig. S2**: Enhancer density in contacted fragments decreases with the distance to contacting baits. Related to **Fig. 2**. **A-D)** Average length fraction of human restriction fragments that is covered by predicted enhancers, as a function of the median distance between the baited fragments and the contacted fragments, for PCHi-C data (orange) and simulated data (blue). The four panels represent the four enhancer datasets: ENCODE, FANTOM5, FOCS GRO-seq and RoadMapEpigenomics. **E-F)** Average length of mouse restriction fragments that is covered by predicted enhancers, as a function of the median distance between the baited fragments and the contacted fragments. The two panels represent the ENCODE and FANTOM5 enhancer datasets.

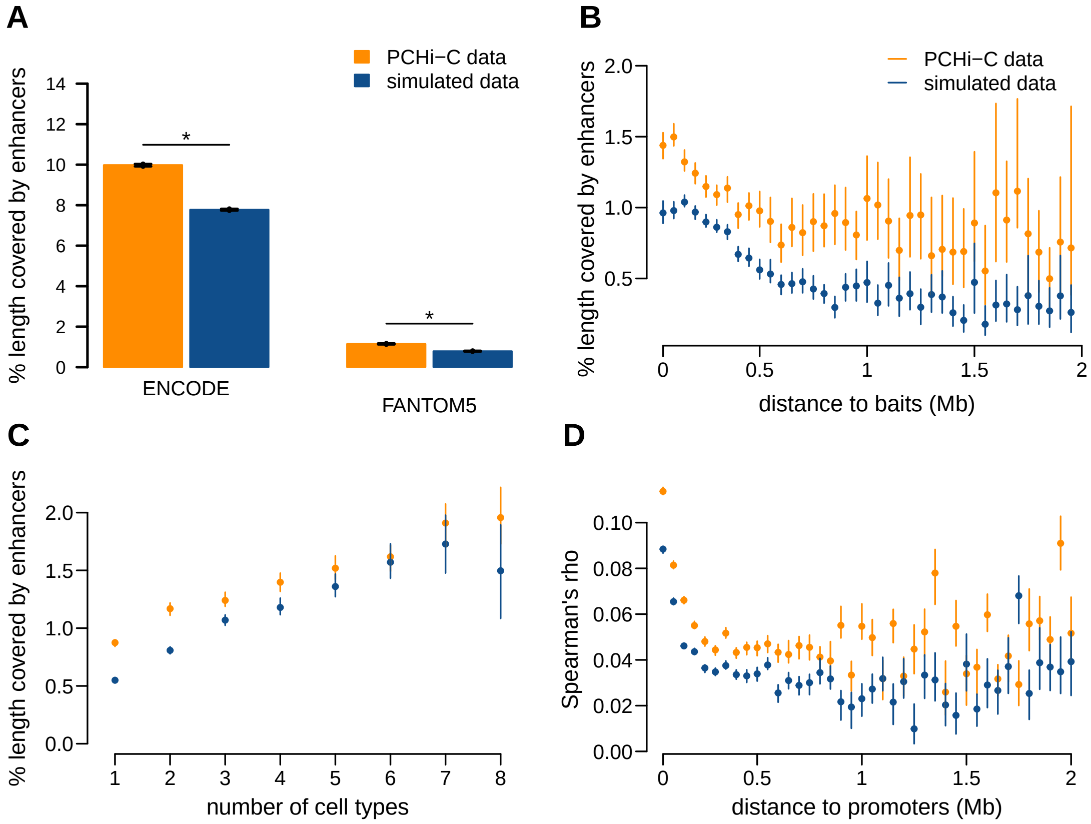

**Replication Fig. S3**: **Mouse** PCHi-C chromatin contacts are enriched in regulatory interactions. Related to **Fig. 2**. **A)** Average length fraction of mouse restriction fragments that is covered by predicted enhancers, for PCHi-C data (orange) and simulated data (blue). **B)** Average length fraction of mouse restriction fragments that is covered by FANTOM5 enhancers, as a function of the median distance between the baited fragments and the contacted fragments. **C)** Average length fraction of mouse restriction fragments that is covered by FANTOM5 enhancers, as a function of the number of cell types in which interactions are observed. **D)** Distribution of Spearman's correlation coefficients between gene expression levels and FANTOM5 enhancer activity. Pairs of promoters-enhancers are divided in classes based on the distance between them.

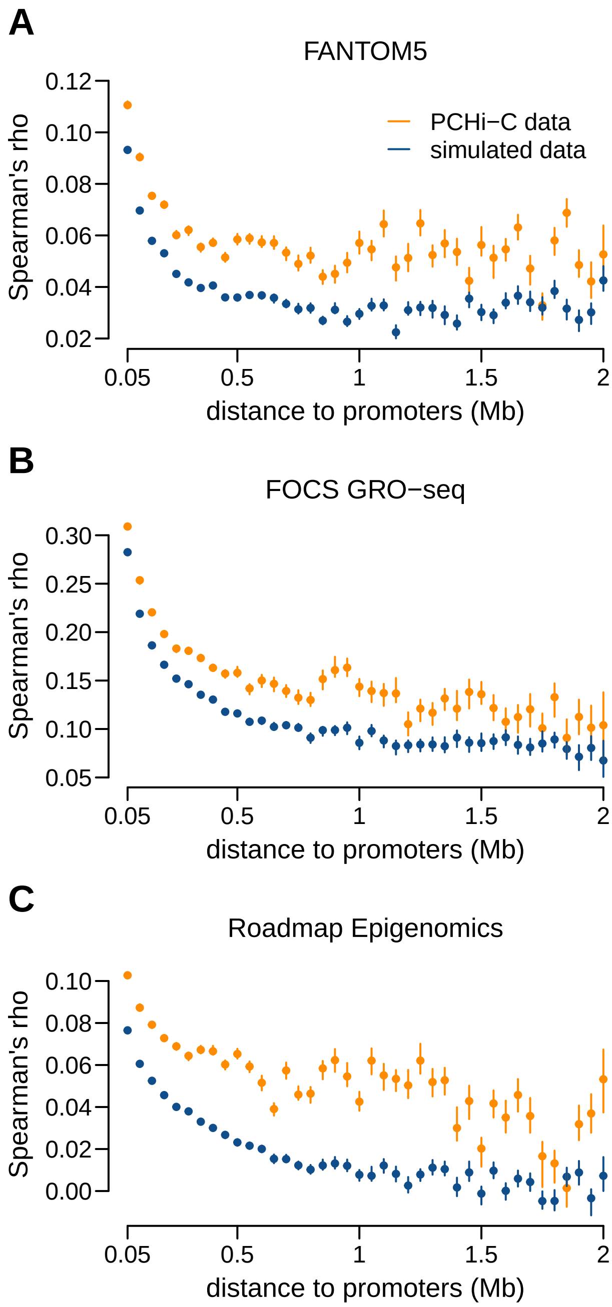

**Replication Fig. S4**: Distribution of Spearman's correlation coefficients between gene expression levels and predicted enhancer activity, for **human** PCHi-C data (orange) and simulated data (blue). Related to **Fig. 2**. Pairs of promoters-enhancers are divided in classes based on the distance between them. The panels represent different enhancer datasets: **A)** FANTOM5, **B)** FOCS GRO-seq, **C)** RoadMap Epigenomics.

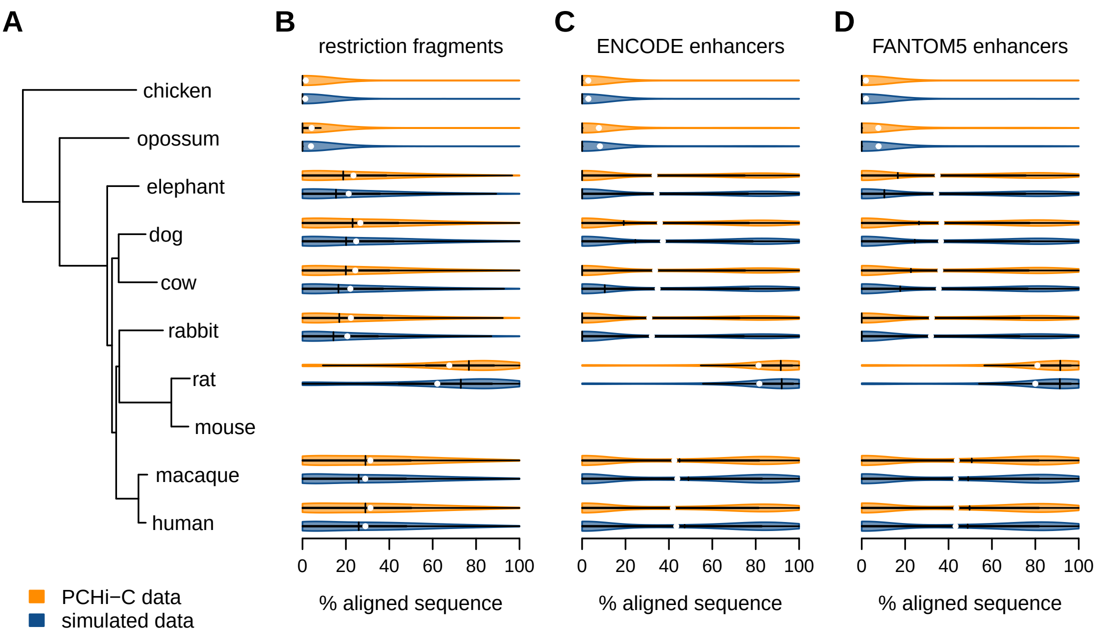

**Replication Fig. S5**: Genomic sequences contacted by **mouse** promoters are conserved during evolution. Related to **Fig. 3**. **A)** Species tree. **B)** Sequence conservation levels for contacted restriction fragments. The violin plots represent the distribution of the percentage of aligned nucleotides, for pairwise comparisons between mouse and other species. The vertical segments represent the median values of the distributions, the white dots represent the average values. **C)** Sequence conservation levels for contacted ENCODE enhancers. **D)** Sequence conservation levels for contacted FANTOM5 enhancers.

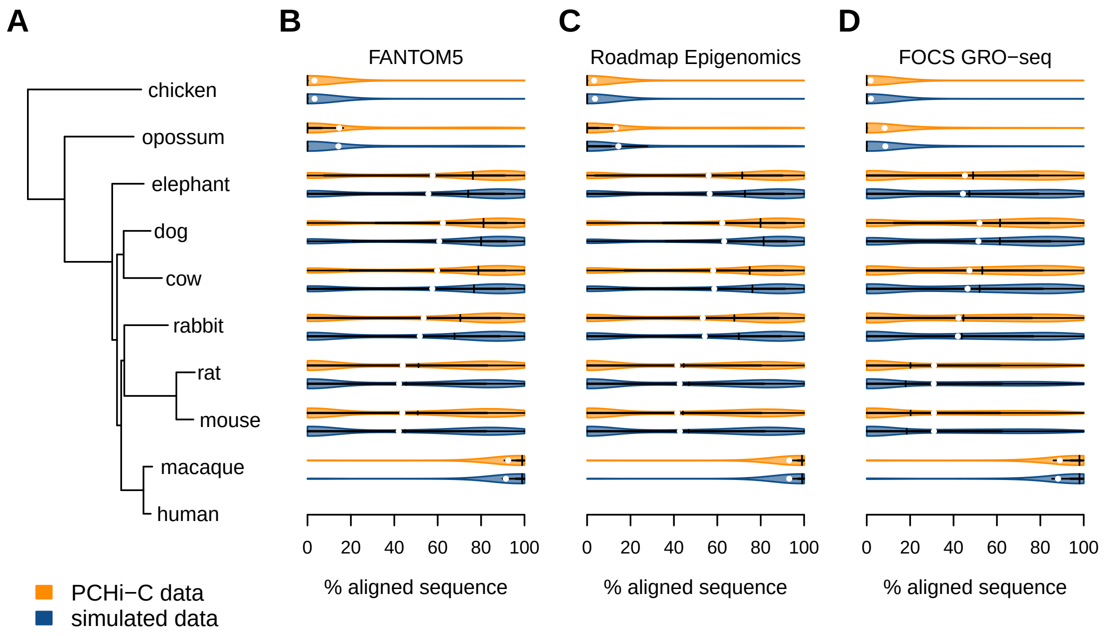

**Replication Fig. S6**: Genomic sequences contacted by **human** promoters are conserved during evolution. Related to **Fig. 3**. **A)** Species tree. **B)** Sequence conservation levels for contacted FANTOM5 enhancers. The violin plots represent the distribution of the percentage of aligned nucleotides, for pairwise comparisons between human and other species. **C)** Sequence conservation levels for contacted Roadmap Epigenomics enhancers. **D)** Sequence conservation levels for contacted FOCS GRO-seq enhancers.

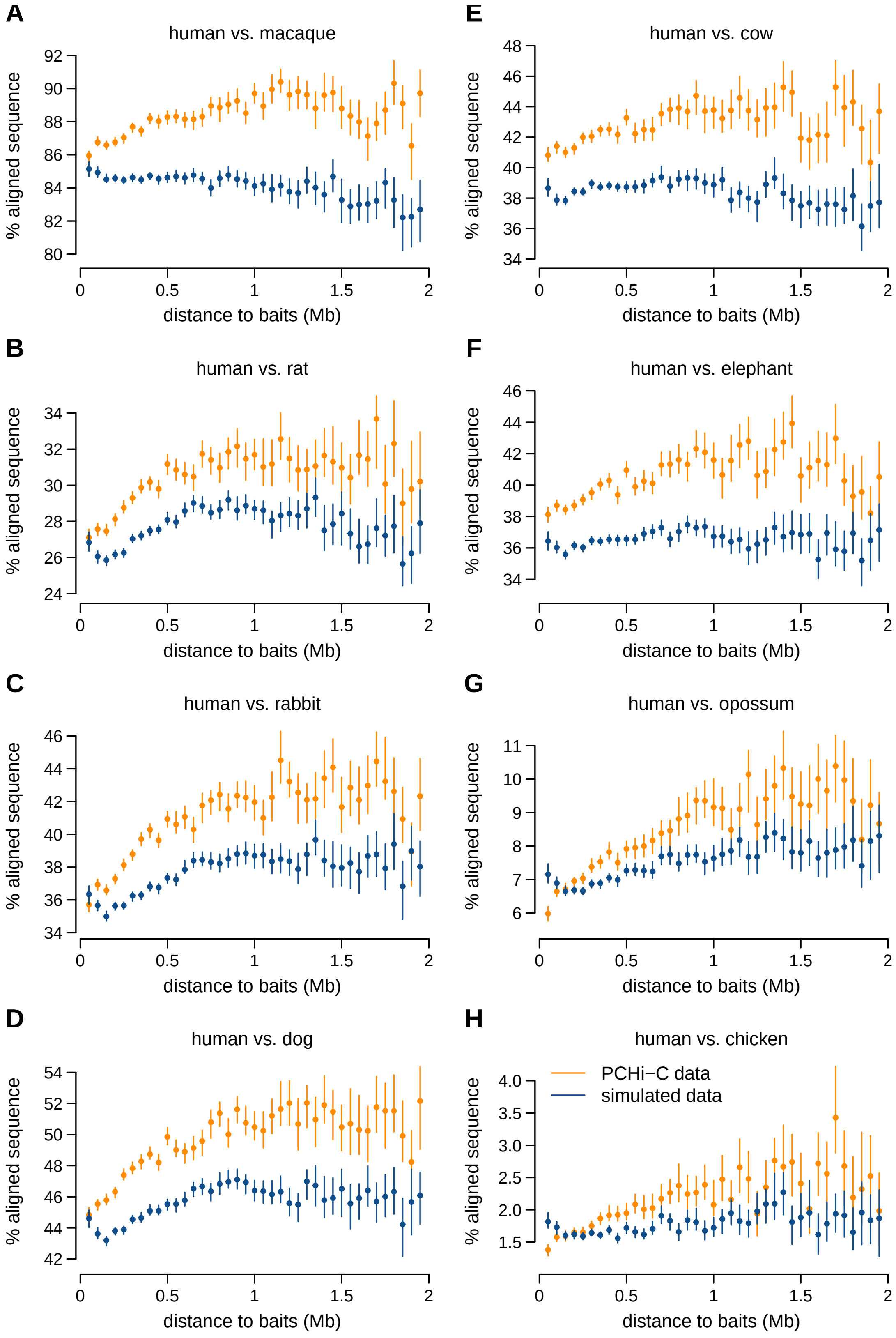

**Replication Fig. S7**: Conservation of **human** contacted restriction fragments as a function of the median genomic distance between the baited fragments and the contacted fragments, for PCHi-C data (orange) and simulated data (blue). Related to **Fig. 3**. We show pairwise comparisons between human and other species: **A)** macaque, **B)** rat, **C** rabbit, **D)** dog, **E)** cow, **F)** elephant, **G)** opossum, **H)** chicken.

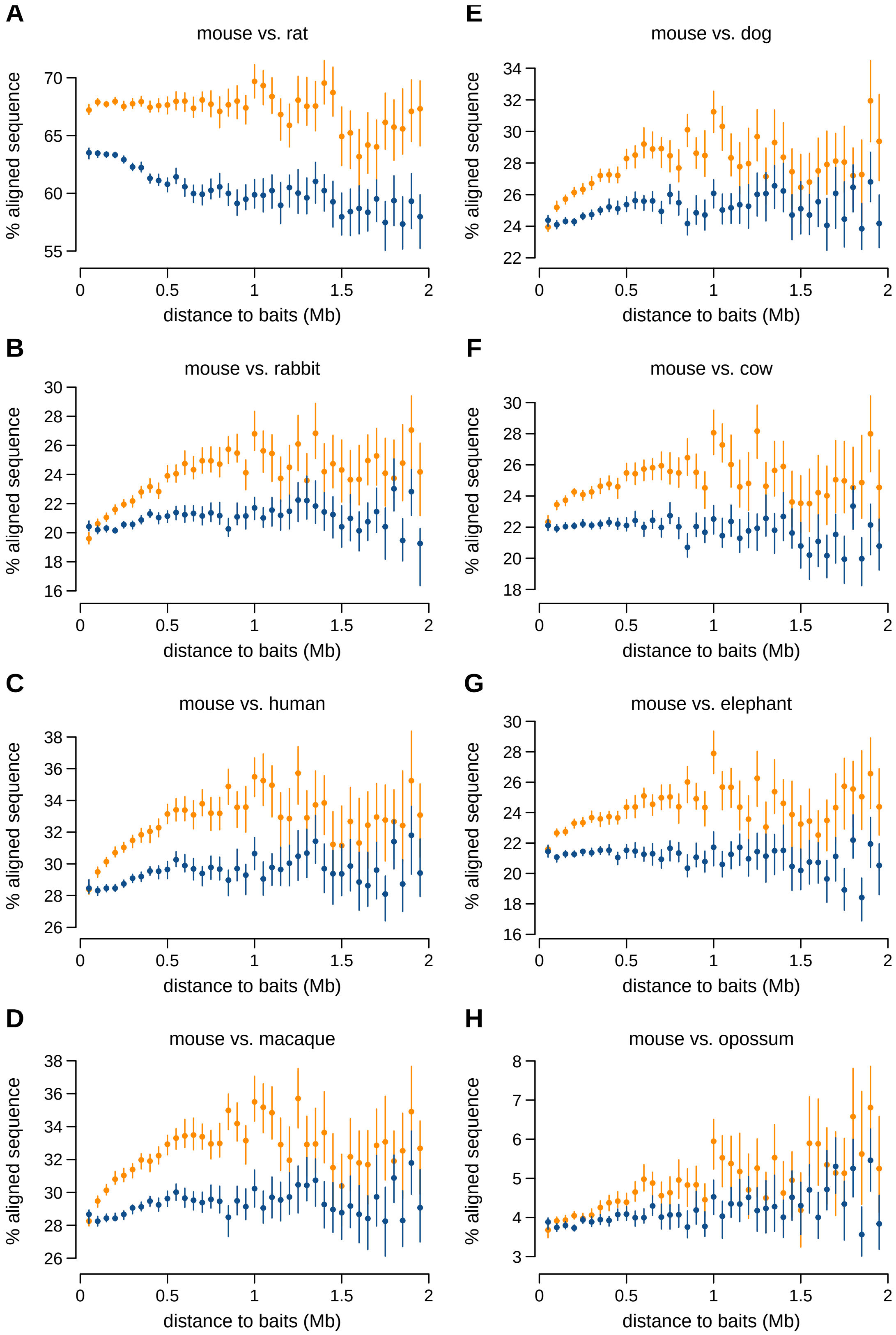

**Replication Fig. S8**: Conservation of **mouse** contacted restriction fragments as a function of the median genomic distance between the baited fragments and the contacted fragments, for PCHi-C data (orange) and simulated data (blue). Related to **Fig. 3**. We show pairwise comparisons between mouse and other species: **A)** rat, **B)** rabbit, **C)** human, **D)** macaque, **E)** dog, **F)** cow, **G)** elephant, **H)** opossum.

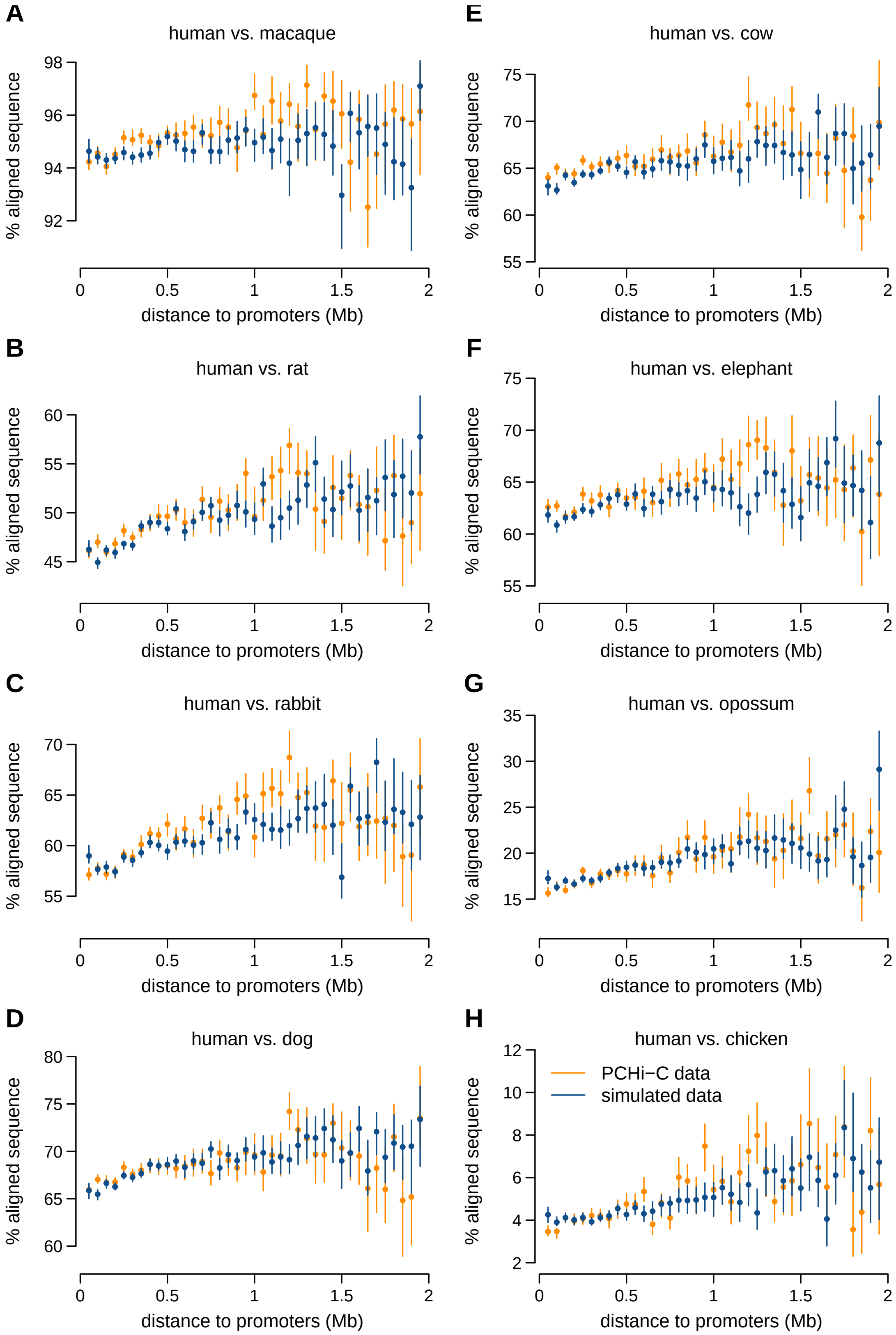

**Replication Fig. S9**: Conservation of **human** contacted ENCODE enhancers as a function of the median genomic distance between promoters and enhancers, for PCHi-C data (orange) and simulated data (blue). Related to **Fig. 3**. We show pairwise comparisons between human and other species: **A)** macaque, **B)** rat, **C** rabbit, **D)** dog, **E)** cow, **F)** elephant, **G)** opossum, **H)** chicken.

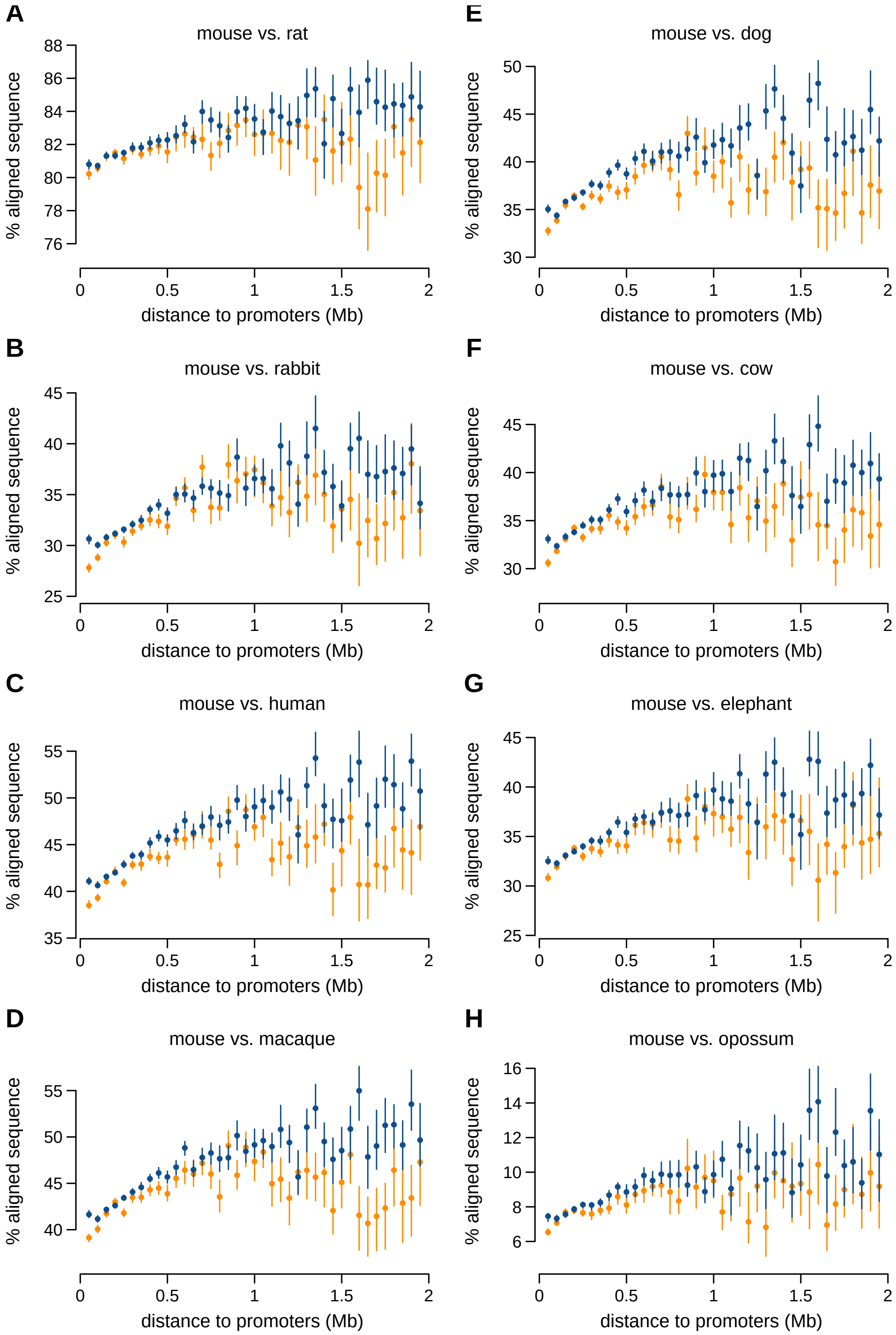

**Replication Fig. S10**: Conservation of **mouse** contacted ENCODE enhancers as a function of the median genomic distance between promoters and enhancers, for PCHi-C data (orange) and simulated data (blue). Related to **Fig. 3**. We show pairwise comparisons between mouse and other species: **A)** macaque, **B)** rat, **C** rabbit, **D)** dog, **E)** cow, **F)** elephant, **G)** opossum, **H)** chicken.

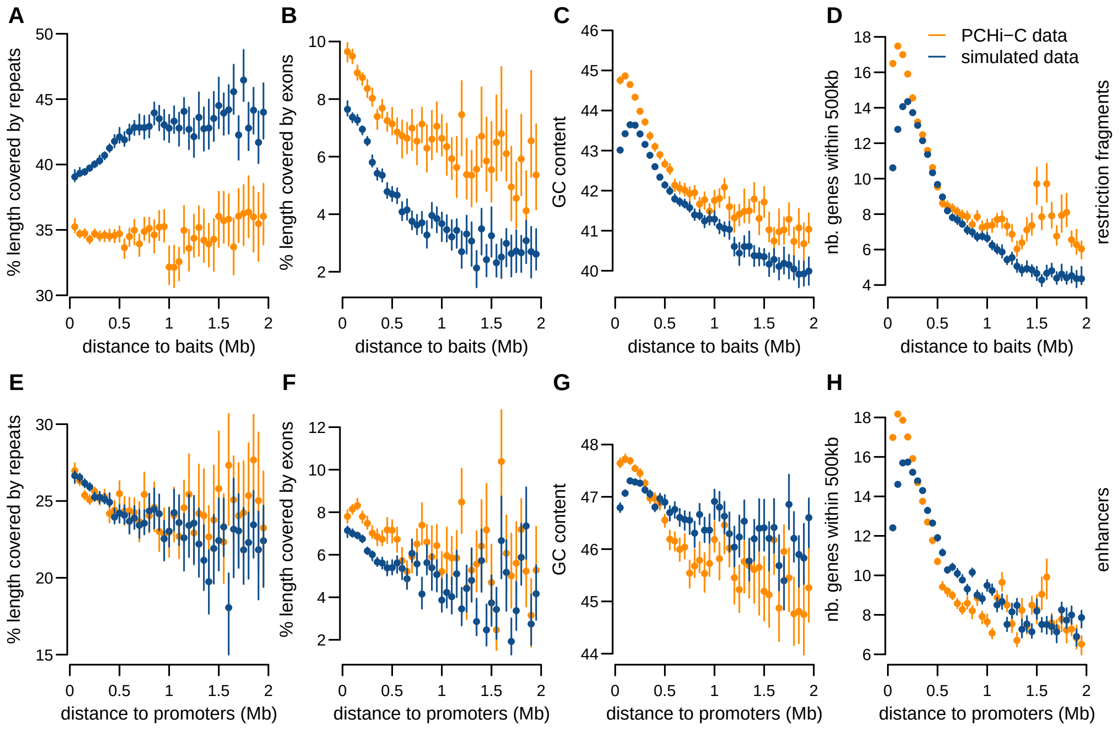

**Replication Fig. S11**: Genomic characteristics of **mouse** contacted sequences according to distance from promoters. Related to **Supplemental Fig. S8**. From left to right: Average percentage of length covered by repeated elements, average percentage of length covered by exons, GC content, number of genes within a maximum distance of 500kb. **A-D)** Genomic characteristics of contacted restriction fragments for PCHi-C data (orange) and simulated data (blue). **E-H)** Genomic characteristics of contacted ENCODE enhancers.

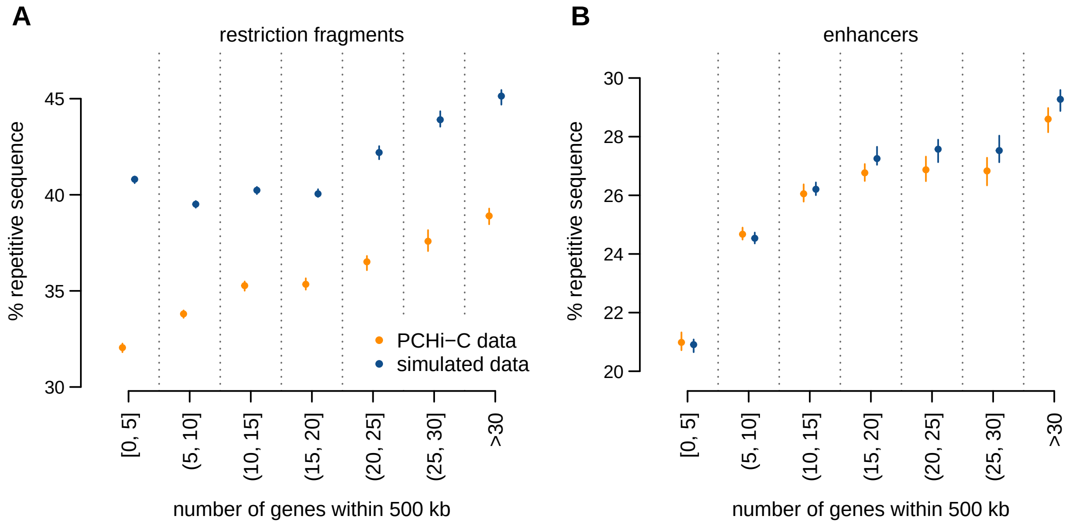

**Replication Fig. S12**: Gene density is correlated with repetitive sequence content for contacted restriction fragments and enhancers, in **mouse**. Related to **Supplemental Fig. S11**.

**A)** Average fraction of restriction fragment length that is covered by repetitive sequences, as a function of the number of genes found at most 500kb away from the restriction fragment.

**B)** Same as **A)**, for ENCODE enhancers.

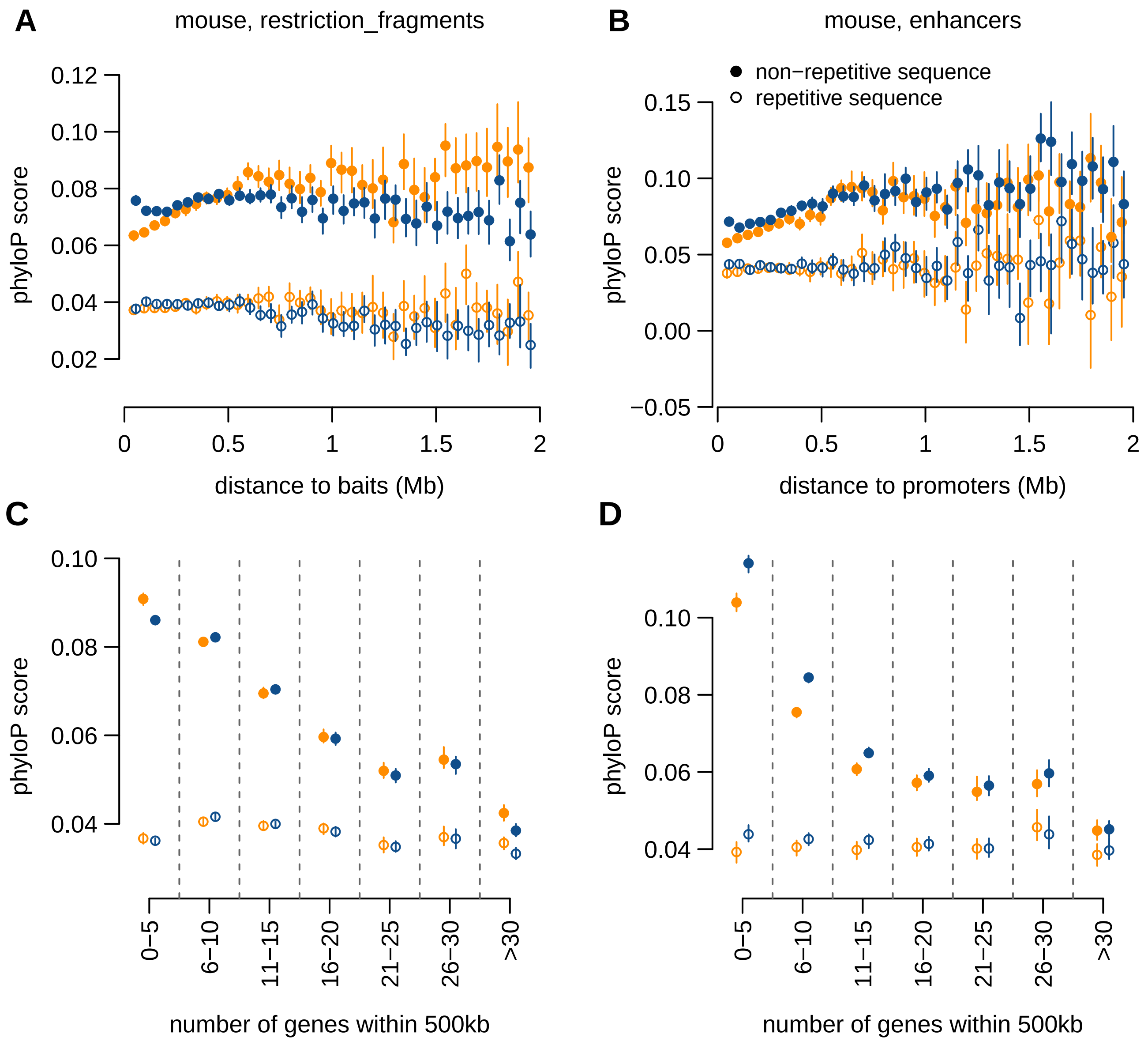

**Replication Fig. S13**: Sequence conservation measured by phyloP score for **mouse** restriction fragments and ENCODE enhancers. Related to **Fig. 3**. **A)** Average phyloP score of contacted restriction fragments in PCHi-C (orange) and simulated (blue) data, as a function of the median genomic distance between restriction fragments and contacting baits. **B)** Same as **A)**, for ENCODE enhancers. **C)** Average phyloP score of contacted restriction fragments, as a function of the number of genes found within at most 500kb from the restriction fragment. **D)** Same as **C)**, for ENCODE enhancers. **A-D)** Filled dots represent non-repetitive part of sequences; empty dots represent the repetitive part. Dots represent mean values, vertical segments represent 95% confidence intervals of the mean, obtained with a non-parametric bootstrap approach (Methods).

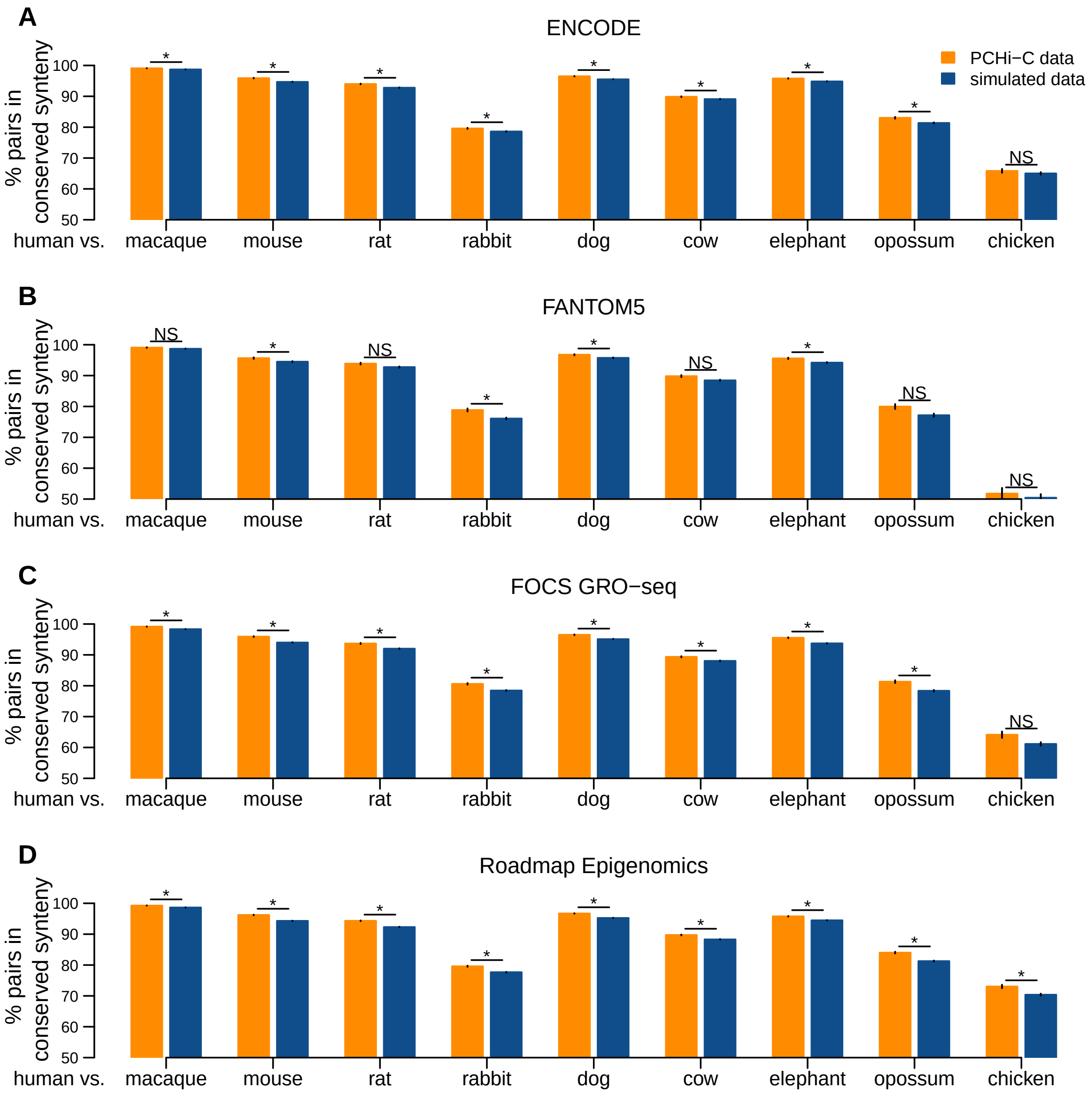

**Replication Fig. S14**: Pairs of **human** promoters and enhancers involved in chromatin contacts are maintained in synteny during evolution. Related to **Fig. 4**.The barplots represent the percentage of human promoter-enhancers pairs maintained in synteny in other vertebrate genomes, for PCHi-C data (orange) and simulated data (blue). **A)** ENCODE enhancers. **B)** FANTOM5 enhancers. **C)** FOCS GRO-seq enhancers. **D)** Roadmap Epigenomics enhancers. **A**-**D)** "*" indicates a significant difference between PCHi-C and simulated data (p-value lower than 1e-10 FDR) based on a chi-squared test.

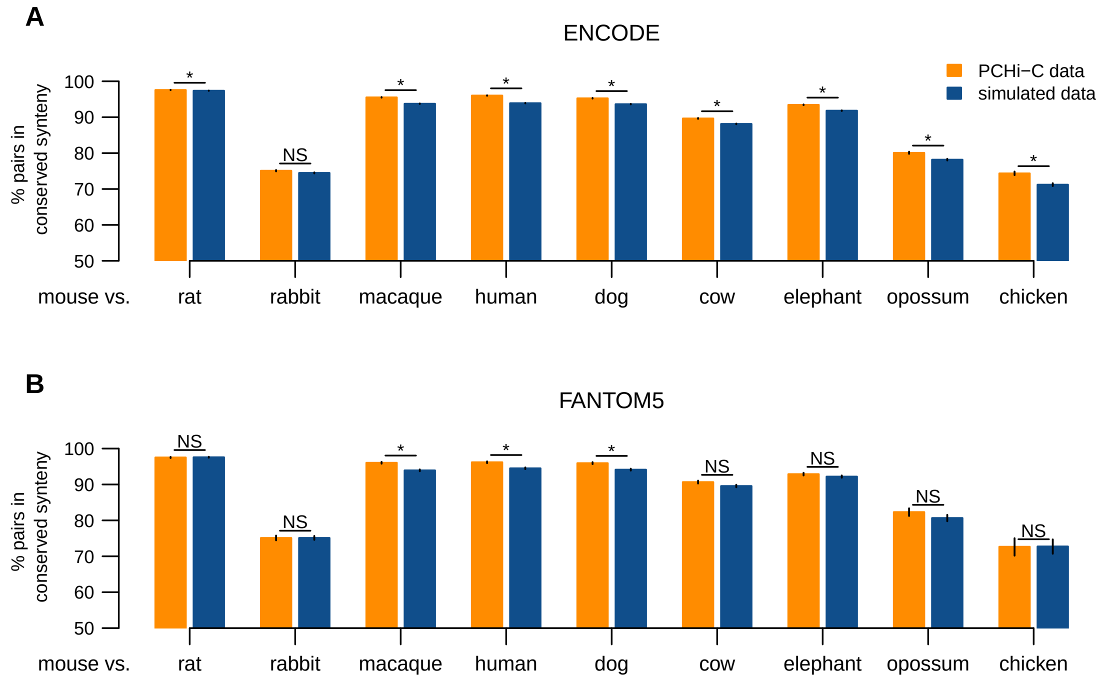

**Replication Fig. S15**: Pairs of **mouse** promoters and enhancers involved in chromatin contacts are maintained in synteny during evolution. Related to **Fig. 4**. The barplots represent the percentage of human promoter-enhancers pairs maintained in synteny in other vertebrate genomes, for PCHi-C data (orange) and simulated data (blue). **A)** ENCODE enhancers. **B)** FANTOM5 enhancers. **A-D)** "*" indicates a significant difference between PCHi-C and simulated data (p-value lower than 1e-10 FDR) based on a chi-squared test.

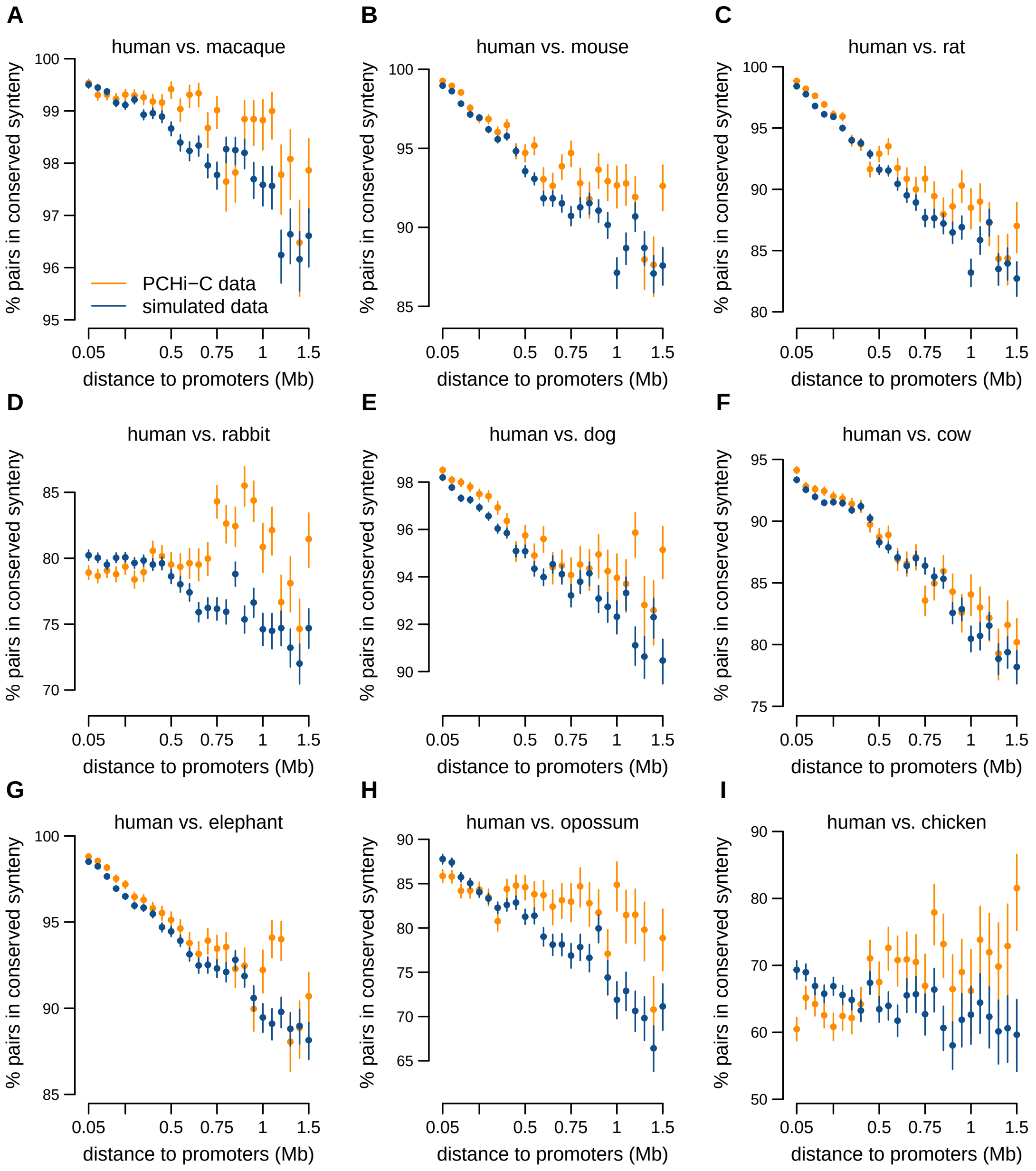

**Replication Fig. S16**: Average proportion of **human** promoter-enhancers pairs maintained in synteny in other vertebrate genomes, as a function of the distance between them in the human genome, for PCHi-C data (orange) and simulated data (blue). Related to **Fig. 4**. We show pairwise comparisons between human and other species: **A)** macaque, **B)** mouse, **C)** rat, **D)** rabbit, **E)** dog, **F)** cow, **G)** elephant, **H)** opossum, **I)** chicken.

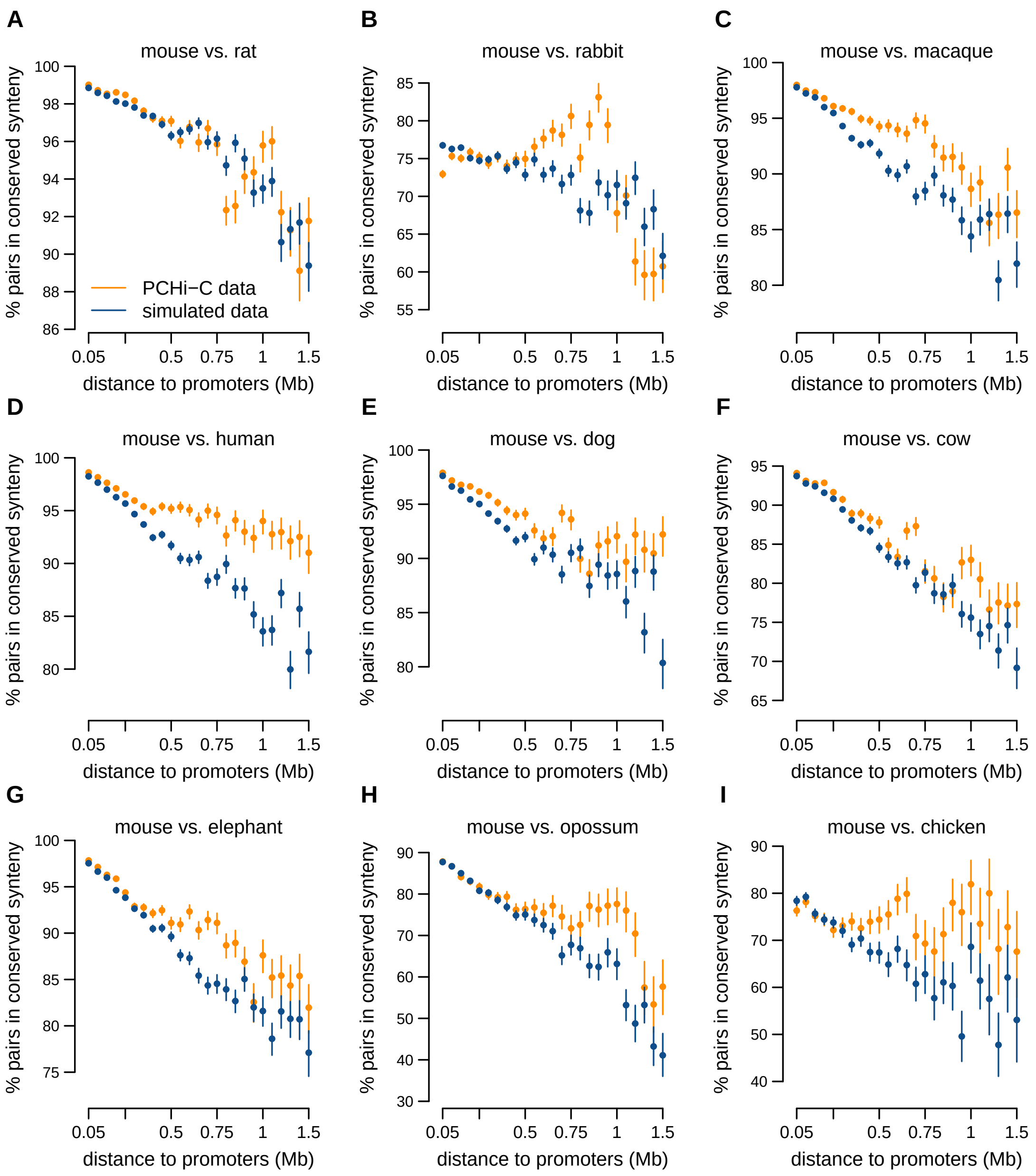

**Replication Fig. S17**: Average proportion of **mouse** promoter-enhancers pairs maintained in synteny in other vertebrate genomes, as a function of the distance between them in the mouse genome, for PCHi-C data (orange) and simulated data (blue). Related to **Fig. 4**. We show pairwise comparisons between mouse and other species: **A)** rat, **B)** rabbit, **C)** macaque, **D)** human, **E)** dog, **F)** cow, **G)** elephant, **H)** opossum, **I)** chicken.

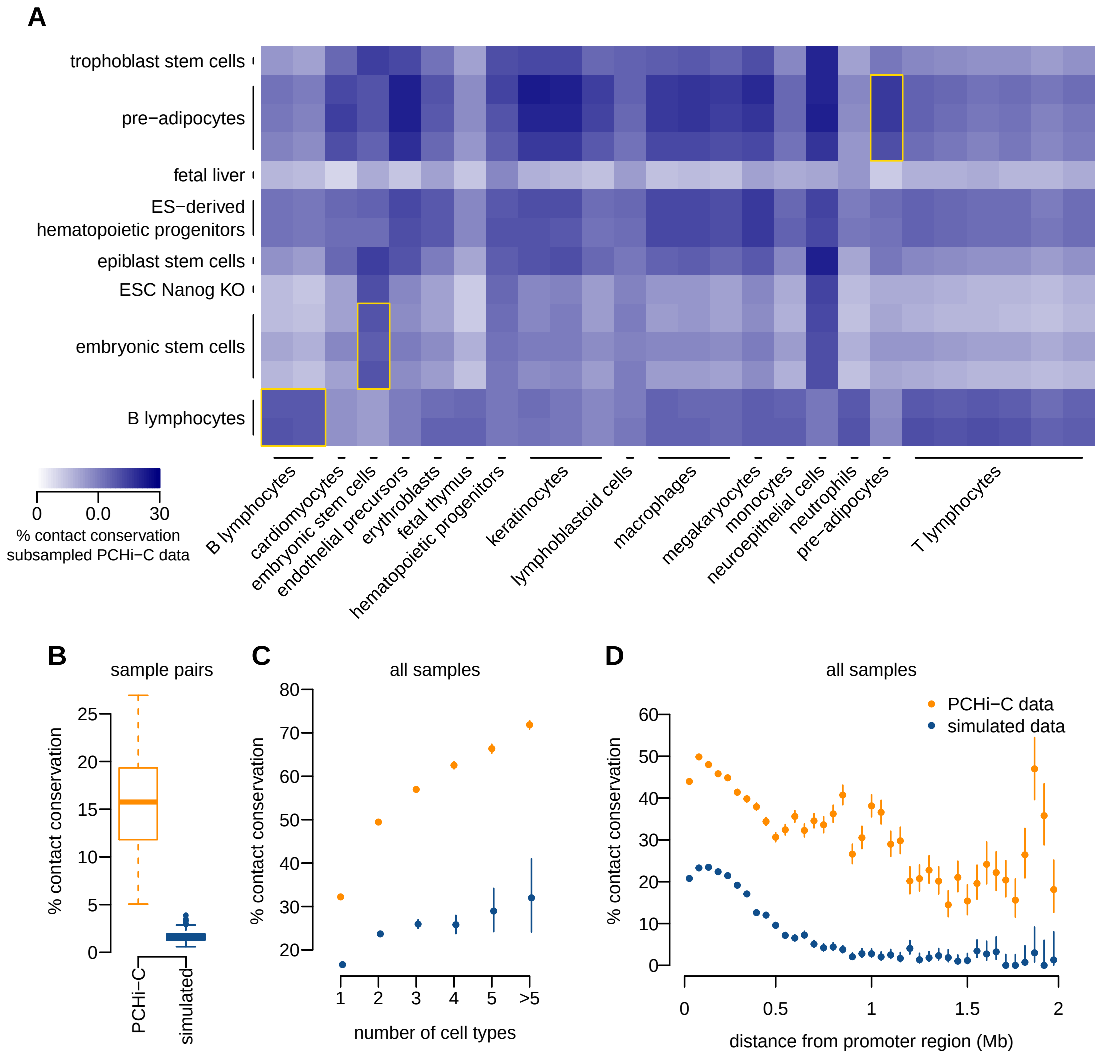

**Replication Fig. S18**: **Mouse** promoter-enhancer contact maps are conserved during evolution. Related to **Fig. 5**. **A)** Heatmap of the percentage of mouse promoter-enhancer contacts conserved between pairs of mouse and human samples. The values represent the difference between in PCHi-C data and simulated data. Yellow squares highlight cell types sampled for both species (embryonic stem cells, B lymphocytes and pre-adipocytes). **B)** Average proportion of mouse promoter-enhancer contacts conserved in human for PCHi-C data (orange) and simulated data (blue), as a function of the number of cell types in which interactions are observed. **C)** Average proportion of mouse promoter-enhancer contacts conserved in human, as a function of the distance between them in the mouse genome.

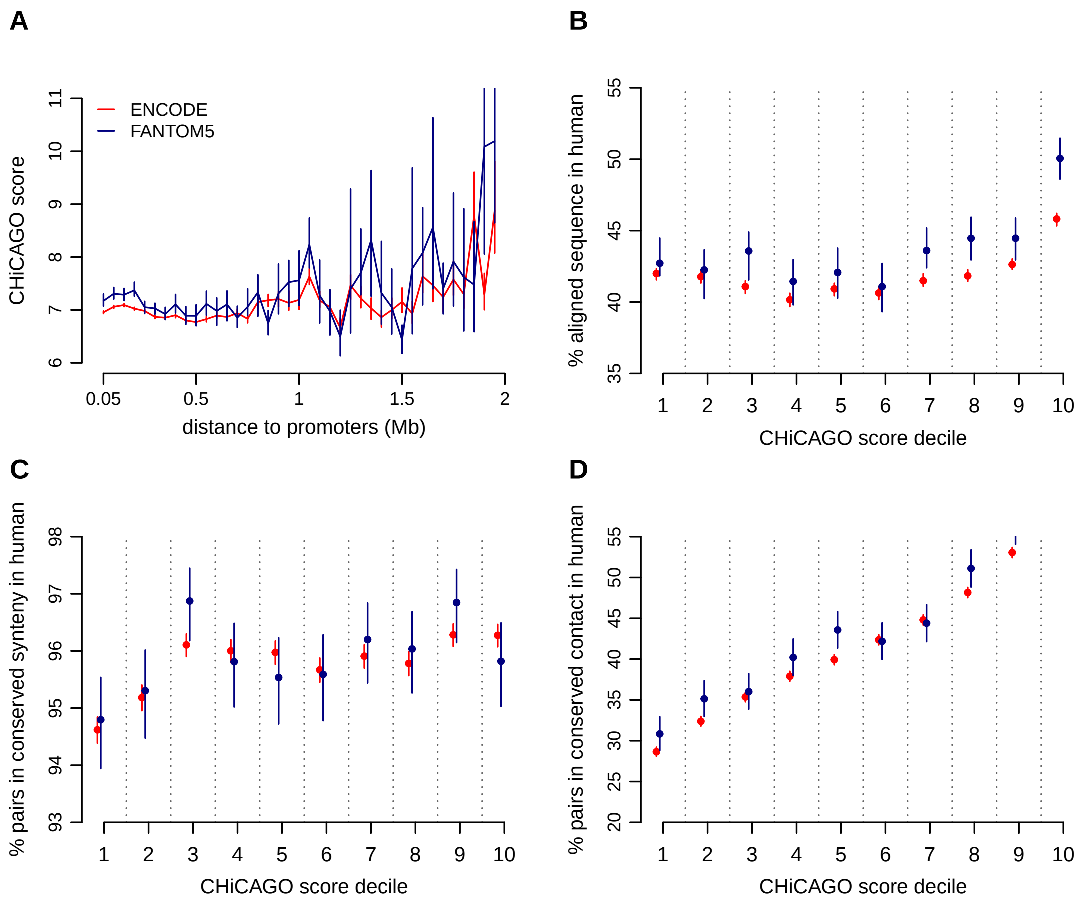

**Replication Fig. S19**: The CHiCAGO score, which measures the statistical significance of promoter-chromatin contacts, is correlated with the conservation of **mouse** *cis*-regulatory landscapes. Related to **Supplemental Fig. S14**. **A)** Average CHiCAGO score as a function of the distance between promoters and predicted enhancers in the genome, for ENCODE (red) and FANTOM5 enhancers (blue). **B)** Sequence conservation of mouse contacted enhancers, calculated as the percentage of aligned nucleotides between mouse and human, as a function of the CHiCAGO score decile. **C)** Proportion of mouse promoter-enhancer pairs maintained in synteny in the human genome, as a function of the CHiCAGO score decile. **D)** Proportion of mouse promoter-enhancer contacts that are conserved in human PCHi-C data, as a function of the CHiCAGO score decile.

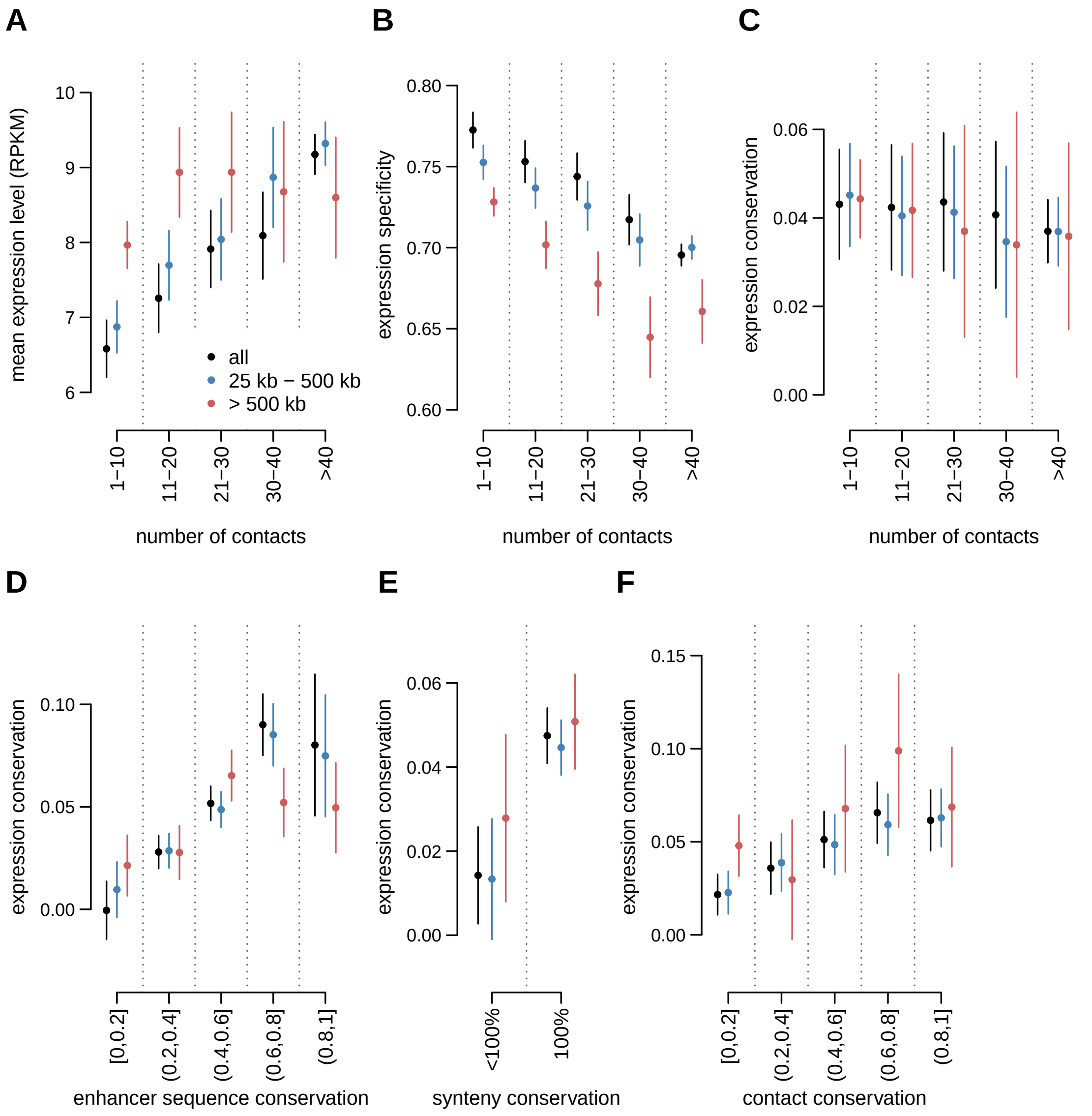

**Replication Fig. S20**: *Cis*-regulatory landscapes co-evolve with gene expression profiles, for **mouse**. Related to **Supplemental Fig. 21**. Gene expression conservation is measured with Spearman's correlation coefficient between human and mouse relative expression profiles, for pairs of 1-to-1 orthologous genes, across organs and developmental stages (expression data from Cardoso-Moreira et al., 2019). Expression conservation is further corrected to account for the effect of expression levels and of expression specificity with a multiple linear regression model (Methods). Enhancer predictions are taken from ENCODE. **A)** Average expression levels as a function of the number of contacted enhancers in mouse PCHi-C data. Promoter-enhancer contacts are divided into two classes according to the distance between them: medium range (25kb-500kb) in blue, long range (above 500Kb) in red. The black dots represent the full set of promoter-enhancer contacts. **B)** Mouse gene expression specificity as a function of the number of contacted enhancers. **C)** Expression conservation as a function of the number of contacted enhancers in mouse PCHi-C data. **D)** Expression conservation as a function of the average sequence conservation of mouse contacted ENCODE enhancers. **E)** Expression conservation, depending on whether or not genes underwent least one break of synteny with the contacted enhancers between human and mouse genomes. **F)** Expression conservation as a function of the proportion of mouse promoter-enhancers contacts conserved in mouse PCHi-C data. **A-F)** Dots represent median values across all genes in a class, vertical segments represents 95% confidence intervals for the median.
