## Supplemental Figures for "Long-range promoter-enhancer contacts are conserved during evolution and contribute to gene expression robustness"

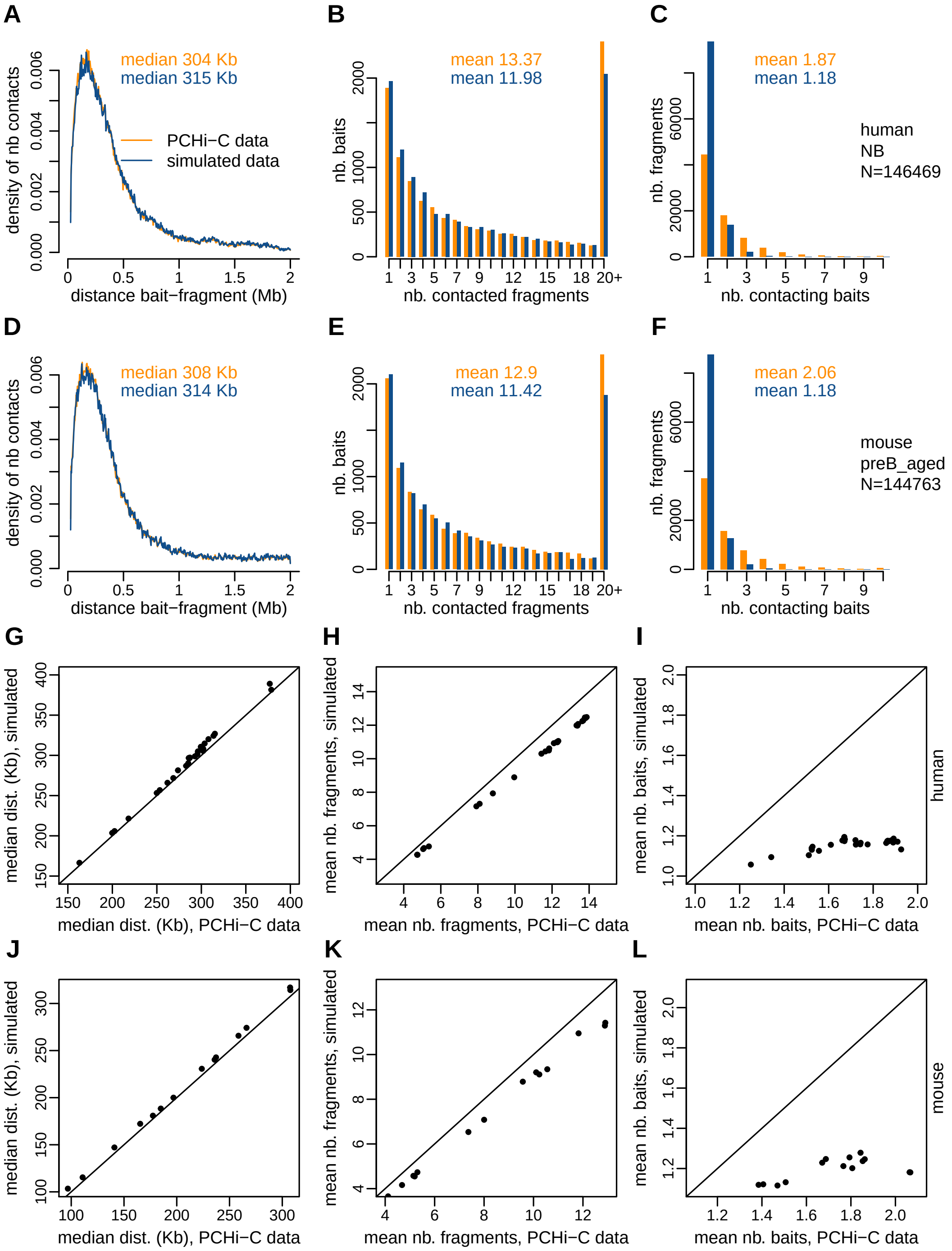


**Supplemental Fig. S1**: **A-C)** Characteristics of PCHi-C and simulated interaction data, for 1 human sample (identifier NB). **A.** Density plot representing the distribution of the distances between baited fragments and contacted fragments, after filtering steps (Methods). **B.** Histogram representing the distribution of the number of contacted fragments *per* bait, after filtering. **C.** Histogram representing the distribution of the number of contacting baits *per* fragment, after filtering. **D-F)** Same as **A-C**, for 1 mouse sample (identifier preB_aged). **G.** Scatterplot representing the median distance between baited fragments and contacted fragments, for the PCHi-C data (X-axis) and for the simulated data (Y-axis), for all human samples. The median distances were computed after filtering steps (Methods). **H.** Scatterplot representing the mean number of contacted fragments *per* bait, for the PCHi-C data (X-axis) and for the simulated data (Y-axis), for all human samples. **I.** Scatterplot representing the mean number of contacting baits *per* contacted restriction fragment, for the PCHi-C data (X-axis) and for the simulated data (Y-axis), for all human samples. **J.** Same as **G**, for mouse samples. **K.** Same as **H**, for mouse samples. **L.** Same as **I**, for mouse samples.

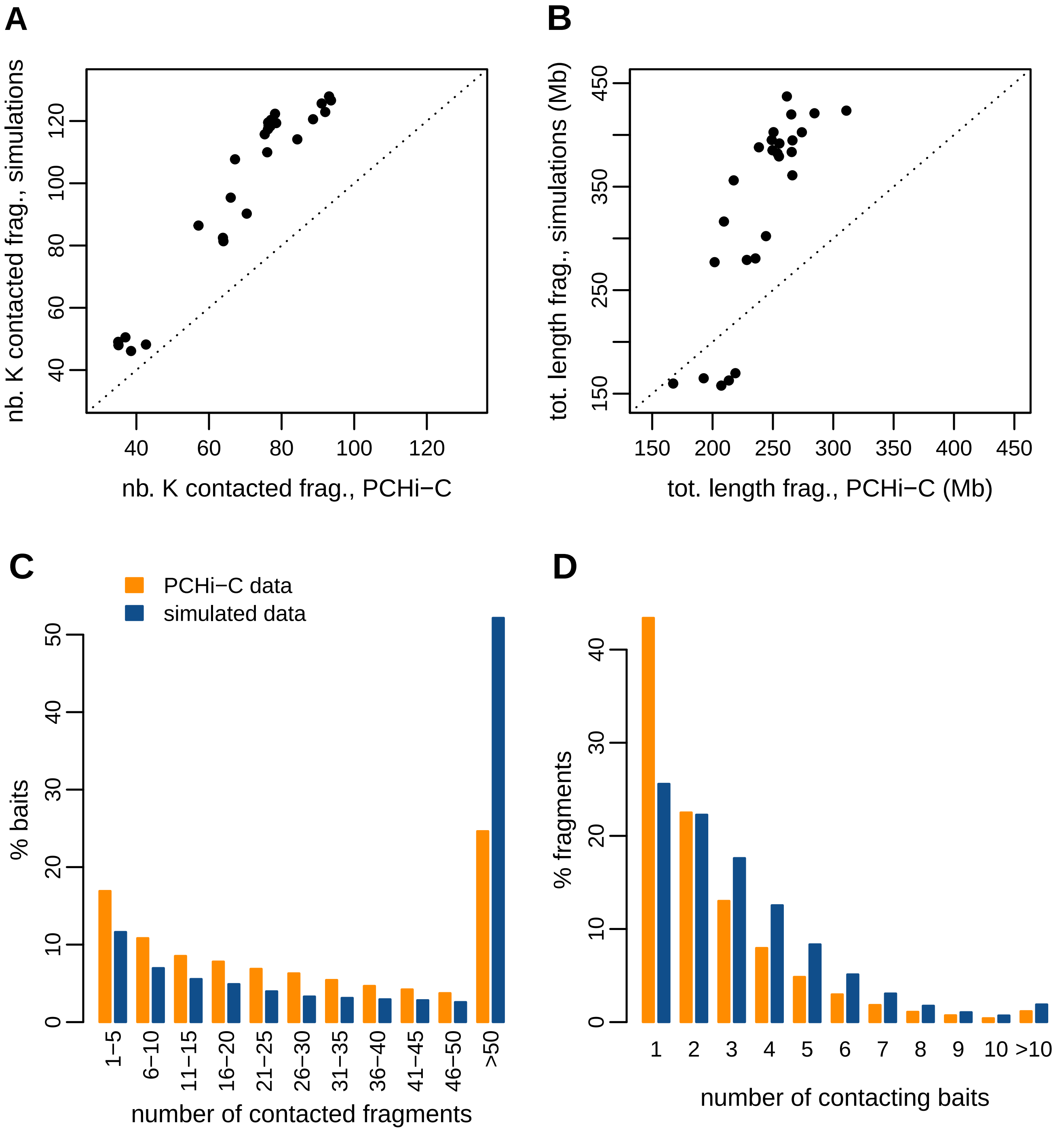


**Supplemental Fig. S2**: Characteristics of **human** PCHi-C and simulated data. **A.** Comparison between the total number of thousands of contacted fragments in PCHi-C data and in simulated data, for human samples. Each dot represents one sample. **B.** Comparison between the total genomic length covered by contacted fragments in PCHi-C data and in simulated data, for human samples. Each dot represents one sample. Genomic lengths are given in megabases (Mb). **C.** Histogram of the number of contacted restriction fragments *per* bait, for human PCHi-C data (orange) and simulated data (blue). **D.** Histogram of the number of contacting baits *per* restriction fragment, for human PCHi-C data (orange) and simulated data (blue). **C-D.** All human samples are combined in a single dataset for this analysis.

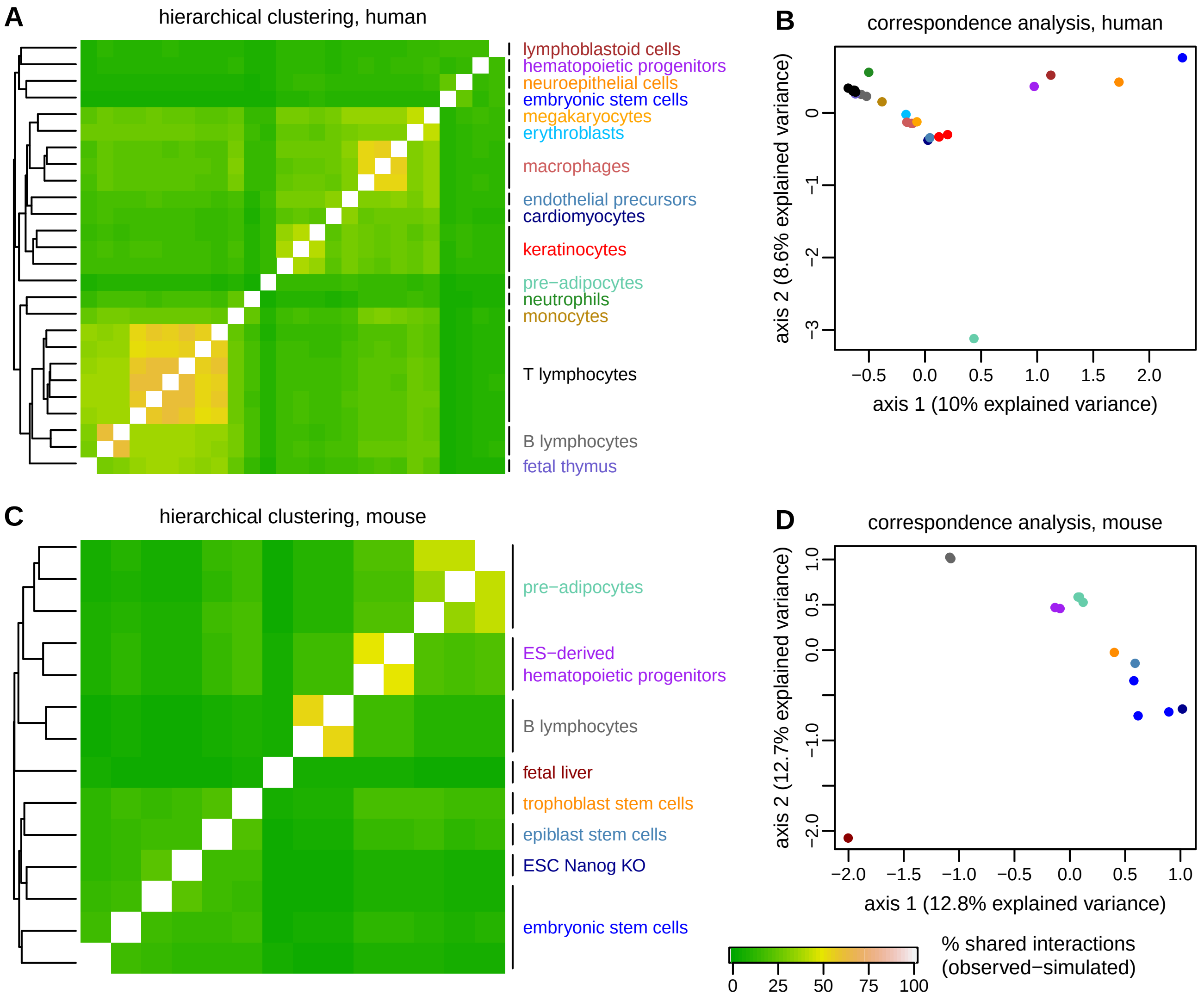


**Supplemental Fig. S3**: Chromatin contact maps cluster by cell type within species. **A.** For each pair of human samples, we display the difference between the percentage of shared interactions in PCHi-C data and the percentage of shared interactions in simulated data. The percentage of shared interactions is computed as 100 times the ratio between the number of interactions observed in both samples and the number of interactions observed in at least one of the two samples. We used subsampled interaction data for this analysis (Methods). The left panel displays a hierarchical clustering of samples based on this similarity measure. **B.** First factorial map of a correspondence analysis computed on human PCHi-C data. **C.** Same as **A**, for mouse. **D.** Same as **B**, for mouse.

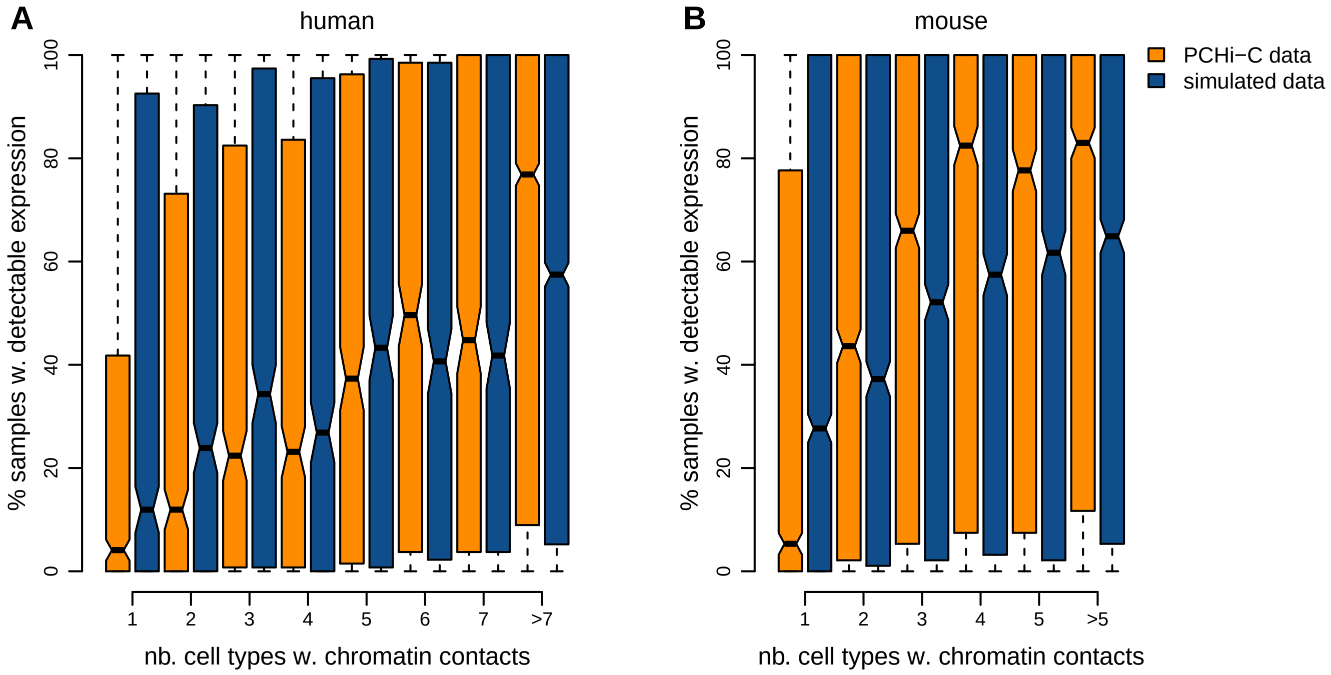


**Supplemental Fig. S4**: Gene expression breadth does not fully explain the presence of ubiquitous PCHi-C interactions. **A.** Distribution of the number of samples in which gene expression is observed (RPKM$\geq 1$), as a function of the maximum number of cell types in which promoter-enhancer interactions are observed, for human PCHi-C and simulated data. Gene expression data was taken from Cardoso-Moreira *et al.*, 2019. **B.** Same as **A**, for mouse.

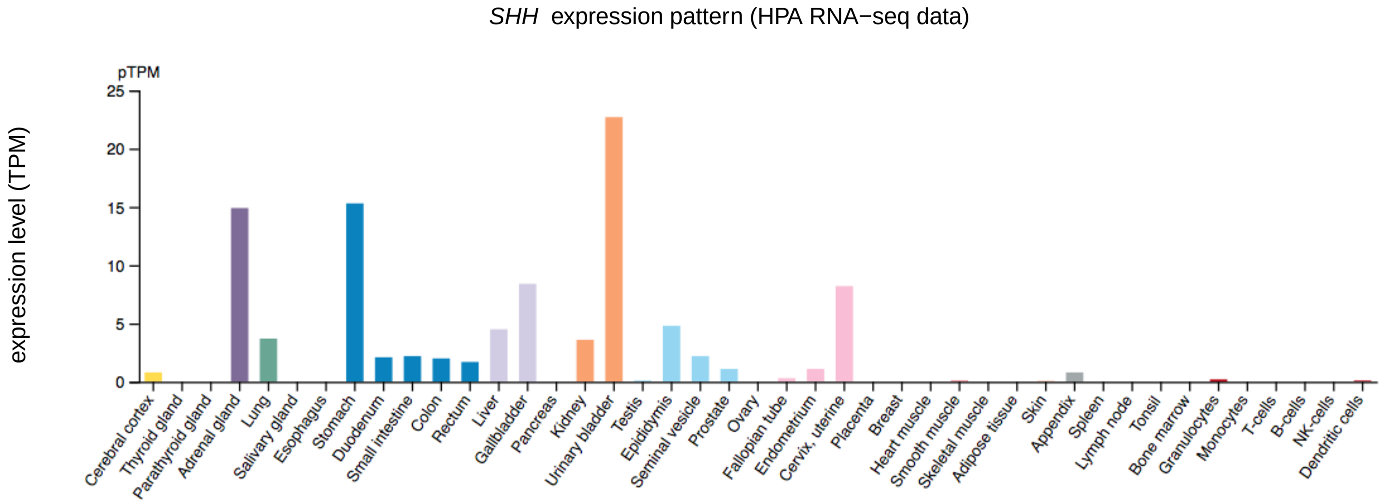


**Supplemental Fig. S5**: Gene expression pattern for the human *SHH* gene. The image was retrieved from the Human Protein Atlas (`www.proteinatlas.org`). The Y axis represents transcript *per* million (TPM) expression levels, compute on the Human Protein Atlas tissue RNA-seq collection (Uhlén *et al.*, 2015). The X axis represents different tissues and cell types, including immune cells sampled in our PCHi-C dataset. There is little or no detectable *SHH* expression in monocytes, T-cells and B-cells.

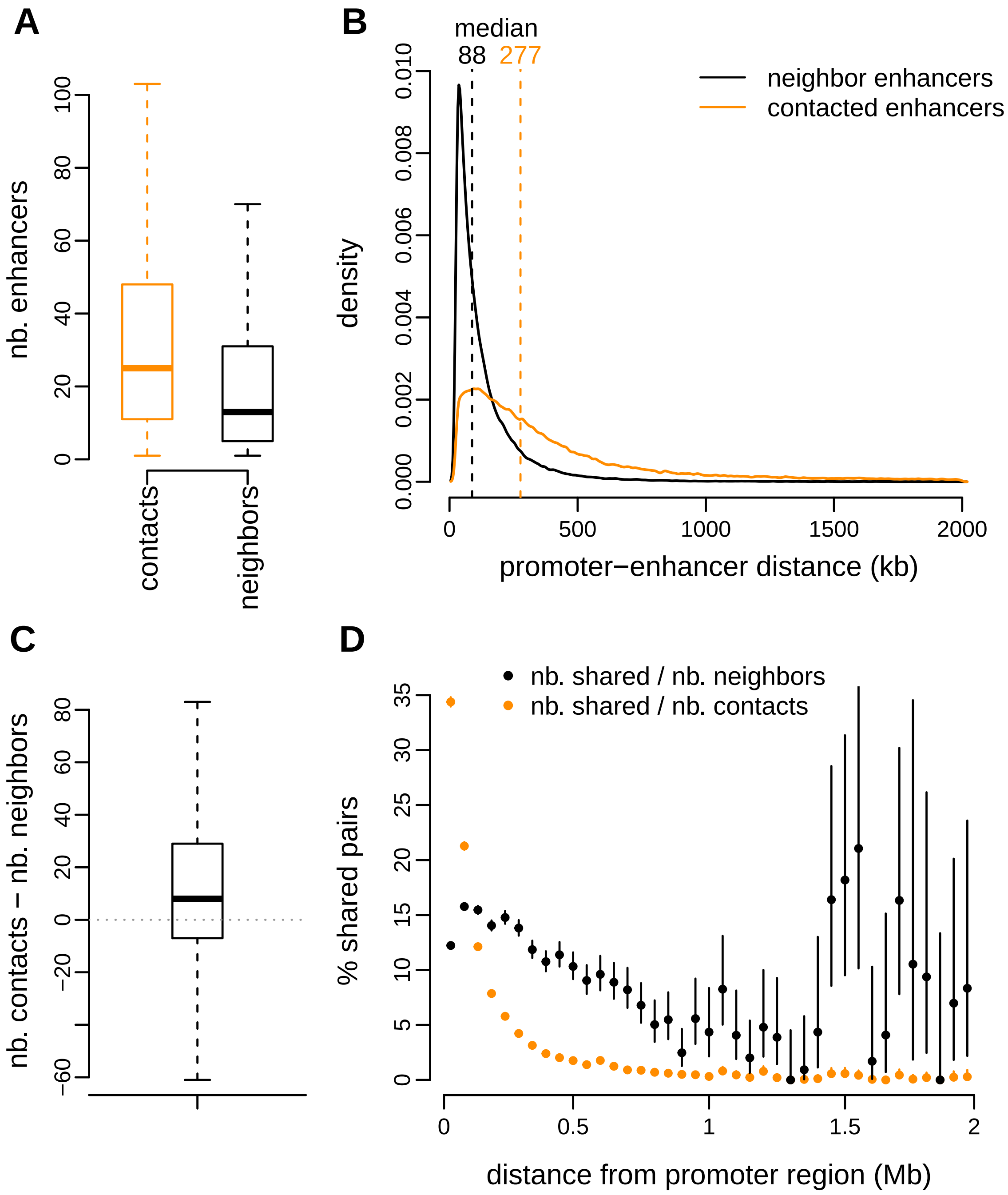


**Supplemental Fig. S6**: Comparisons between gene-enhancer regulatory pairs inferred with PCHi-C data (orange) and with the genomic proximity approach (black), for **human** data. **A.** Boxplots representing the distribution of the number of enhancers attributed to each gene with PCHi-C data or with the genomic proximity approach. **B.** Density plot representing the distribution of distances between transcription start sites and enhancers, for both approaches. **C.** Boxplot representing the distribution of the difference in the number of enhancers attributed to each gene with the PCHi-C approach and with the genomic proximity approach. **D.** Percentage of gene-enhancer pairs shared between the two approaches, as a function of the genomic distance between promoters and enhancers.

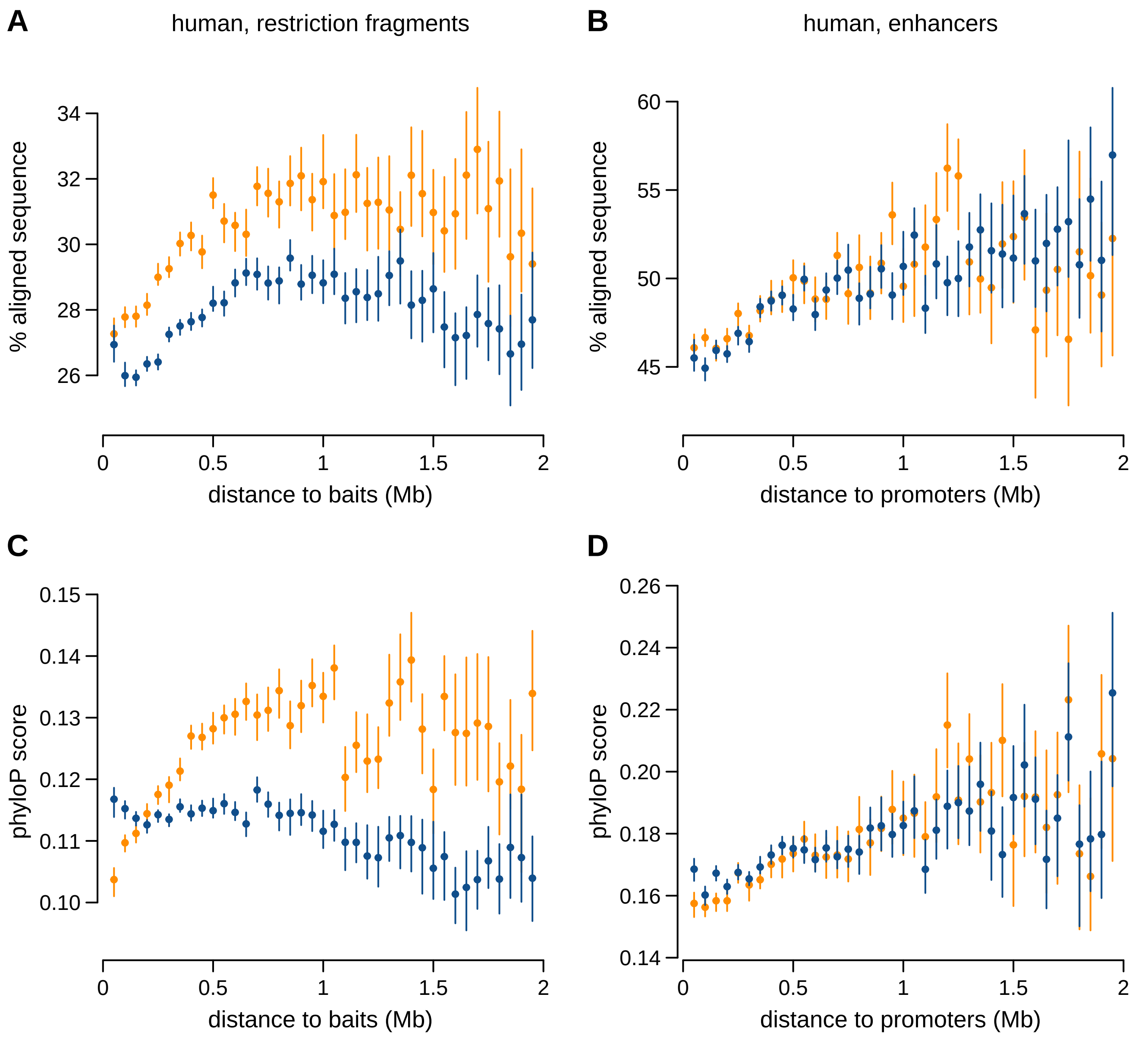


**Supplemental Fig. S7**: Sequence conservation measured by our pairwise alignment score or by the phyloP score, for **human** restriction fragments and enhancers. **A.** Average pairwise alignment scores (percentage of aligned sequence, human *vs.* mouse) of contacted restriction fragments in PCHi-C data (orange) and simulated data (blue), as a function of the median genomic distance between restriction fragments and contacting baits. **B.** Same as **B.**, for ENCODE enhancers. **C)** Average phyloP score of contacted restriction fragments in PCHi-C data (orange) and simulated data (blue), as a function of the median genomic distance between restriction fragments and contacting baits. **D)** Same as **B.**, for ENCODE enhancers. **A-D)** Dots represent mean values, vertical segments represent 95% confidence intervals of the mean, obtained with a non-parametric bootstrap approach (Methods). Exonic sequences are masked for these computations.

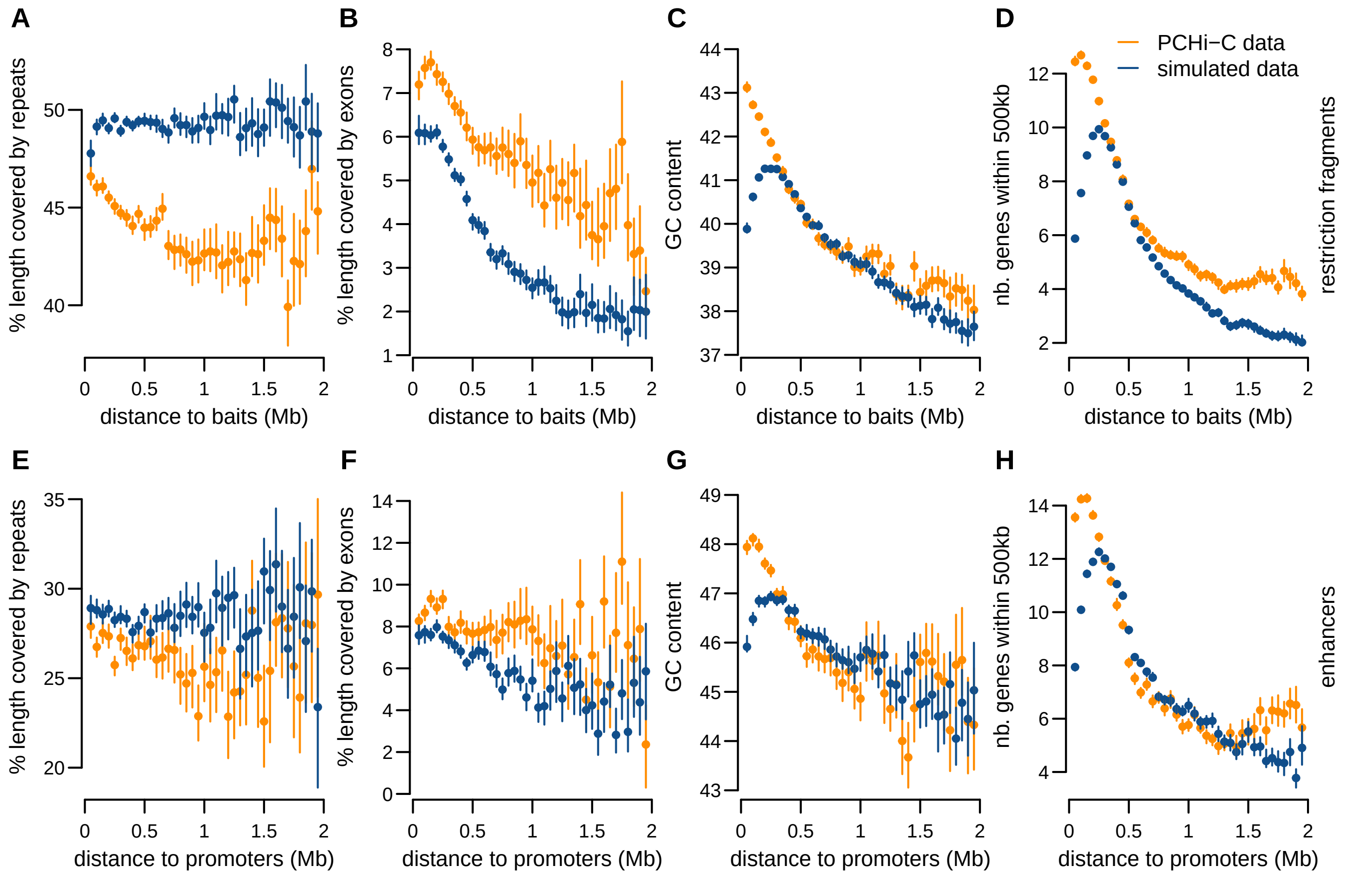


**Supplemental Fig. S8**: Genomic characteristics of restriction fragments and enhancers contacted by baited promoters, in **human** PCHi-C data and simulated data. From left to right: Average percentage of length covered by repeated elements, average percentage of length covered by exons, GC content, number of genes found within a maximum distance of 500 kb. **A-D)** Genomic characteristics of restriction fragments, for PCHi-C data (orange) and simulated data (blue). **E-H)** Genomic characteristics of ENCODE enhancers. **A-H)** Dots represent average values across all elements in a given distance class; vertical segments represent 95% confidence intervals, computed with a non-parametric bootstrap approach (Methods).

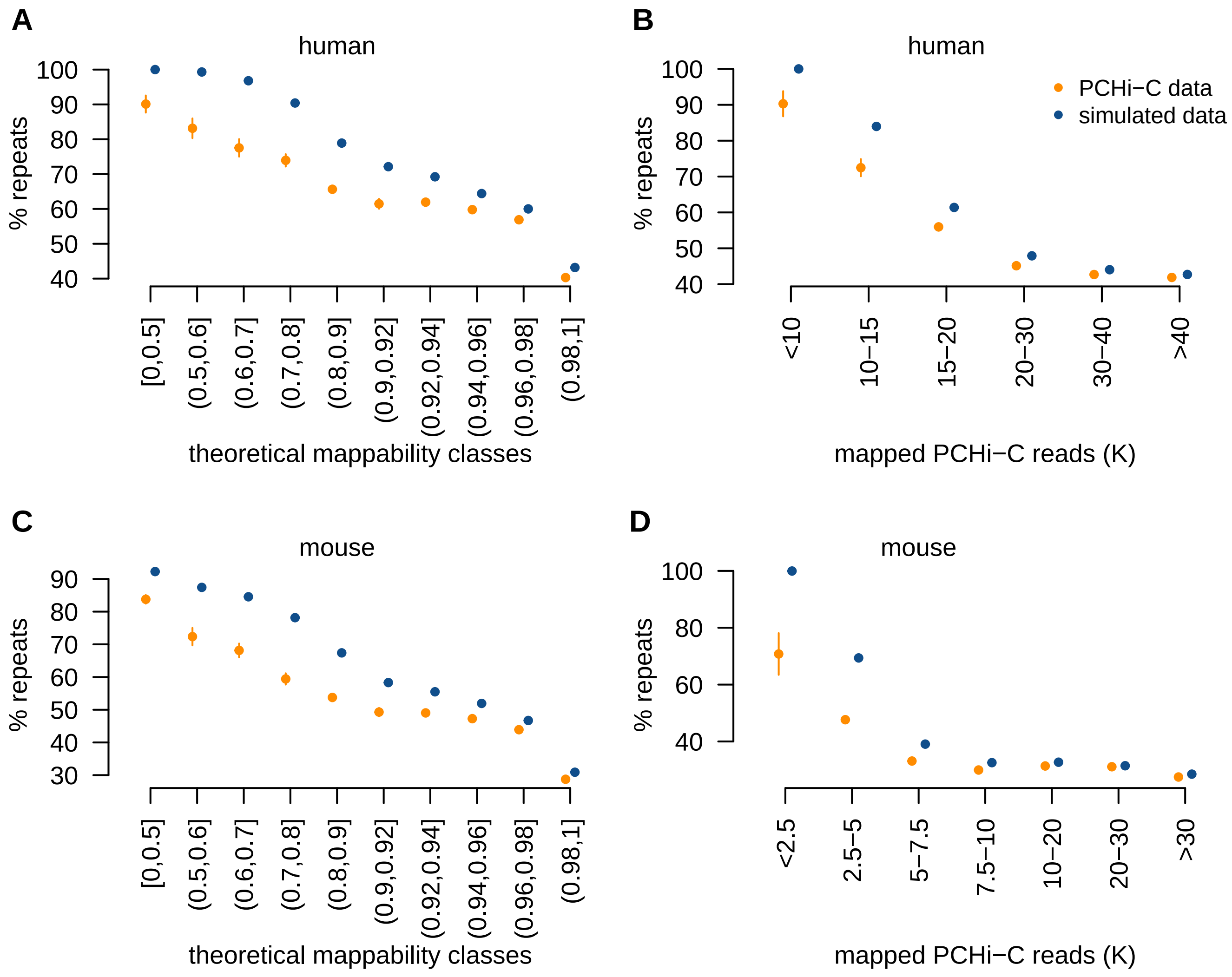


**Supplemental Fig. S9**: Restriction fragments that are contacted in the PCHi-C data contain lower proportions of repeated elements than those included in the simulated data, irrespective of their theoretical mappability and PCHi-C read count. **A.** Proportion of repeated elements as a function of classes of theoretical mappability of contacted restriction fragments in **human** PCHi-C data (orange) and simulated data (blue). Mappability classes are determined based on the mappable length fraction (number of mappable bases/total length). **B.** Proportion of repeated elements as a function of the total number of thousands of mapped reads, combined across human PCHi-C samples. **C)** Same as **A.**, for **mouse**. **D)** Same as **B.**, for **mouse**. **A-D)** Dots represent median values across restriction fragments in a given mappability class; vertical segments represent 95% confidence intervals.

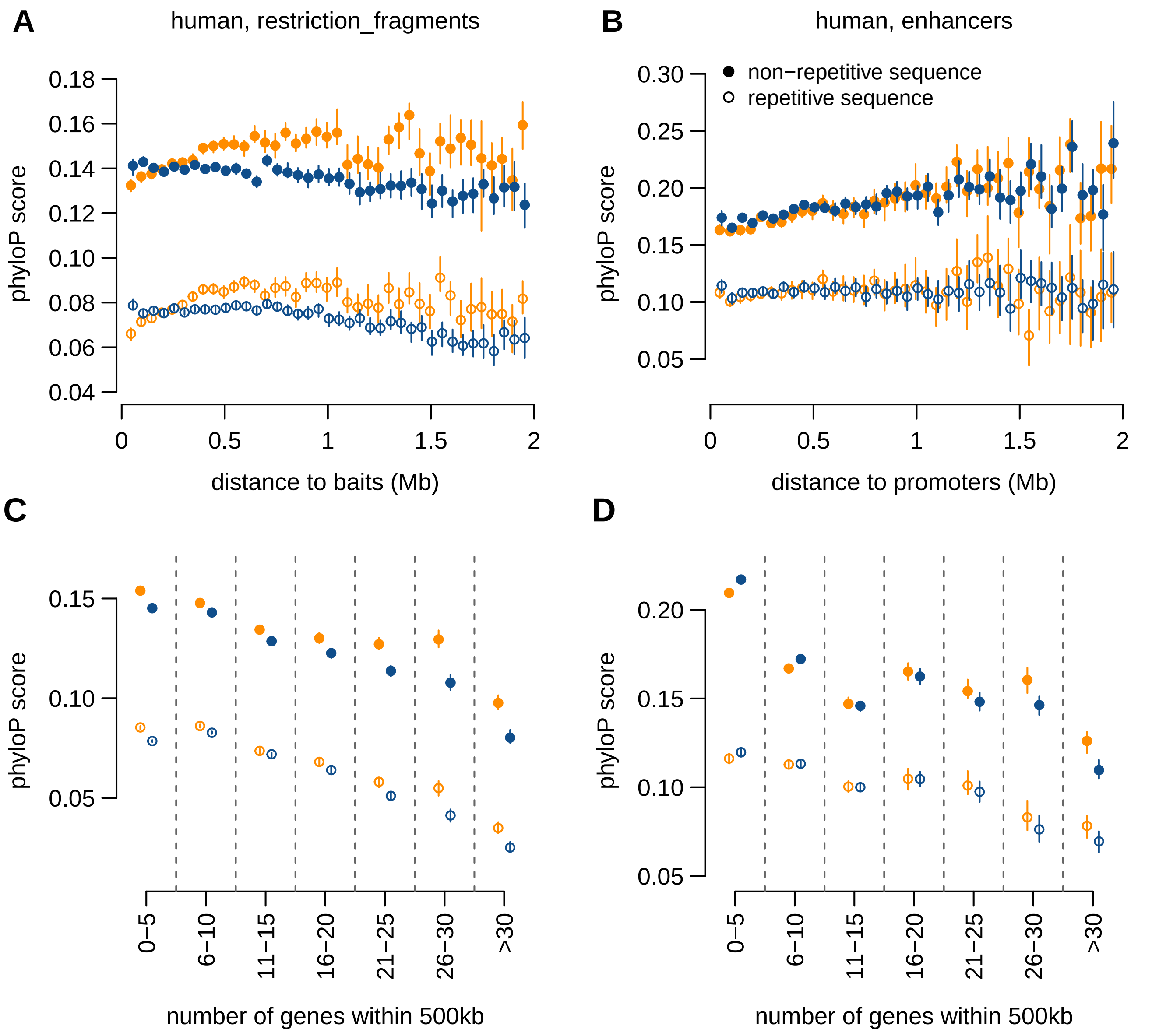


**Supplemental Fig. S10**: Sequence conservation measured by phyloP score for **human** restriction fragments and enhancers. **A.** Average phyloP score of contacted restriction fragments in PCHi-C data (orange) and simulated data (blue), as a function of the median genomic distance between restriction fragments and contacting baits. **B.** Same as **A.**, for ENCODE enhancers. **C)** Average phyloP score of contacted restriction fragments, as a function of the number of genes found within a maximum distance of 500 kb from the restriction fragment. **D)** Same as **C)**, for ENCODE enhancers. **A-D)** Filled dots represent non-repetitive part of sequences; empty dots represent the repetitive part. Dots represent mean values, vertical segments represent 95% confidence intervals of the mean, obtained with a non-parametric bootstrap approach (Methods).

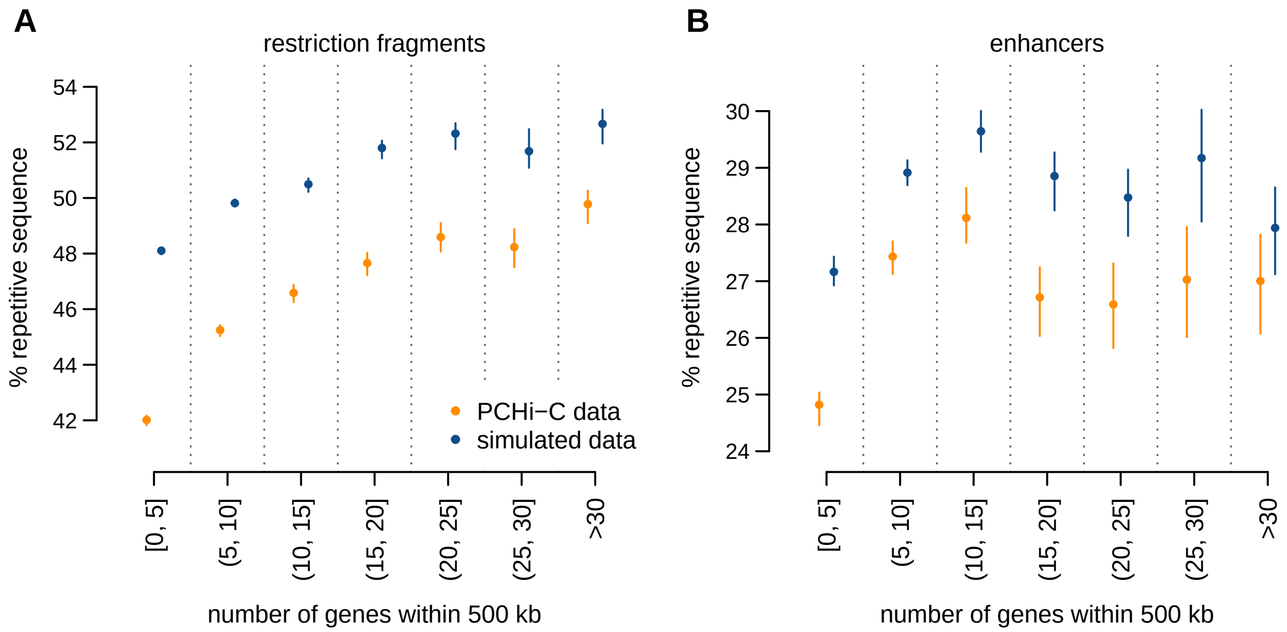


**Supplemental Fig. S11**: Gene density in the neighboring regions is correlated with repetitive sequence content for **human** contacted restriction fragments and enhancers. **A.** Average fraction of restriction fragment length that is covered by repetitive sequences, as a function of the number of genes found at most 500kb away from the restriction fragment. **B.** Same as **A.**, for ENCODE enhancers. **A.,B.** Dots represent average values across all elements in a class; vertical segments represent 95% confidence intervals, computed with a non-parametric bootstrap approach (Methods).

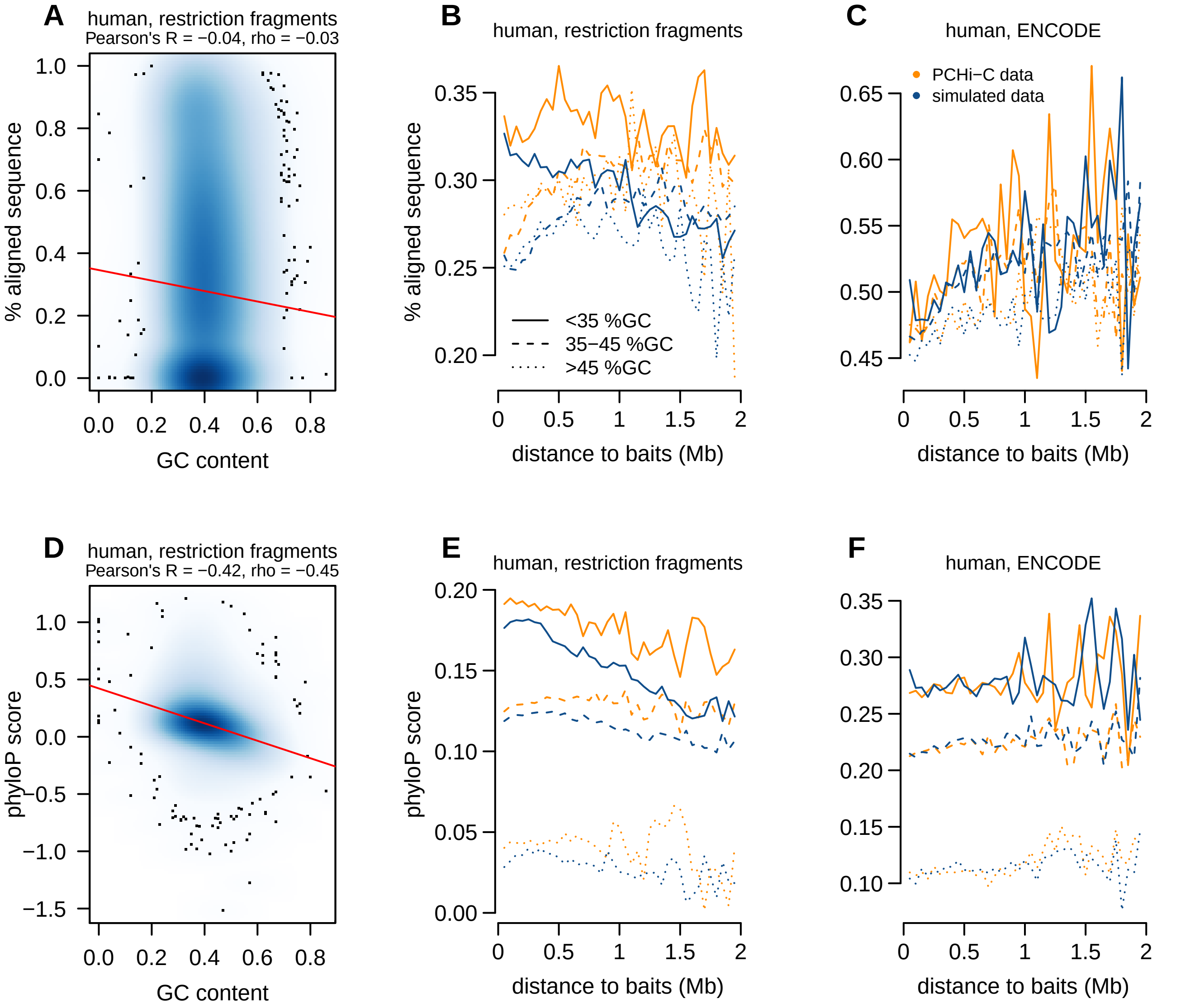


**Supplemental Fig. S12**: Sequence conservation and GC content. **A.** Relationship between the average sequence conservation levels (percentage aligned sequence) and the sequence GC content, for human restriction fragments. The red line represents the linear regression of the two variables. **B.** Average sequence conservation levels by classes of GC content of human restriction fragments contacted in PCHi-C (orange) and in simulated (blue) data, as a function of the median genomic distance between restriction fragments and contacting baits. **C)** Same as **B.**, for contacted ENCODE enhancers. **D-F)** Same as **A-C)**, for sequence conservation measured with the phyloP score.

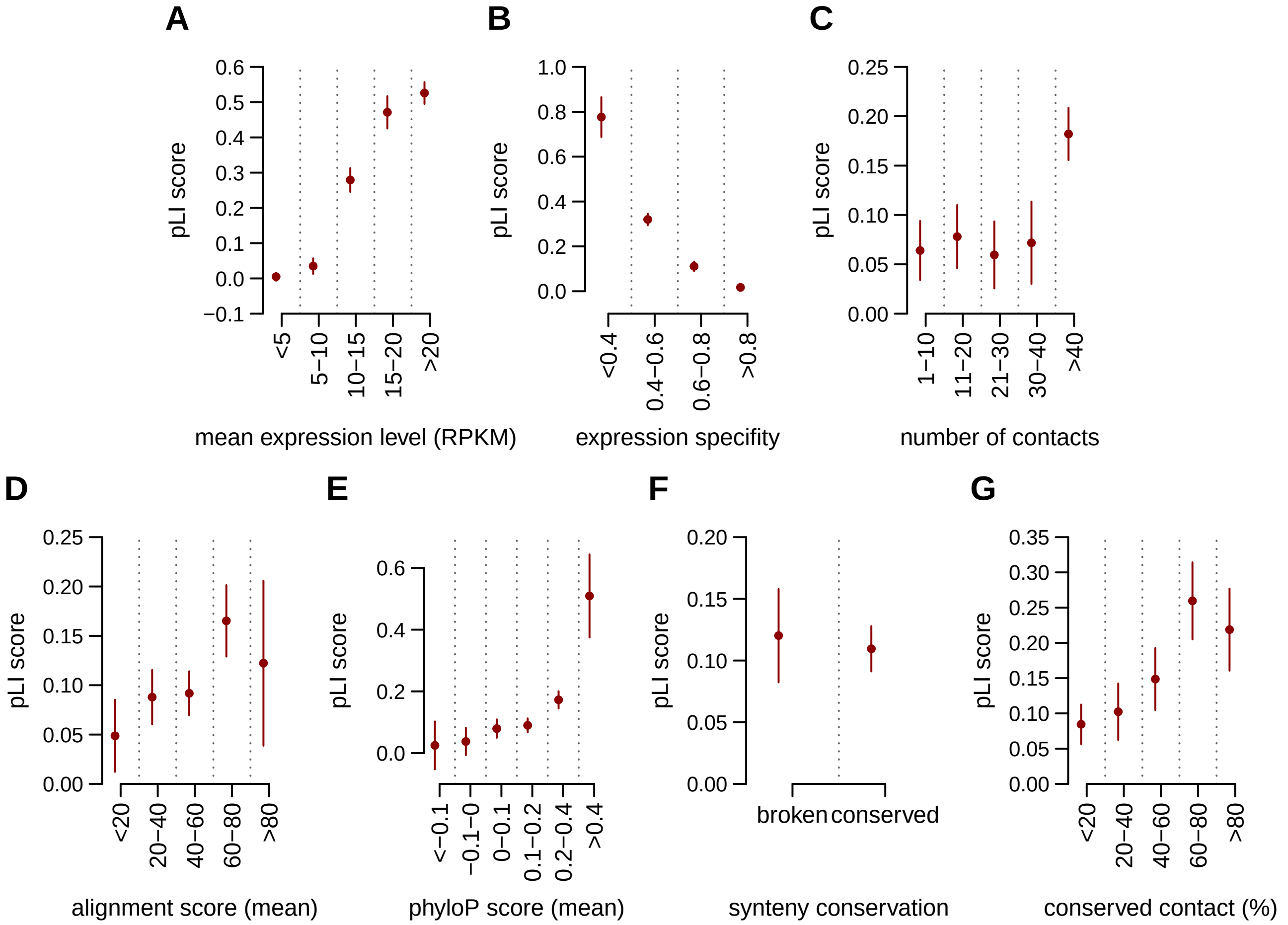


**Supplemental Fig. S13**: The constraint on **human** gene sequence is associated with gene expression, and the evolution of *cis*-regulatory landscapes. Gene constraint is measured by the probability of intolerance to loss-of-function mutations (pLI score, data from patterns of variation in human exome data Lek et al. 2016, Methods). Enhancer predictions are taken from ENCODE. **A.** pLI score as a function of the average human expression levels (data from Cardoso-Moreira et al., 2019). **B.** pLI score as a function of the human gene expression specificity index (reaches values close to 0 for housekeeping genes, close to 1 for highly specific genes (Methods)). **C.** pLI score as a function of the number of contacted enhancers in human PCHi-C data. **D.** pLI score as a function of the average sequence conservation of human contacted enhancers. **E.** pLI score as a function of the average phyloP score of human contacted enhancers. **F.** pLI score depending on whether or not genes underwent least one break of synteny with the contacted enhancers between human and mouse genomes. **F.** pLI score as a function of the proportion of human promoter-enhancers contacts conserved in mouse PCHi-C data. **A-G.** Dots represent median values across all genes in a class, vertical segments represents 95% confidence intervals for the median.

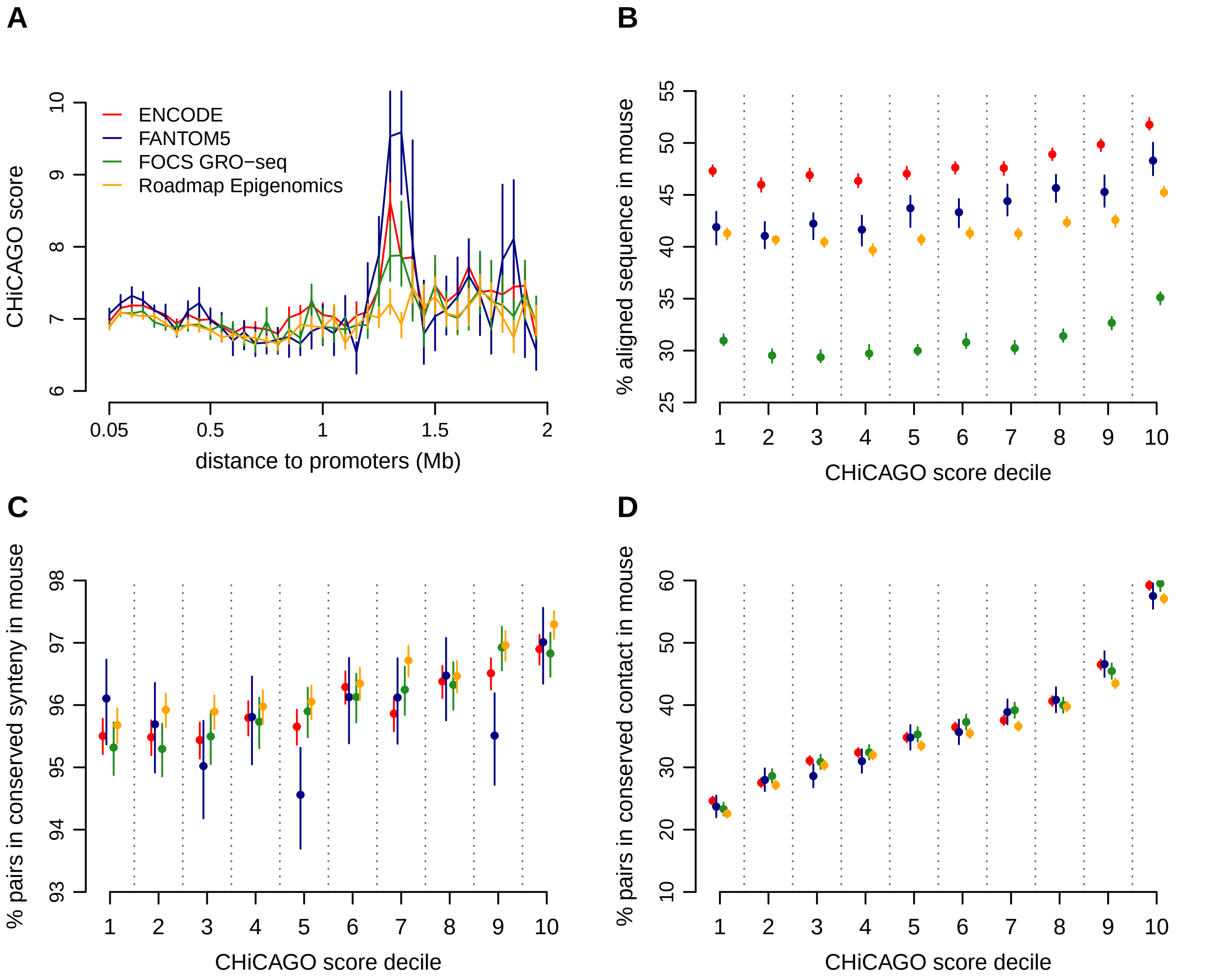


**Supplemental Fig. S14**: The CHiCAGO score, which measures the statistical significance of promoter-chromatin contacts, is correlated with the conservation of **human** *cis*-regulatory landscapes. **A.** Average CHiCAGO score as a function of the distance between promoters and predicted enhancers in the genome, for ENCODE (red) and FANTOM5 (blue), FOCS GRO-seq (green) and Roadmap Epigenomics enhancers (orange). **B.** Sequence conservation of contacted enhancers, calculated as the percentage of aligned nucleotides between human and mouse, as a function of the CHiCAGO score decile. **C)** Proportion of human promoter-enhancer pairs maintained in synteny in the mouse genome, as a function of the CHiCAGO score decile. **D)** Proportion of human promoter-enhancer contacts that are conserved in mouse PCHi-C data, as a function of the CHiCAGO score decile.

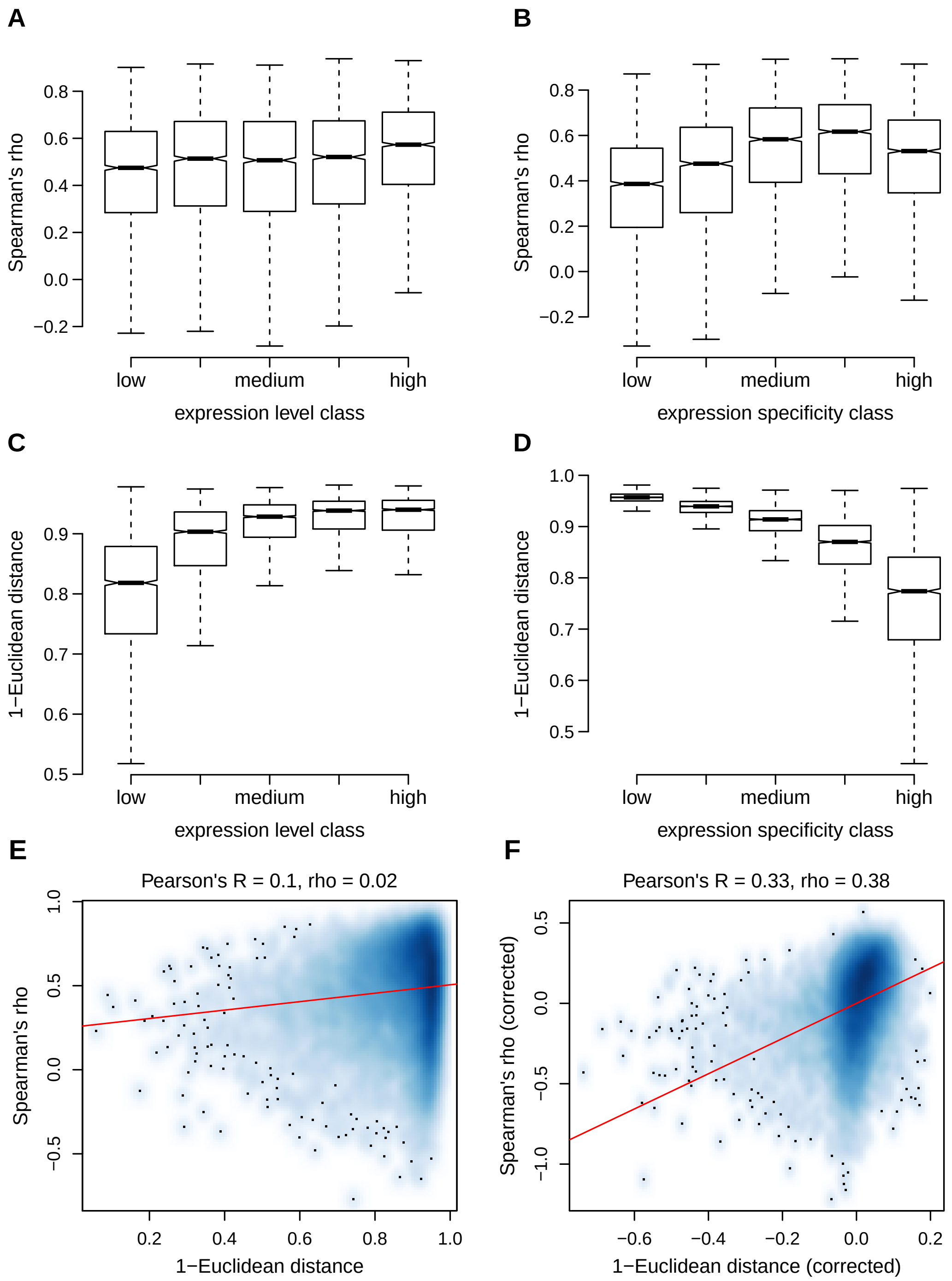


**Supplemental Fig. S15**: Gene expression level and gene expression specificity are correlated with the rate of gene expression profile evolution. **A.** Distribution of Spearman's correlation coefficients between human and mouse relative gene expression profiles, for different expression level classes. Genes were divided into 5 equal-sized classes depending on their average expression levels across mouse and human samples. **B.** Distribution of Spearman's correlation coefficients between human and mouse relative gene expression profiles, for different expression specificity classes. Genes were divided into 5 equal-sized classes depending on their expression specificity (Methods), averaged across human and mouse. **C.** Euclidean similarity score (1-Euclidean distance) of gene expression profile evolution according to gene expression level class. **D.** Euclidean similarity score of gene expression profile evolution according to gene expression specificity class. **E.** Relationship between Spearman's correlation coefficients and Euclidean similarity scores. The red line represents the linear regression of the two variables. **F.** Relationship between Spearman's correlation coefficients and Euclidean similarity scores, after correcting for the effect of the gene expression level and gene expression specificity with multiple regression models (Methods). **A-D.** Boxplots exclude outlier points; notches represent 95% confidence intervals for the median values.

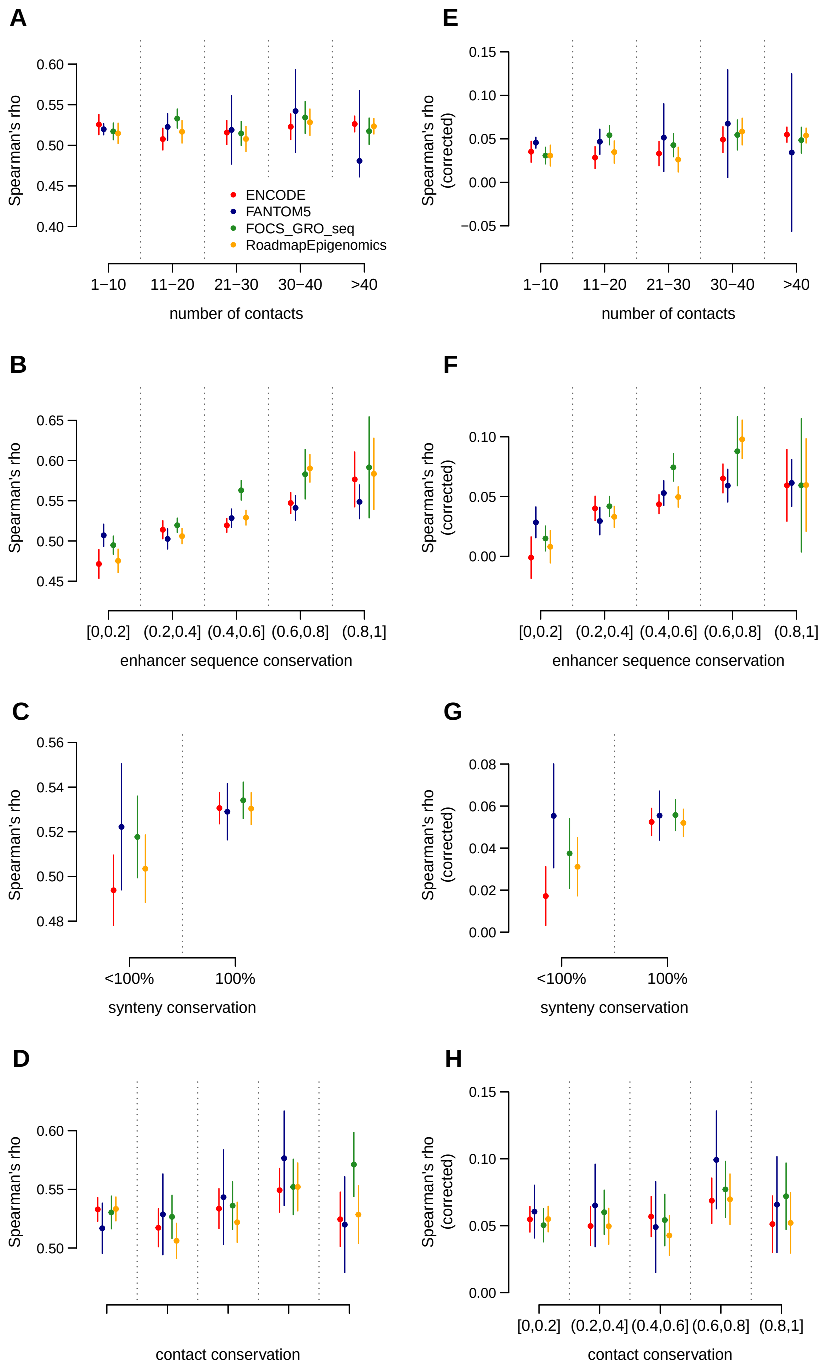


**Supplemental Fig. S16**: Human *cis*-regulatory landscape evolution and gene expression profile evolution. We measure gene expression conservation based on Spearman's correlation between relative expression profiles across organs and developmental stages, for each pair of orthologous genes. We analyze the relationship between expression similarity and regulatory landscape characteristics, evaluated using enhancers from ENCODE (red), FANTOM5 (blue), FOCS GRO-seq (green) and Roadmap Epigenomics (orange). Left panels represent the raw expression similarity data; right panels represent the expression similarity measure after correcting for the effect of gene expression levels and gene expression specificity with a multiple linear regression model. **A,E)** Average expression similarity as a function of the number of human contacted enhancers in PCHi-C. **B,F)** Average expression similarity as a function of the average sequence conservation of human contacted enhancers. **C,G)** Average expression similarity for genes that have undergone no breaks of synteny (left) and at least one break of synteny (right) with the homologous enhancers between human and mouse genomes. **D,H)** Average expression similarity as a function of the proportion of human promoter-enhancers contacts conserved in mouse PCHi-C data.

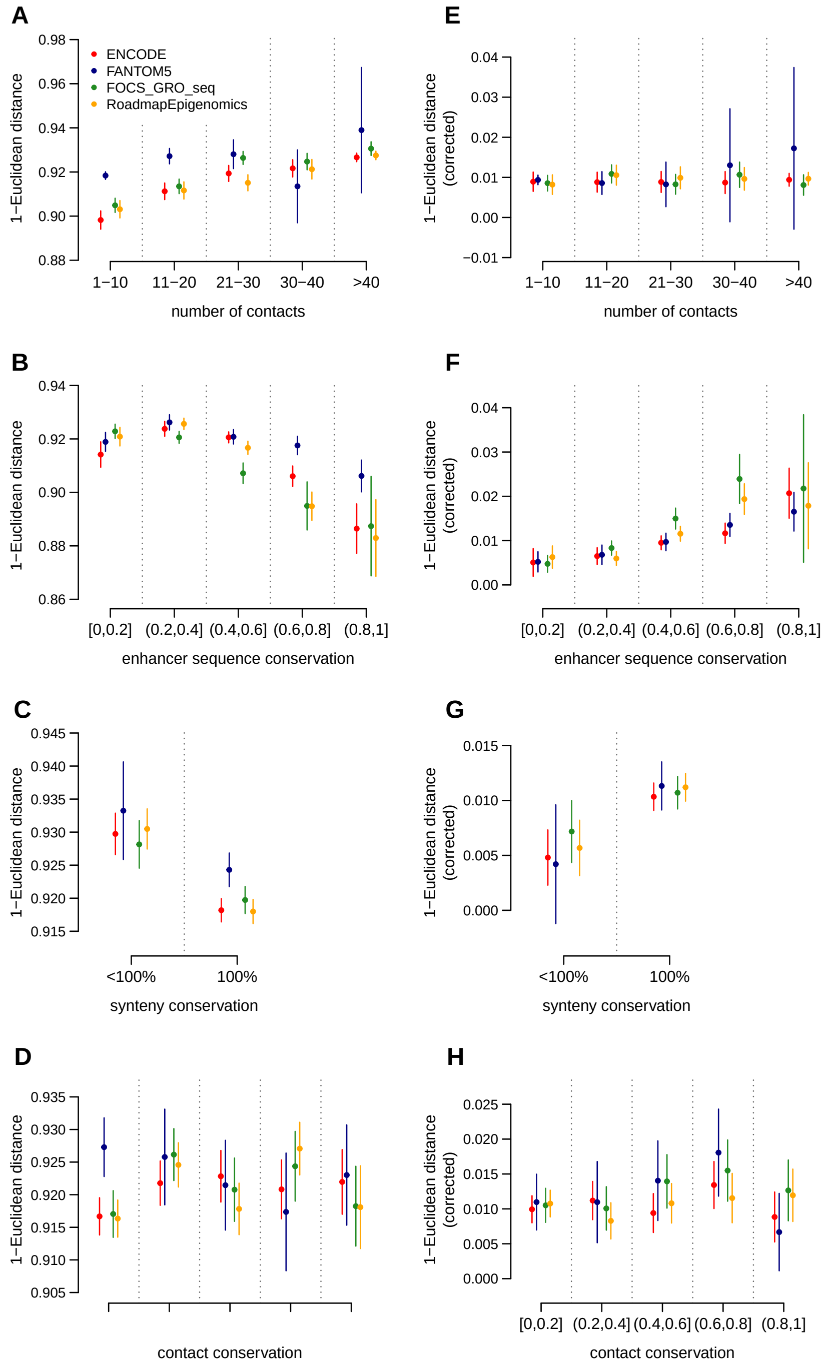


**Supplemental Fig. S17**: Human *cis*-regulatory landscape evolution and gene expression profile evolution. We measure gene expression conservation based on the Euclidean distance between relative expression profiles across organs and developmental stages, for each pair of orthologous genes. The Euclidean distance is transformed into a similarity measure (1-Euclidean distance). We analyze the relationship between expression similarity and regulatory landscape characteristics, evaluated using enhancers from ENCODE (red), FANTOM5 (blue), FOCS GRO-seq (green) and Roadmap Epigenomics (orange). Left panels represent the raw expression similarity data; right panels represent the expression similarity measure after correcting for the effect of gene expression levels and gene expression specificity with a multiple linear regression model. **A,E)** Average expression similarity as a function of the number of human contacted enhancers in PCHi-C. **B,F)** Average expression similarity as a function of the average sequence conservation of human contacted enhancers. **C,G)** Average expression similarity for genes that have undergone no breaks of synteny (left) and at least one break of synteny (right) with the homologous enhancers between human and mouse genomes. **D,H)** Average expression similarity as a function of the proportion of human promoter-enhancers contacts conserved in mouse PCHi-C data.

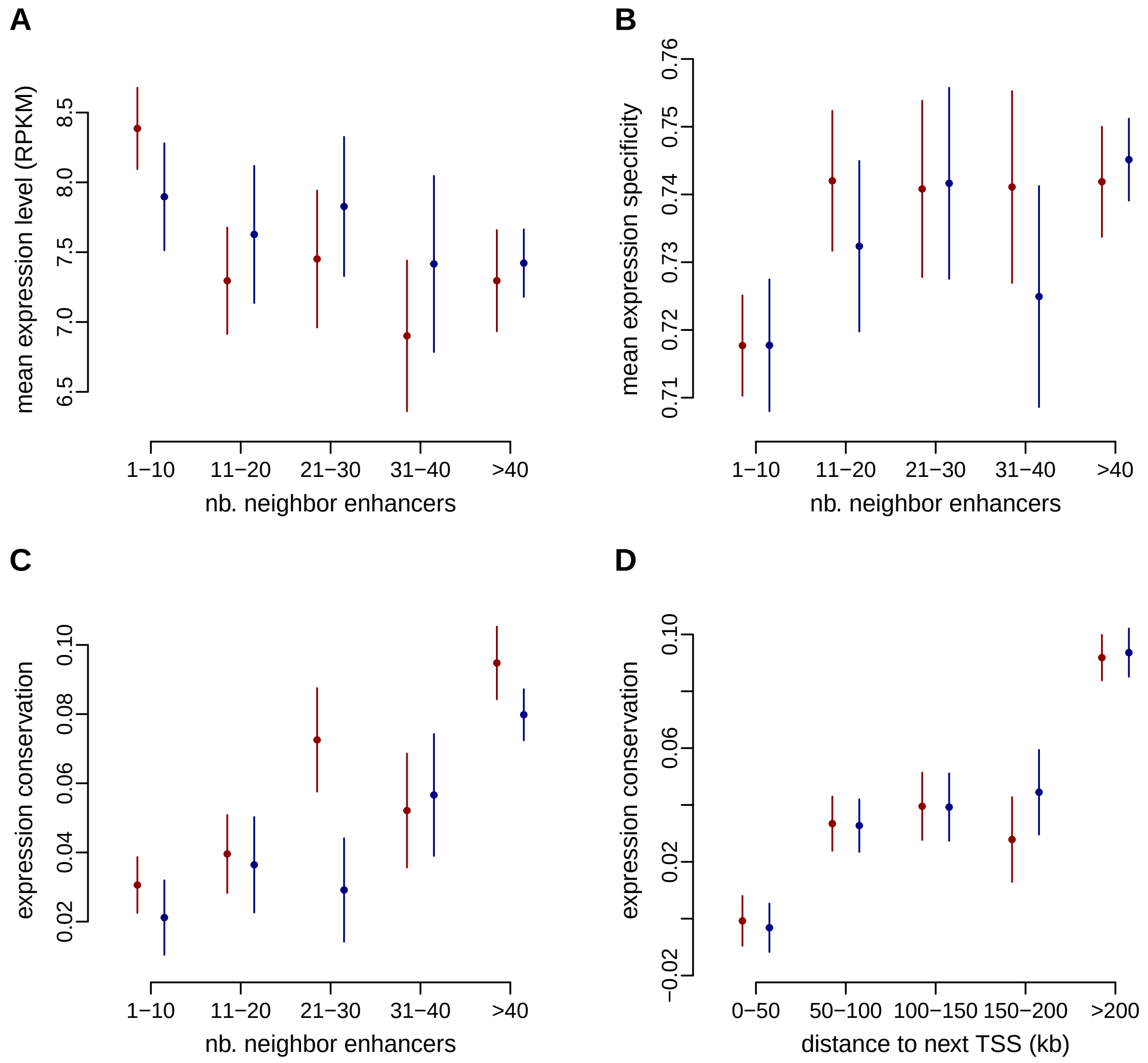


**Supplemental Fig. S18**: Relationship between gene expression characteristics and the number of enhancers attributed to each gene with the genomic proximity approach (neighbor enhancers below), for human (red) and mouse (blue). **A.** Average expression levels as a function of the number of neighbor enhancers. **B.** Gene expression specificity index (broadly expressed genes have values close to 0, genes expressed in few samples have values close to 1; Methods) as a function of the number of neighbor enhancers. **C)** Expression conservation as a function of the number of neighbor enhancers. **D)** Expression conservation as a function of the distance to next TSS . **A-D)** Dots represent median values across all genes in a class, vertical segments represent 95% confidence intervals for the median. **C,D)** Gene expression conservation is measured with Spearman's correlation coefficient between human and mouse relative expression profiles, for pairs of 1-to-1 orthologous genes, across organs and developmental stages (expression data from Cardoso-Moreira et al., 2019). Expression conservation is further corrected to account for the effect of expression levels and of expression specificity with a multiple linear regression model (Methods). Expression conservation values are the same for both species, but the numbers of enhancers attributed to each gene may differ.

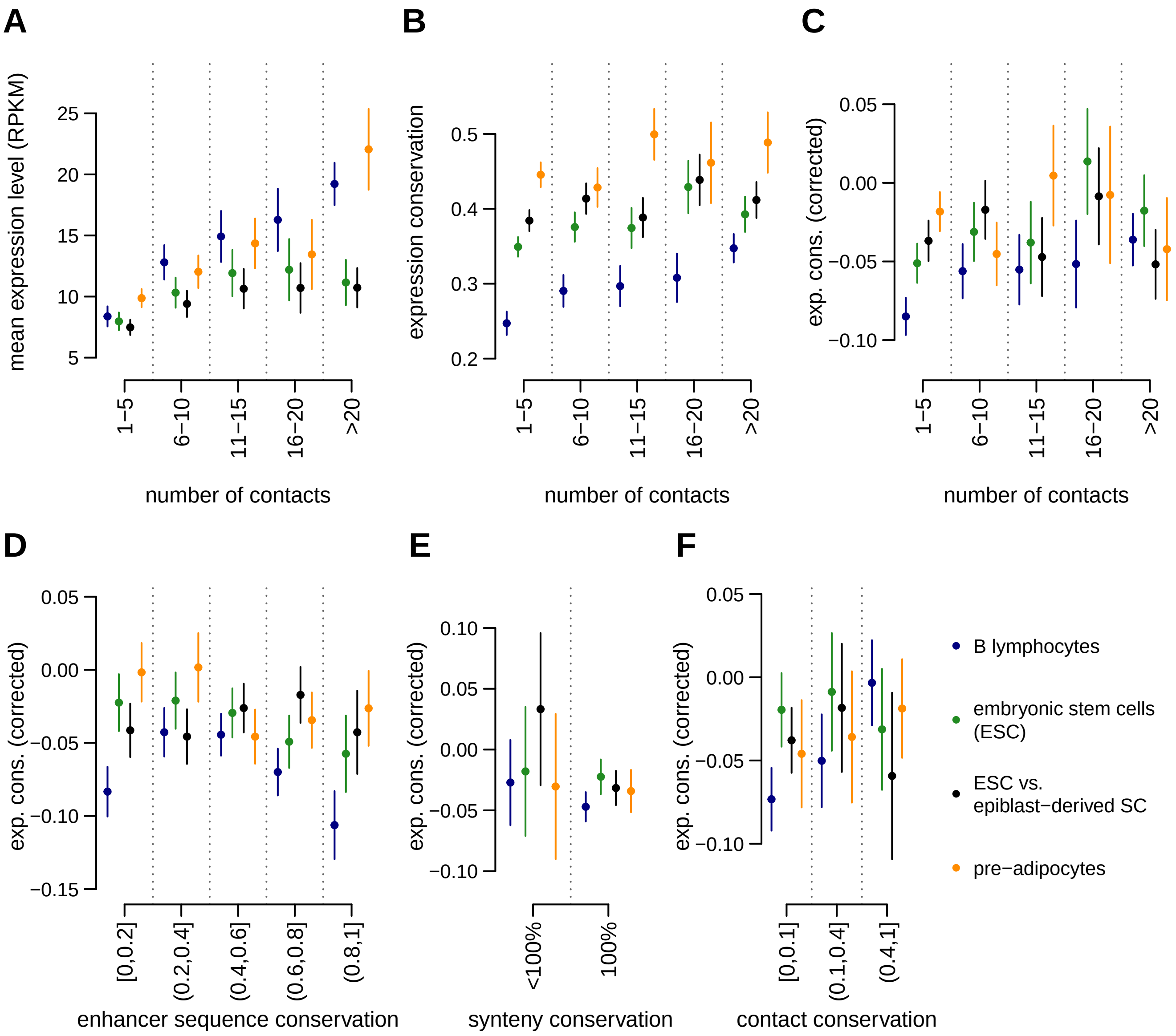


**Supplemental Fig. S19**: The complexity of *cis*-regulatory landscapes is positively correlated with gene expression levels in individual cell types. **A.** Gene expression levels in different cell types, as a function of the number of contacted enhancers in the corresponding PCHi-C samples. We show the average expression level for human and mouse, for the two cell types being compared. **B.** Gene expression level conservation between human and mouse in similar cell types, as a function of the number of contacted enhancers in the corresponding human PCHi-C samples. **C)** Gene expression level conservation between human and mouse in similar cell types, after correcting for the effect of the expression level, as a function of the number of contacted enhancers in human PCHi-C data. **D)** Gene expression level conservation, as a function of the average sequence conservation of human contacted ENCODE enhancers. **E)** Gene expression level conservation for genes that have undergone no breaks of synteny (left) or at least one (right) with the homologous ENCODE enhancers between human and mouse genome. **F)** Gene expression level conservation, as a function of the proportion of human promoter-enhancers contacts conserved in mouse PCHi-C. **A-F)** Dots represent median values across all genes in a class, vertical segments represents 95% confidence intervals for the median.

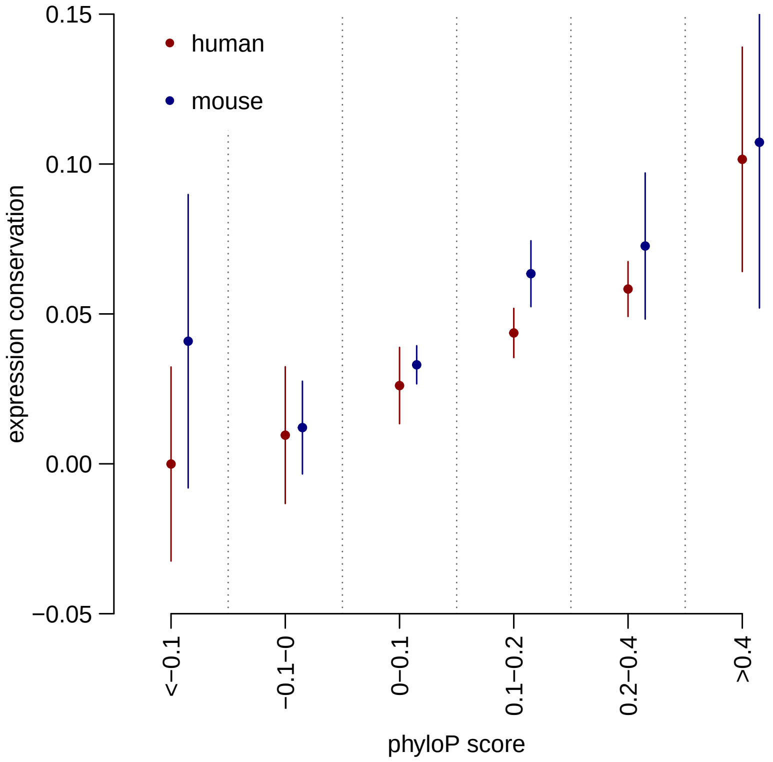


**Supplemental Fig. S20**: Average expression conservation as a function of the average sequence conservation of contacted ENCODE enhancers, measure by phyloP score, in PCHi-C data, for human (red) and mouse (blue). Dots represent median values across all genes in a class, vertical segments represents 95% confidence intervals for the median. Gene expression conservation is measured with Spearman's correlation coefficient between human and mouse relative expression profiles, for pairs of 1-to-1 orthologous genes, across organs and developmental stages (expression data from Cardoso-Moreira et al., 2019). Expression conservation is further corrected to account for the effect of expression levels and of expression specificity with a multiple linear regression model (Methods). Expression conservation values are the same for both species, but PCHi-C contact maps differ.

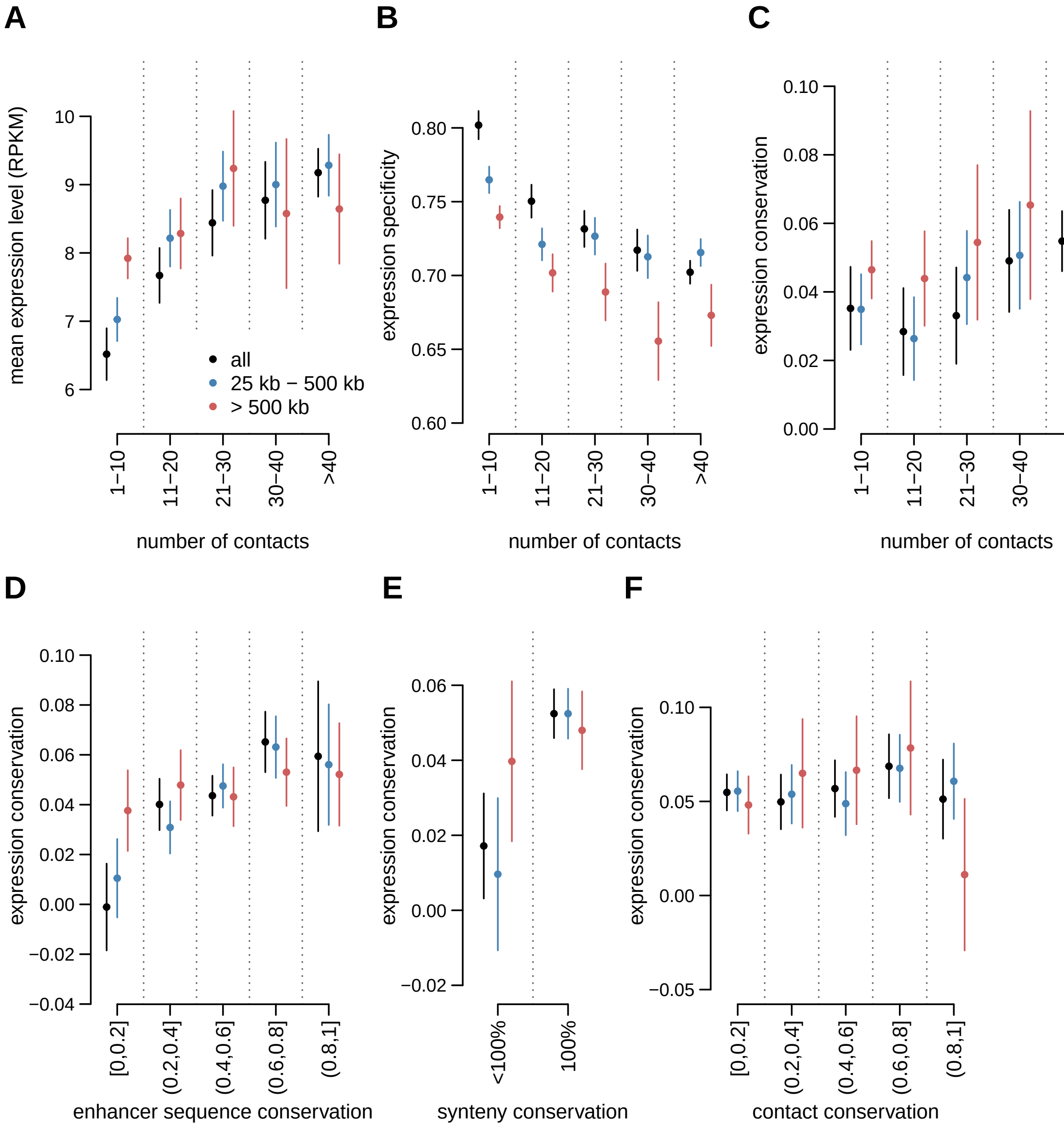


**Supplemental Fig. S21**: *Cis*-regulatory landscapes co-evolve with gene expression profiles. Gene expression conservation is measured with Spearman's correlation coefficient between human and mouse relative expression profiles, for pairs of 1-to-1 orthologous genes, across organs and developmental stages (expression data from Cardoso-Moreira et al., 2019). Expression conservation is further corrected to account for the effect of expression levels and of expression specificity with a multiple linear regression model (Methods). Enhancer predictions are taken from ENCODE. **A.** Average expression levels as a function of the number of contacted enhancers in PCHi-C data. Promoter-enhancer contacts are divided into two classes according to the distance between them: medium range (25 kb-500 kb). in blue, long range (above 500 Kb). in red. The black dots represent the full set of promoter-enhancer contacts. **B.** Human gene expression specificity as a function of the number of contacted enhancers. **C)** Expression conservation as a function of the number of contacted enhancers in human PCHi-C data. **D)** Expression conservation as a function of the average sequence conservation of human contacted ENCODE enhancers. **E)** Expression conservation, depending on whether or not genes underwent at least one break of synteny with the contacted enhancers between human and mouse genomes. **F)** Expression conservation as a function of the proportion of human promoter-enhancers contacts conserved in mouse PCHi-C data. **A-F)** Dots represent median values across all genes in a class, vertical segments represents 95% confidence intervals for the median.

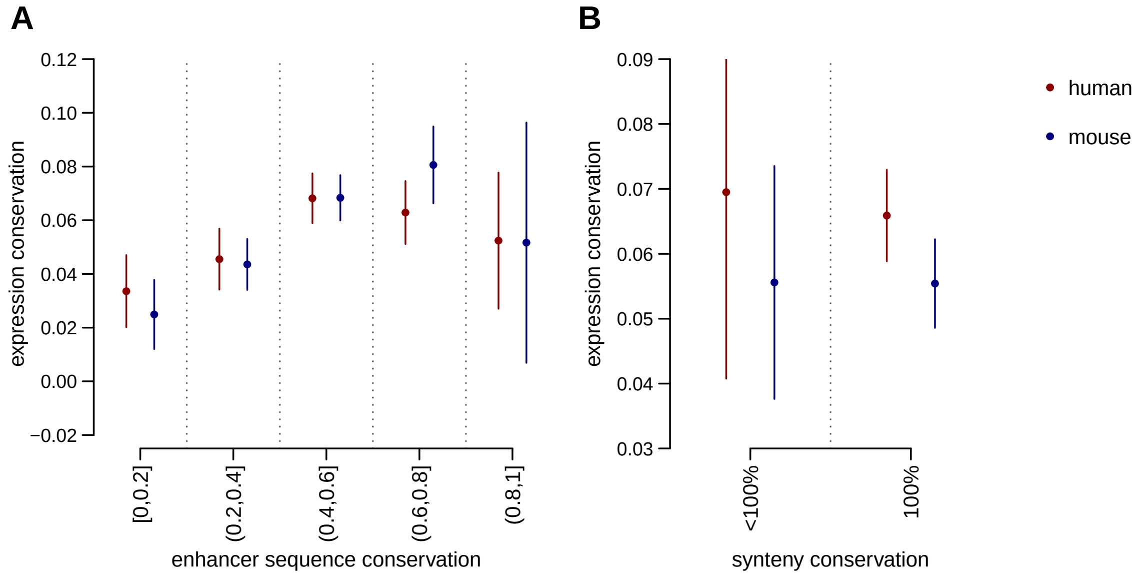


**Supplemental Fig. S22**: Gene expression conservation and evolution of neighbor enhancers, for human (red) and mouse (blue). **A.** Expression conservation as a function of the sequence conservation of neighbor enhancers. **B.** Expression conservation, depending on whether or not genes underwent at least one break of synteny with the neighbor enhancers between human and mouse genomes. **A-B.** Dots represent median values across all genes in a class, vertical segments represents 95% confidence intervals for the median. Enhancers are assigned to genes with the nearest-gene approach (Method).
